# An RNAi screen for genes that affect nuclear morphology in *Caenorhabditis elegans* reveals the involvement of unexpected processes

**DOI:** 10.1101/2021.06.29.450421

**Authors:** Richa Maheshwari, Mohammad M. Rahman, Daphna Joseph-Strauss, Orna Cohen-Fix

## Abstract

Aberration in nuclear morphology is one of the hallmarks of cellular transformation. However, the processes that, when mis-regulated, result aberrant nuclear morphology are poorly understood. In this study we carried out a systematic, high-throughput RNAi screen for genes that affect nuclear morphology in *Caenorhabditis elegans* embryos. The screen employed over 1700 RNAi constructs against genes required for embryonic viability. Nuclei of early embryos are typically spherical and their NPCs are evenly distributed. The screen was performed on early embryos expressing a fluorescently tagged component of the nuclear pore complex (NPC), allowing visualization of nuclear shape as well as the distribution of NPCs around the nuclear envelope. Our screen uncovered 182 genes whose down-regulation resulted in one or more abnormal nuclear phenotypes, including multiple nuclei, micronuclei, abnormal nuclear shape, anaphase bridges and abnormal NPC distribution. Many of these genes fall into common functional groups, including some that were not previously known to affect nuclear morphology, such as genes involved in mitochondrial function, the vacuolar ATPase and the CCT chaperonin complex. The results of this screen add to our growing knowledge of processes that affect nuclear morphology and that may be altered in cancer cells that exhibit abnormal nuclear shape.

## Introduction

The nucleus is bound by a nuclear envelope (NE) composed of two nuclear membranes, an inner nuclear membrane (INM) and an outer nuclear membrane (ONM), that are connected at sites of nuclear pore complexes (NPCs). The ONM is continuous with the endoplasmic reticulum (ER), while the INM interacts with the underlying lamina and chromatin-associated proteins (Cohen-Fix and Askjaer 2017; Ungricht and Kutay 2017; De Magistris and Antonin 2018). The lamina provides rigidity to the nucleus and helps in various nuclear processes including chromatin organization, DNA damage repair, and transcriptional regulation (Gruenbaum and Foisner 2015; De Leeuw *et al*. 2018; Dos Santos and Toseland 2021). The nucleus undergoes dramatic changes during the cell cycle. Most eukaryotic organisms follow an “open” mitosis scheme where the lamina and NPCs disassemble and the nuclear membranes becomes highly fenestrated, abolishing the permeability barrier that existed between the cytoplasm and nucleoplasm (Cohen-Fix and Askjaer 2017; Ungricht and Kutay 2017; Luckner and Wanner 2018; Rahman *et al*. 2020). This process of nuclear envelope breakdown (NEBD) is regulated by CDK1, PLK1 and other mitotic kinases, leading to the disassembly of the NPCs and nuclear lamina (Heald and Mckeon 1990; Laurell *et al*. 2011; Rahman *et al*. 2015; Linder *et al*. 2017; Martino *et al*. 2017; Velez-Aguilera *et al*. 2020). Once mitosis is over, the NE reappears around the chromosomes to establish a single nucleus in each daughter cell, and it then expands during interphase (Lee *et al*. 2000; Cohen-Fix and Askjaer 2017; Ungricht and Kutay 2017).

Defects in nuclear morphology are associated with abnormal nuclear function (Mukherjee *et al*. 2016; Romero-Bueno *et al*. 2019). Mutations in components of the nuclear lamina lead to several diseases (a.k.a. laminopathies) such as Emery-Dreifuss muscular dystrophy, limb girdle muscular dystrophy, dilated cardiomyopathy, Dunnigan-type familial partial lipodystrophy, and Hutchinson-Gilford progeria syndrome (HGPS) (Worman *et al*. 2010; Dobrzynska *et al*. 2016; Donnaloja *et al*. 2020; Pathak *et al*. 2021), all of which are associated with abnormal nuclear shape. Alterations in nuclear morphology have also been associated with cancer (Mukherjee *et al*. 2016), and irregularity in nuclear shape and size is used in cancer diagnosis (Zink *et al*. 2004). Changes in nuclear shape and size have also been observed in normal aging (Haithcock *et al*. 2005) and conditions associated with premature aging (Goldman *et al*. 2004). Importantly, however, whether defects in nuclear structure are the cause or consequence of the above-mentioned disease states and conditions remains to be elucidated. What is missing is an understanding of what regulates nuclear size and shape so that nuclear morphology can be manipulated and the consequences explored.

For these reasons, there is interest in identifying genes that when down-regulated or inactivated lead to changes in nuclear morphology. We chose to do an RNAi screen for genes whose down-regulation affects nuclear morphology using *C. elegans* early embryos: *C. elegans* is amenable to high throughput RNAi screens by feeding, and nuclei of the early embryo are large and nearly spherical, facilitating detection of morphological perturbations. Furthermore, the *C. elegans* strain we used expressed a fluorescently tagged subunit of the NPC, allowing sensitive detection of nuclear shape alterations, as well as alteration in NPC distribution. Finally, unlike siRNA screens done in tissue culture cells, RNAi screens in *C. elegans* are in a physiological developmental context. The high degree of conservation of NE and NPC proteins between nematodes and vertebrates makes it likely that the information garnered from a *C. elegans* screen will help better understand processes that affect mammalian nuclear morphology.

Based on the assumption that defects in nuclear morphology will likely affect embryonic viability, our screen employed RNAi against genes that were reported at the time to be essential for embryonic viability. This screen differed from earlier *C. elegans* screens in two ways: first, some of the earlier screens used either Differential Interference Contrast (DIC) microscopy, which provides less information on nuclear shape abnormalities (Gonczy *et al*. 2000; Piano *et al*. 2000; Zipperlen *et al*. 2001; Sonnichsen *et al*. 2005), or used chromosome detection methods (e.g. DAPI staining, fluorescently-tagged histone proteins; (Colaiacovo *et al*. 2002; Green *et al*. 2011), that only indirectly inform on nuclear morphology. Second, earlier screens that surveyed nuclei encompassed a different set of genes (e.g. genes whose down-regulation leads to sterility (Green *et al*. 2011), genes on a particular chromosome (Fraser *et al*. 2000; Gonczy *et al*. 2000) or examined a different developmental stage (Colaiacovo *et al*. 2002; Green *et al*. 2011). Indeed, our screen uncovered many essential genes and pathways not previously known to affect nuclear morphology. Overall, we have identified 182 genes to be required for normal nuclear morphology in *C. elegans*. This screen has already led two studies describing the involvement of spliceosome components in NPC distribution and the requirement of the TCA cycle in progression past the first embryonic G2 phase (Joseph-Strauss *et al*. 2012; Rahman *et al*. 2014). The present study describes the rest of the results from this screen, including the involvement of unexpected processes, such as the vacuolar ATPase and components of the CCT chaperonin complex, in nuclear morphology.

## Materials and Methods

### Screen design

At the time of primary screening, 1713 genes essential for embryonic development were listed on WormBase and were present in our IPTG-inducible RNAi feeding library (Open Biosystems, Huntsville, AL) (https://wormbase.org/tools/ontology_browser/show_genes?focusTermName=embryonic%20lethal&focusTermId=WBPhenotype:0000050). To monitor the changes in nuclear morphology, the screen was performed on strain OCF3 (unc-119(ed3); jjIs1092[pNUT1::npp-1::gfp + unc-119(+)]; ltIs37 [pAA64: pie-1p::mCherry::his-58 + unc-119 (+)] expressing GFP tagged NPP-1 (NPP-1::GFP), a subunit of the NPC, to visualize the NE, and histone H2B fused to the monomeric red fluorescent protein mCherry (H2B::mCherry) to visualize chromatin (Golden *et al*. 2009). The screen was performed by feeding L4 stage worms with bacteria expressing dsRNA from the RNAi feeding library for 48-52 hours at 20°C. For the control RNAi, worms were fed with dsRNA against *smd-1* (Golden *et al*. 2009). For making RNAi plates, a single colony of dsRNA-expressing bacteria was inoculated in 360 µl Luria-Bertani (LB) liquid medium with carbenicillin (50 µg/ml) and tetracycline (20 µg/ml) in 96 deep-well plates. This culture was spread on RNAi plates containing 6 mM IPTG and carbenicillin (50 µg/ml). For the primary screen, 3 L4s were fed on each RNAi plate and 2 animals were scored for any abnormality in nuclear morphology after 48 hours of treatment. The remaining worm was allowed to lay embryos to determine the embryonic viability. This resulted in 325 RNAi clones with a putative effect on nuclear morphology.

For the secondary screen, the identity of all 325 clones was verified by sequencing. A single colony of dsRNA-expressing bacteria was inoculated in 1-2 ml LB liquid medium with 50 µg/ml of ampicillin and grown overnight at 37°C. On the next day, 0.1 % starter inoculum from the overnight culture was used for inoculating 5 ml of LB liquid medium with 50 µg/ml ampicillin, and was grown at 37°C. After reaching OD_600_ of around 0.5 (∼ 4-5 hours), 10 µl of 0.5 M IPTG were added and the induced culture was grown for ∼ 4 hours at 37°C. The culture was then spun at 5000g for 5 minutes at room temperature and the pellet was resuspended in 1 ml LB liquid medium with 50 µg/ml ampicillin. Around 200 µl of this culture was spread on each RNAi plate and allowed to grow for 1-2 days at 25 °C. The RNAi condition used in secondary screen was found to be more effective than the one in the primary screen. ∼15-20 L4-stage larvae were transferred to RNAi plates, and embryos were examined after 48-52 hours of RNAi. To determine embryonic lethality, 4-5 adult hermaphrodites were transferred to new RNAi plates following 48-52 hours treatment and removed 3–6 hours thereafter. Hatching was scored 24 hours later. The screen was repeated at least 3 times. This narrowed the list to 182 genes whose down-regulation consistently led to one or more nuclear phenotypes. For each treatment, embryos from at least 10 hermaphrodites were scored.

### Scoring nuclear phenotypes

Nuclear phenotypes were scored manually. The vast majority of abnormal nuclei could be classified into 9 different phenotypes which fell into 3 broad categories (Figure1). *Category 1: multi-nucleate cells*, including (i) paired nuclei, (ii) one-cell multi-nucleated embryos, (iii) ≥ 2 cell multi-nucleated embryos, and (iv) micronuclei; *Category 2: abnormally shaped nuclei*, including (i) anaphase bridges and (ii) deformed nuclei; *Category 3: NPC distribution defects*, including (i) abnormal NE distribution of NPCs, (ii) cytoplasmic distribution of NPCs, and (iii) intra-nuclear NPCs (see Results and Discussion for more detail). To qualify as a positive hit, at least 7% of the embryos analyzed per RNAi treatment had to exhibit one or more of these phenotypes.

**Figure 1.**
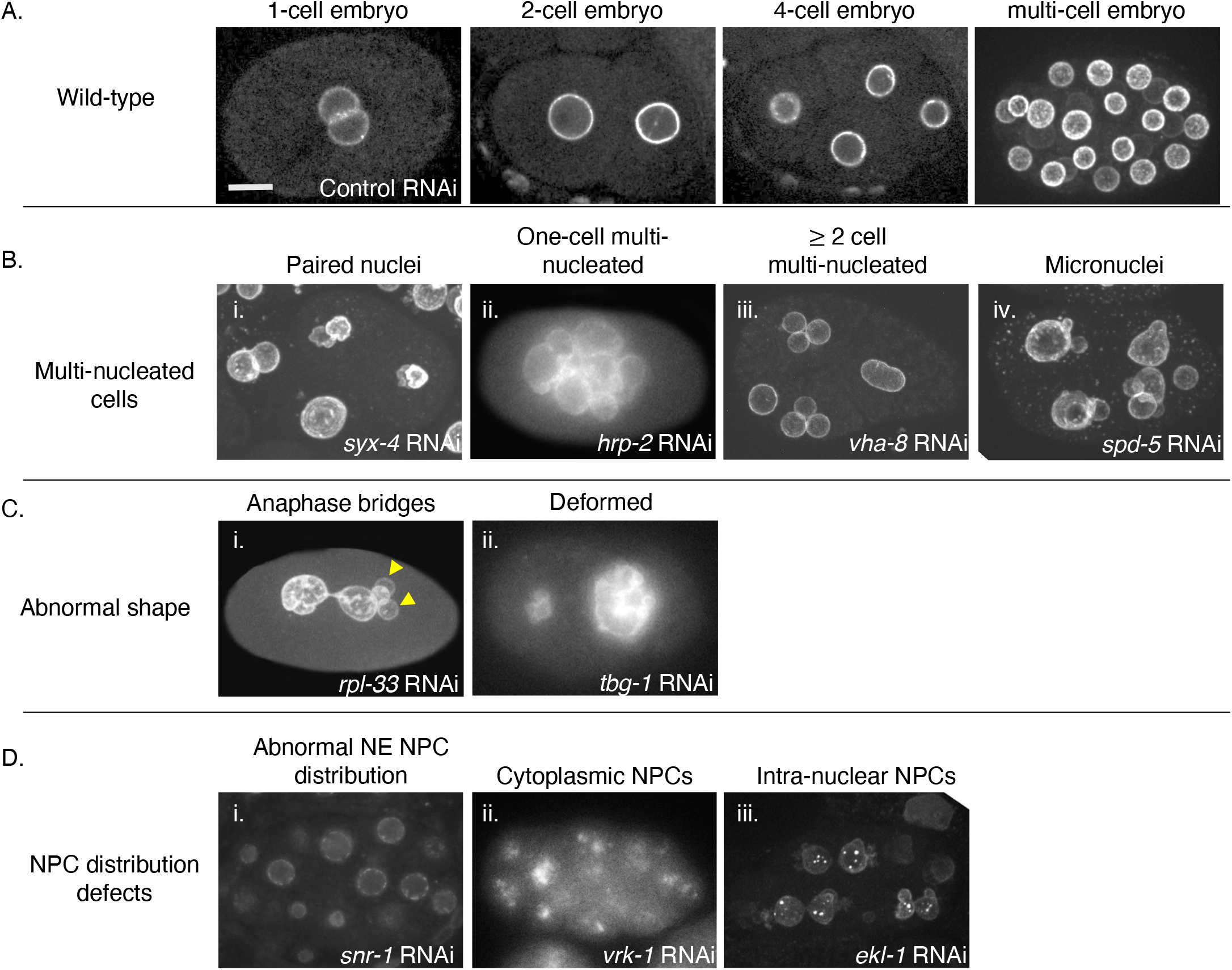
Classes of abnormal nuclear phenotypes. Shown are representative images of *C. elegans* embryos expressing NPP-1 fused to GFP (NPP-1::GFP) from worms treated with RNAi against the indicated genes. **(A)** Wild-type phenotype on control RNAi: examples of 1-, 2-, 4-and multi-cell embryos (some nuclei are on an out-of-focus plane). **(B)** The multi-nucleated cell phenotype: (i) paired nuclei (some nuclei in this case particular are also deformed), (ii) multiple nuclei in a 1-cell embryo, (iii) Multi-nucleated cells in a ≥ 2 cell embryo, and (iv) micronuclei (along with nuclei). **(C)** The abnormal nuclear shape phenotype: (i) anaphase bridges (accompanied here by micronuclei, arrowheads) and (ii) deformed nuclei. **(D)** NPC distribution defect phenotypes: abnormal distribution of NPP-1::GFP (i) on the nuclear envelope, (ii) in the cytoplasm, or (iii) inside the nucleus. Scale bar (for all images)=10 μm.

### Examining ER phenotype

As a tertiary screen, we looked at the endoplasmic reticulum (ER) morphology following RNAi of select genes. This was performed on strain OCF5 (unc-119(ed3); ojIs23 [SP12::GFP + unc-119(+)]; ltIs37[pAA64: pie-1p::mCherry::his-58 + unc-119(+)]) expressing an ER signal peptide, SP12, tagged with GFP and histone H2B tagged with mCherry marking the chromatin (Golden *et al*. 2009). The results of RNAi treatments against *dhc-1, dyci-1, rpl-21, rps-22, cct-2, cct-6, vha-8* and *vha-14* are shown in Figures 6 and 7. The RNAi treatment condition was similar to the one used for the secondary screen.

### Microscopy and image analysis

Images of the primary screen were obtained using a Nikon E800 microscope, equipped with a 60x 1.4 NA Plan Apo objective, using a charge-coupled device camera (CCD; C4742-95; Hamamatsu Photonics) and operated by IPLab 3.9.5 software (BD Biosciences). Images of the secondary screen were obtained using a Nikon Eclipse TE2000U microscope equipped with a 60x 1.4 NA Plan Apo objective. This system was outfitted with a Spectral Applied Research LMM5 laser merge module to control the output of four diode lasers (excitation at 405, 491, 561 and 655 nm), a Yokogawa CSU10 spinning-disk unit, and a Hamamatsu C9100-13 EM-CCD camera. Images were acquired using IPLab 4.0.8 software (BD Biosciences). Images of ER in Figures 6 and 7 were taken using a Nikon confocal Ti2 fitted with a Nikon water 60x 1.2-NA Apo Plan objective, a Yokagawa CSU-X1 spinning disk and a photometrix Prime 95B camera. Image acquisition was done using Elements software (Nikon Instruments, Inc.). Image processing was done with FIJI (ImageJ) (Schindelin *et al*. 2012).

### Gene Ontology (GO) overrepresentation analysis

Statistical overrepresentation analysis of Gene Ontology (GO) terms was analyzed using PANTHER Classification System v16.0 (http://pantherdb.org; (Mi *et al*. 2019). WormBase IDs were entered for input and the Fisher’s Exact test with the default false discovery rate (FDR) calculation settings was used to determine enriched GO terms.

## Results and Discussion

### Down-regulation of 182 *C. elegans* genes led to abnormal nuclear morphology or abnormal NPC distribution

To identify genes whose down-regulation affects nuclear morphology, we carried out a large-scale RNAi screen in *C. elegans*, visualizing nuclei in embryos. We targeted genes essential for embryonic development reasoning that severe alterations in nuclear morphology would likely result in embryonic lethality. At the time this screen was initiated, nearly 2000 genes, when down-regulated, were known to cause embryonic lethality. Of those, 1713 were present in our RNAi feeding library. To monitor the changes in nuclear morphology, the screen was performed using a transgenic line expressing *C. elegans* homolog of Nup54 (NPP-1), a subunit of the NPC, fused to GFP (NPP-1::GFP) to visualize the NE, and histone H2B fused to the monomeric red fluorescent protein mCherry (H2B::mCherry) to visualize chromatin (Golden *et al*. 2009; Joseph-Strauss *et al*. 2012). NPP-1::GFP also allowed us to identify the genes that affect NPC distribution.

The screen was performed by feeding L4 stage worms with bacteria expressing dsRNA from the RNAi feeding library, as described under Materials and Methods. RNAi against a large set of genes, about two thirds of the entire collection, did not lead to a robust embryonic lethal phenotype. There could be multiple reasons for this: first, the efficiency of feeding RNAi is likely less than that of other forms of gene inactivation (e.g. dsRNA injection, mutations) on which the reported embryonic lethal phenotype could have been based. Second, the time of RNAi treatment was not optimized, and it is possible that longer exposures may have resulted in a higher faction of RNAi constructs causing robust lethality. Finally, our cutoff for an embryonic lethal phenotype was ≥15% of dead embryos, and in theory some of the reported embryonic lethal phenotypes could have been below this rate. In total, down-regulation of 529 genes by RNAi caused embryonic lethality of at least 15%. Of those, in the primary screen, down-regulation of 325 genes by RNAi led to abnormal nuclear morphology. Of note, the abnormal nuclear morphology was not a non-specific result of embryonic lethality because (a) we observed a variety of abnormal nuclear phenotypes (see below) and (b) not all RNAi treatments that led to embryonic lethality also resulted in abnormal nuclear morphology. None of the RNAi treatments that did not cause embryonic lethality resulted in an abnormal nuclear morphology. In addition, 25 genes, when down-regulated, led to phenotypes that precluded assessing the nuclear morphology of an early embryo, such as meiotic arrest, unfertilized embryos or lack of NPP-1::GFP expression. The initial 325 genes were rescreened to verify the phenotype(s) and ensure reproducibility, as described under Materials and Methods. This narrowed down the list to 182 genes that, when down-regulated, consistently exhibited a robust nuclear morphology phenotype, as discussed below.

### Nuclear morphology defects could be divided into 3 major categories

In wild-type *C. elegans* early embryos, nuclei are spherical and the NPCs are distributed evenly around the nuclear envelope, as shown by NPP-1::GFP localization (Figure 1A) (Golden *et al*. 2009). In our screen, deviation from the normal nuclear phenotype resulted in one or more defects that could be divided into three major phenotypic categories. It should be noted that many RNAi treatments resulted in multiple nuclear phenotypes (see Figure 1 for examples and also below), which is not uncommon (Fraser *et al*. 2000).

#### Category 1: multi-nucleate cells (Figure 1B)

This category included: (i) Paired nuclei, as seen by the presence of two attached nuclei in each cell, likely due to a defect in pronuclear fusion after fertilization (Audhya *et al*. 2007; Galy *et al*. 2008; Nishi *et al*. 2008; Rivers *et al*. 2008; Golden *et al*. 2009; Gorjánácz and Mattaj 2009; Noatynska *et al*. 2010; Rahman *et al*. 2015; Rahman *et al*. 2020); (ii) one-cell multi-nucleated embryos, likely caused by multiple nuclear divisions without intervening cytokinesis (Swan *et al*. 1998; Gonczy *et al*. 1999; Echard *et al*. 2004); (iii) ≥2 cell multi-nucleated embryos, which could be due to cytokinesis defects at later stage embryos (Green *et al*. 2011); and (iv) micronuclei, one or more smaller sized nuclei along with a relatively regular-sized nucleus, which could have resulted from chromosome fragmentation or defective chromosome segregation (Strome *et al*. 2001).

#### Category 2: abnormally shaped nuclei (Figure 1C)

This category included: (i) anaphase bridges, likely a result of defects in chromosome segregation, such as improper attachment of chromosomes to the spindle or chromosome catenation that precludes full segregation (chromatin image data not shown) (Moore *et al*. 1999; Hagstrom *et al*. 2002; Bembenek *et al*. 2013); and (ii) deformed nuclei, which could have resulted from defective lamina, alteration in chromatin architecture or defects in cytoskeleton, among other reasons.

#### Category 3: NPC distribution defects (Figure 1D)

This category included: (i) abnormal NE distribution of NPCs, shown by the mislocalization of NPP-1::GFP at the nuclear envelope, as described previously (Joseph-Strauss *et al*. 2012); (ii) cytoplasmic distribution of NPCs, as seen by cytoplasmic localization of NPP-1::GFP, possibly due to defects in nuclear envelope reassembly or proper NPC subunit targeting, and (iii) intra-nuclear NPCs, as seen by the mislocalization of NPP-1::GFP inside the nucleus and that could have also resulted from an improper nuclear envelope reassembly and perhaps internal nuclear membrane structures (Figure 1D).

Embryos from each RNAi treatment were scored for these 9 phenotypic categories (Supplemental Table S1). Out of the 182 genes whose down-regulation resulted in a consistent abnormal nuclear morphology, the most common defect was the multi-nucleated cell phenotype (RNAi against 147 genes). Down-regulation of 119 genes caused an abnormal nuclear shape, while down-regulation of 39 genes caused NPC distribution defects (Supplemental Table S1). As noted above, in most cases down-regulation of one gene resulted in multiple nuclear phenotypes, and only about a third of the RNAi treatments resulted in a single abnormal nuclear phenotype (Figure 2A). For example, RNAi against *cks-1*, a cyclin-dependent kinase (Polinko and Strome 2000), led to 7 different nuclear phenotypes (Supplemental Table S1). Depletion of ∼30 percent of genes led to at least 2 nuclear phenotypes, and ∼25 percent resulted in 3 different nuclear phenotypes (Figure 2A). Some of the phenotypes tended to occur together (Figure 2B and Supplemental Table S2). For example, deformed nuclei were concurrent in 50% of embryos with paired nuclei (26/50), ∼ 70% of ≥ 2-cell multi-nucleated embryos (71/106), ∼ 75% of one-cell multi-nucleated embryos (9/13) and ∼ 75% of embryos with micronuclei (40/54). Micronuclei were most common in embryos also exhibiting anaphase bridges (18/33, 55%), followed by embryos with deformed nuclei (40/104, 38%), ≥ 2-cell multi-nucleated embryos (36/106, 33%), 1-cell multi-nucleated embryos (4/13, 31%) and embryos with paired nuclei (12/50, 24%). RNAi treatments that led to uneven NPC distribution exhibited few other nuclear phenotypes, but those that led to cytoplasmic or intra-nuclear NPCs were often accompanied by other nuclear defects, and in particular multi-nuclei in ≥2 cell embryos (54% for embryos with cytoplasmic NPCs (7/13) and 100% for embryos with intra-nuclear NPCs (2/2).

**Figure 2:**
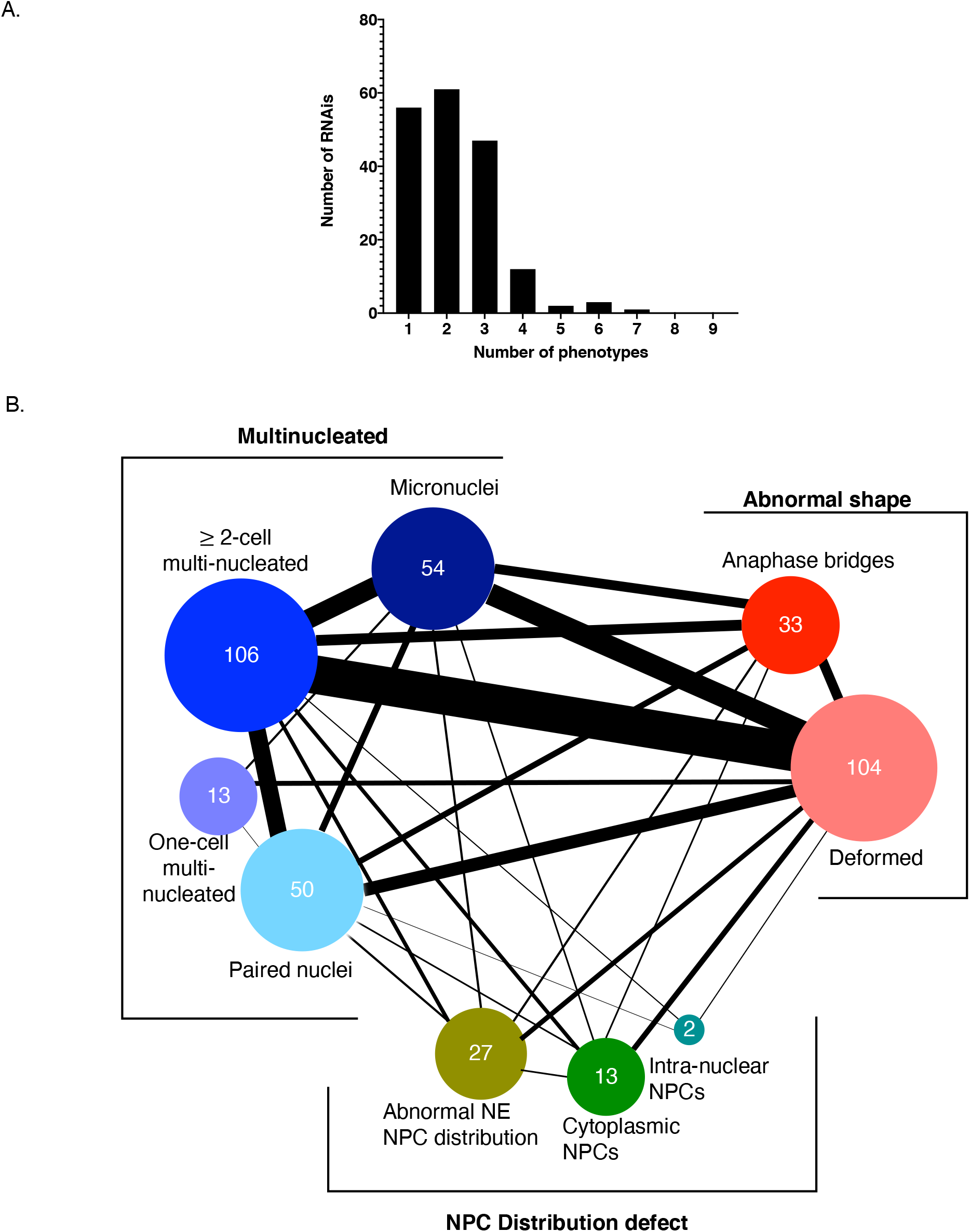
Distribution of abnormal nuclear phenotypes among the 182 RNAi treatments. (A) Bar graph showing the number of RNAi treatments that resulted in the indicated number of phenotypes. (B) Connectivity of phenotypic categories. The thickness of each connecting line corresponds to the number of genes that, when down-regulated, result in both phenotypes. See Supplemental Table S2 for the exact number of shared phenotypes.

### The relationship between phenotype and gene function

Our screen uncovered multiple genes in 13 distinct functional categories, plus a number of genes that could not be placed in a larger functional category (Figure 3A and Supplemental Table S1). Moreover, the screen uncovered clusters of functionally related genes, either within the same complex same or in the same pathway. To determine the significance of these data, we analyzed the biological functions of our gene list by manual curation based on primarily *C. elegans* literature and homology to genes in other organisms with known or predicted function (Figure 3A and Supplemental Table S1). Our functional assignments were in good agreement with the ascribed functions on WormBase. To determine the overrepresentation of a functional class in our dataset compared to the *C. elegans* genome, we performed a statistical overrepresentation test of Gene Ontology (GO) terms associated with those 182 genes using PANTHER Classification System v16.0. PANTHER recognized 181 genes out of 182 (Mi *et al*. 2019). Figures 3B-D show some of the most highly enriched biological process, cellular component, and molecular function classes (the complete lists, including p values, are shown in Supplemental Tables S3-S5). Mitochondrial ATP synthesis coupled transport (GO:0042776) is the most enriched GO biological process (> 87 fold enrichment) relative to the *C. elegans* reference genome among the biological classes (Figure 3B and Supplemental Table S3). Consistently, the mitochondrial functions gene class formed one of the largest in our manual classification (Figure 3A and Supplemental Table S1). Lysosomal acidification was highly enriched among the biological process, cellular component and molecular function classifications (Figures 3B-D and Supplemental Table S3-S5). There is no doubt that our screen has missed many genes that could affect nuclear morphology, either because they were not present in our RNAi collection or because the feeding RNAi treatment did not sufficiently down-regulate their activity. Nonetheless, the genes that were uncovered serve as a basis for further investigation into the involvement of specific processes in affecting nuclear morphology.

**Figure 3.**
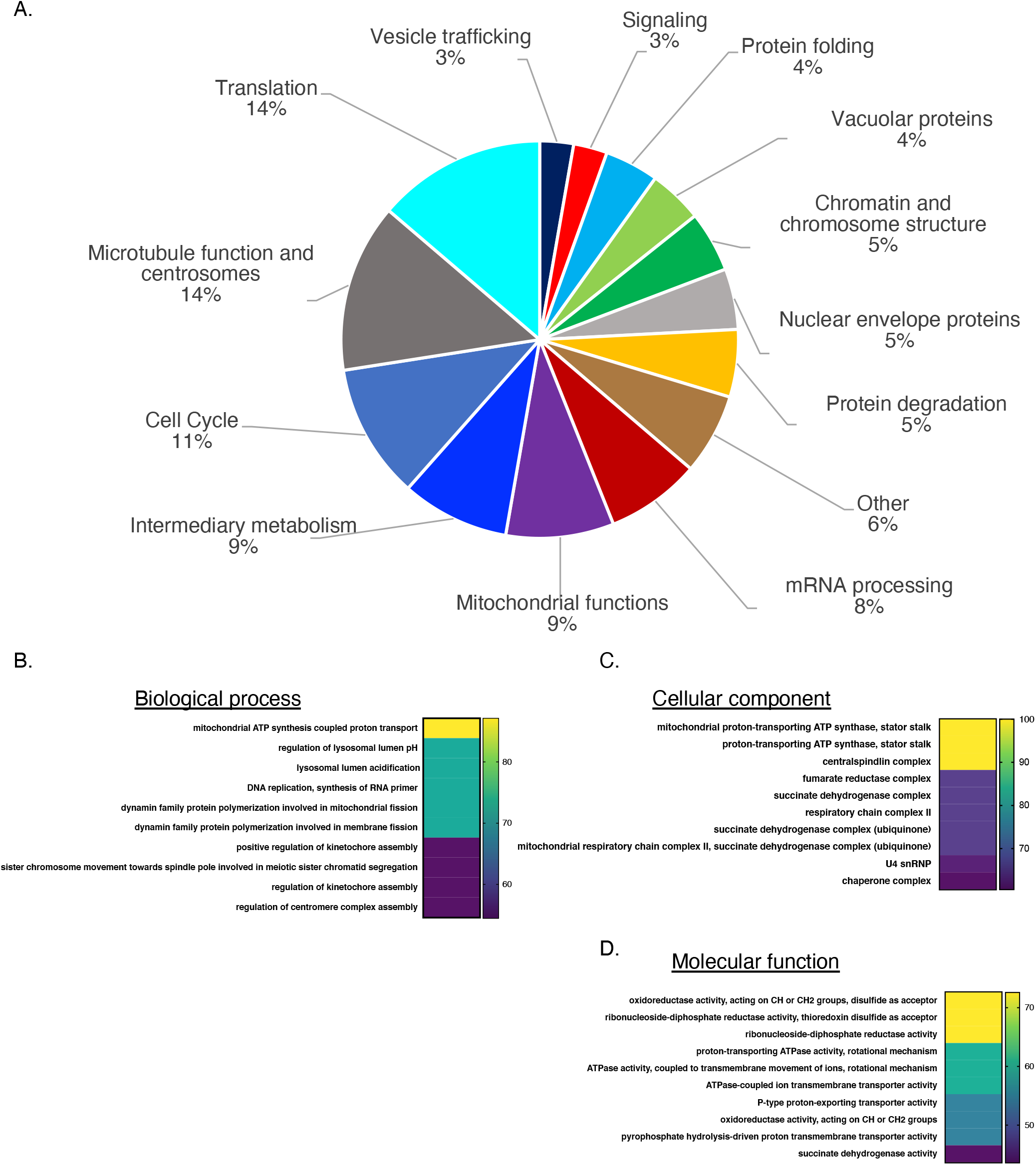
Functional categories of genes whose down-regulation results in one or more abnormal nuclear or NPC phenotype. **(A)** The relative abundance (in percentage) of each functional category within the 182 hits is indicated. **(B, C, D)** Statistical overrepresentation analysis of Gene Ontology (GO) terms using the PANTHER Classification System. The top statistically significant overrepresented subclasses based on Biological Process **(B)**, Cellular Component **(C)**, and Molecular Function classes **(D)** are shown. Number on the right represent fold enrichment. See Supplemental Tables S3-S5 for the complete list and p values.

As alluded to above, our screen uncovered genes in processes whose involvement in nuclear morphology was unexpected. We previously reported the surprising link between down-regulation of splicing factors and a defect in NPC distribution [(Joseph-Strauss *et al*. 2012); Supplementary Table S1], and between TCA cycle genes and the accumulation of embryos in the 1-cell stage with paired nuclei due to a block in mitotic progression [(Rahman *et al*. 2014); Supplementary Table S1]. In addition, we found that down-regulation of translation, chaperonin, vacuolar function, intermediary metabolism, and mitochondrial function also affected nuclear morphology (Figures 3A and 4; Supplementary Table S1). This revealed that a defect in a variety of core cellular processes can cause alteration in nuclear morphology, a finding that could be relevant to the abnormal nuclear morphology often seen in cancer cells. Below we discuss the relationship between nuclear morphology and several of the functionally related genes. Importantly, different functional categories displayed distinct patterns of abnormal nuclear phenotypes: for example, the most prevalent phenotype of the protein degradation functional group was deformed nuclei, whereas the most common phenotypes of the mRNA processing and the chromatin and chromosome functions groups were abnormal NE NPC distribution and anaphase bridges, respectively (Figure 4). This supports the conclusion that the abnormal nuclear phenotype is not merely a consequence of dying cells, and suggests that distinct processes impact the nucleus differently. Interestingly, we observed similar phenotypic signatures in disparate functional categories, such as the ones resulting from the inactivation for protein folding genes and vacuolar genes (Figure 4), suggesting that the underlying cause for abnormal nuclear morphology may be similar. In contrast, the NPC distribution defect phenotypes were mostly seen when mRNA processing and nuclear envelope genes were down-regulated, but the rest of the phenotypes of these two functional groups are quite dissimilar.

**Figure 4.**
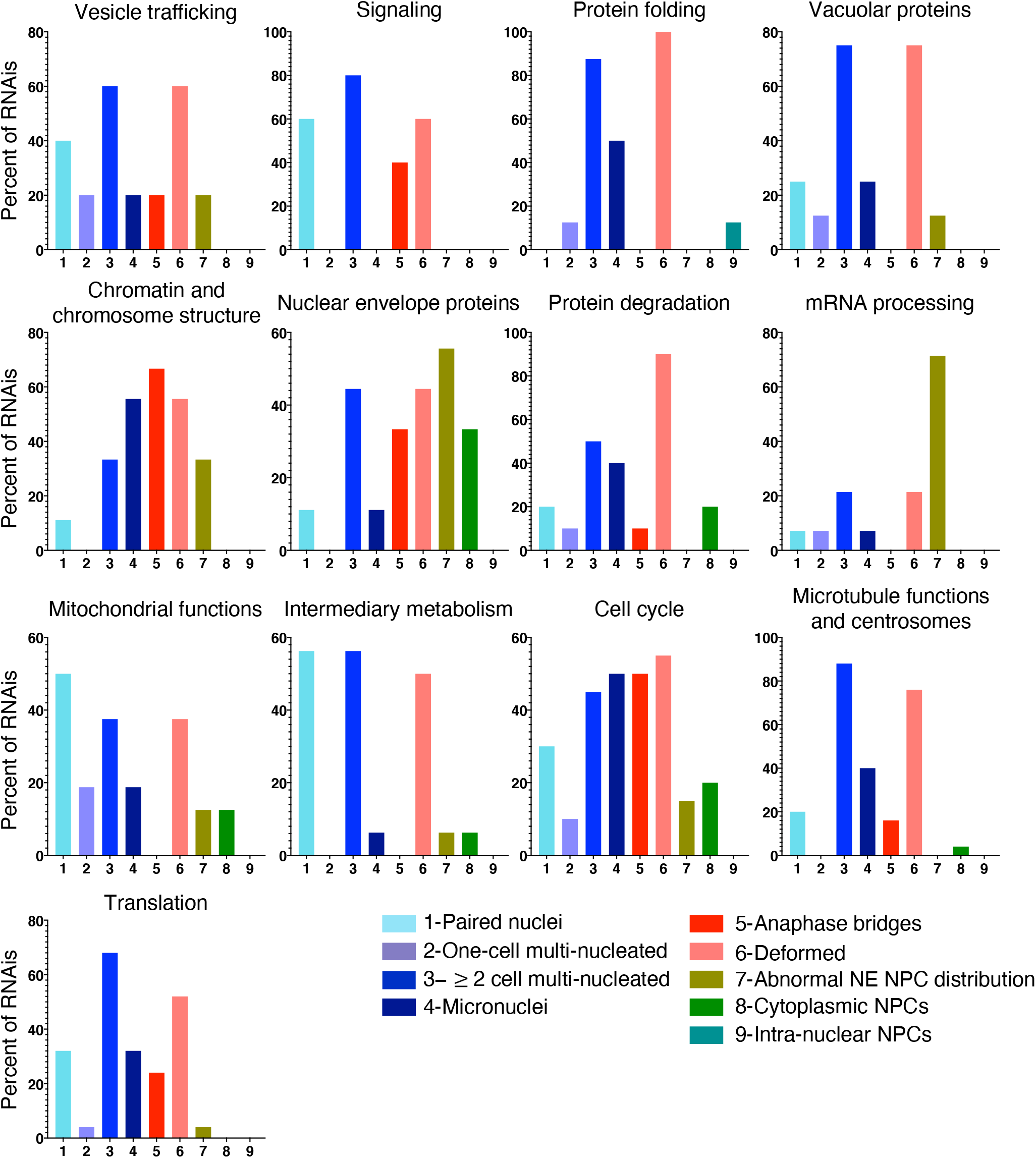
Functional categories of genes whose down-regulation results in one or more abnormal nuclear or NPC phenotype. The prevalence of each of the 9 phenotypes observed within each functional category: Phenotypic categories are on the X axis, and the percent of RNAis within a functional category displaying a particular phenotype is on the Y axis.

### Cell cycle and chromosome segregation-related functions

Cell cycle genes were expected to affect nuclear morphology, and down-regulation of genes in this category sometimes led to pleotropic effects as a consequence of their involvement in a wide range of processes (Figure 4 and Supplemental Table S1). Within the cell cycle functional category, DNA replication genes and kinetochore genes, when down-regulated, led predominantly to anaphase bridges (7/7 genes, Supplemental Table S1), likely because of defects in segregating incompletely replicated of mis-aligned chromosomes. Not surprisingly, nuclear morphology was affected by the down-regulation of numerous genes associated with spindle/microtubule function, such as genes coding for of α and β tubulin (*tba-2, tba-4* and *tbb-2*), centrosome proteins (*tbg-1, gip-2, spd-5, sas-5, sas-6* and *syz-4*) and dynein and dynactin subunits (*dlc-1, dyci-1, dli-1, nud-1* and *dnc-1*), as well as other motor proteins (*zen-4, klp1-18* and *klp-19*) (Supplemental Table S1). These genes are involved in the precise positioning and orientation of the mitotic spindle necessary for both chromosome segregation and cytokinesis (Carminati and Stearns 1997; Shaw *et al*. 1997; Skop and White 1998; Gonczy *et al*. 1999; Karki and Holzbaur 1999; Dujardin and Vallee 2002; Nguyen-Ngoc *et al*. 2007; Moore *et al*. 2008; Markus and Lee 2011; Collins *et al*. 2012; Laan *et al*. 2012; Kotak and Gonczy 2013; Pintard and Bowerman 2019). Indeed, down-regulation of 88% of genes (22 out of 25) in the microtubule and centrosome function category exhibited multi-nucleated cells in embryos with 2 or more cells (Figure 4 and Supplemental Table S1). Likewise, the most common phenotypes observed when genes within the “chromatin and chromosome structure” category were down-regulated were micronuclei, anaphase bridges and deformed nuclei, consistent with the effect of chromosome mis-segregation on nuclear morphology (Figure 4 and Supplemental Table S1).

### Protein translation

The translation machinery is comprised of ribosomal subunits, tRNA aminoacylation factors, translation factors, and more. This machinery also depends on genes that regulate the biogenesis of these factors, such as ribosome biogenesis genes. Down-regulation of genes in this class has also appeared in other *C. elegans* genome-wide screens, displaying defects such as multi-nucleated embryos, aberrant cytoplasmic texture and osmotic sensitivity (Gonczy *et al*. 2000; Sonnichsen *et al*. 2005; Green *et al*. 2011). We also observed a multi-nucleated phenotype: Down-regulation of 21 of 25 genes involved in translation resulted in at least one of the multi-nucleated category phenotypes (Figure 4 and Supplemental Table S1). Moreover, Down-regulation of 7 of 10 ribosomal genes and 5 of 7 tRNA synthetase genes (Supplemental Table S1) resulted in ≥ 2-cell multi-nucleated embryos. The range of phenotypes within the “translation” functional category was not uniform (Supplemental Table S1). For example, down-regulation of over half of the ribosomal subunits picked up in our screen resulted in anaphase bridges (5/10, Supplemental Table S1). In contrast, none of seven different tRNA synthetases, and only 1 of 6 translation factors, exhibited anaphase bridges when down-regulated (Supplemental Table S1). Thus, while all these factors participate in translation, depletion of ribosomal proteins appear to affect the nucleus in ways that tRNA synthetases and translation factors do not (Supplemental Table S1). How this group of genes affect nuclear morphology is not known. A defect in translation could have resulted in a reduced level of one or more proteins required for nuclear integrity, making the involvement of translation in nuclear morphology indirect. Conversely, the association of ribosomes with the ER may have a direct effect on membrane conformation, such that a defect in the translation machinery could deform the ER, resulting, conceivably, in difficulties to undergo cytokinesis and leading to a multi-nuclear phenotype. The effect of down-regulation of the translation machinery and genes from other functional categories on ER structure is examined below.

### Chaperonin

The eukaryotic cytosolic chaperonin containing TCP-1 (CCT) has a role in the folding of cytoskeletal components (Horwich *et al*. 2007). There are 8 CCT subunits (cct-1 to cct-8) encoded by the *C. elegans* genome (Leroux and Candido 1995; Leroux and Candido 1997; Matus *et al*. 2010). They are involved in tubulin folding and microtubule growth in the early embryo (Srayko *et al*. 2005; Lundin *et al*. 2008). Our screen picked up 5 *cct* genes: *cct-1, cct-2, cct-5, cct-6 and cct-7*. Down-regulation of each of the 5 resulted in multi-nucleated embryos with deformed nuclei (Supplementary Table S1). This pattern of phenotypes is similar to that observed for defects in microtubule function and centrosomes (Figure 4), consistent with the nuclear morphology defect due to microtubule malfunction when *cct* genes are down-regulated.

### Vacuolar ATPase

Several subunits of the vacuolar-type H+ ATPase (V-ATPase) were also picked up in our screen. The V-ATPase is a large protein complex that acidifies intracellular compartments (Nelson 1992). These proteins comprise of two functional domains V1 (8 subunits A-H) and V0 (6 subunits a, c, c’, c”, d and e). The worm genome encodes all the subunits except c′. Our screen has picked up 8 of these subunits: *vha-1* (V0 subunit c), *vha-2* (V0 subunit c), *vha-4* (V0 subunit b), *vha-8* (V1 subunit E1), *vha-9* (V1 subunit F), *vha-11* (V1 subunit C1), *vha-14* (V1 subunit D) *and unc-32* (V0 subunit a1) [reviewed in (Lee *et al*. 2010)]. Interestingly, when down-regulated, all but one resulted in a multi-nucleated phenotype (Supplementary Table S1). Previously, Choi et al (Choi *et al*. 2003) reported a clustered nuclei phenotype equivalent to our multi-nucleated phenotype upon VHA-8 depletion. Furthermore, down-regulation of 6 of the 8 *vha* genes that we isolated led to ≥ 2-cell multi-nucleated embryos that were often accompanied by deformed nuclei (Figure 4 and Supplementary Table S1). This was reminiscent of the range of phenotypes obtained when down-regulating microtubule and centrosome function genes (Figure 4), suggesting that defects in vacuolar function may result in chromosome segregation and/or cytokinesis defects.

Overall, different phenotypes were predominantly caused by defects in distinct cellular functions (Figure 5). For example, genes related to microtubule and centrosome functions made up 19% and 21% of genes that, when down-regulated, resulted in micronuclei and ≥2-cell multi-nucleated embryos, respectively, whereas they only made up 10% of paired nuclei embryo, and none of the 1-cell multi-nucleated embryos (Figure 5, left column). Not surprisingly, the relative weight of the functional categories that affected NPC distribution was distinct from those that affected nuclear morphology (Figure 5, compare right column with the middle and left columns): mRNA processing genes were prominent in the “abnormal NE NPC distribution” category (37%), but only a few percent or none at all in all other categories. Mitochondrial function genes also appeared in a subset of phenotypes. Thus, while most functional groups, when down-regulated, could lead to many different phenotypes, each phenotype had its own unique combination of functional groups.

**Figure 5.**
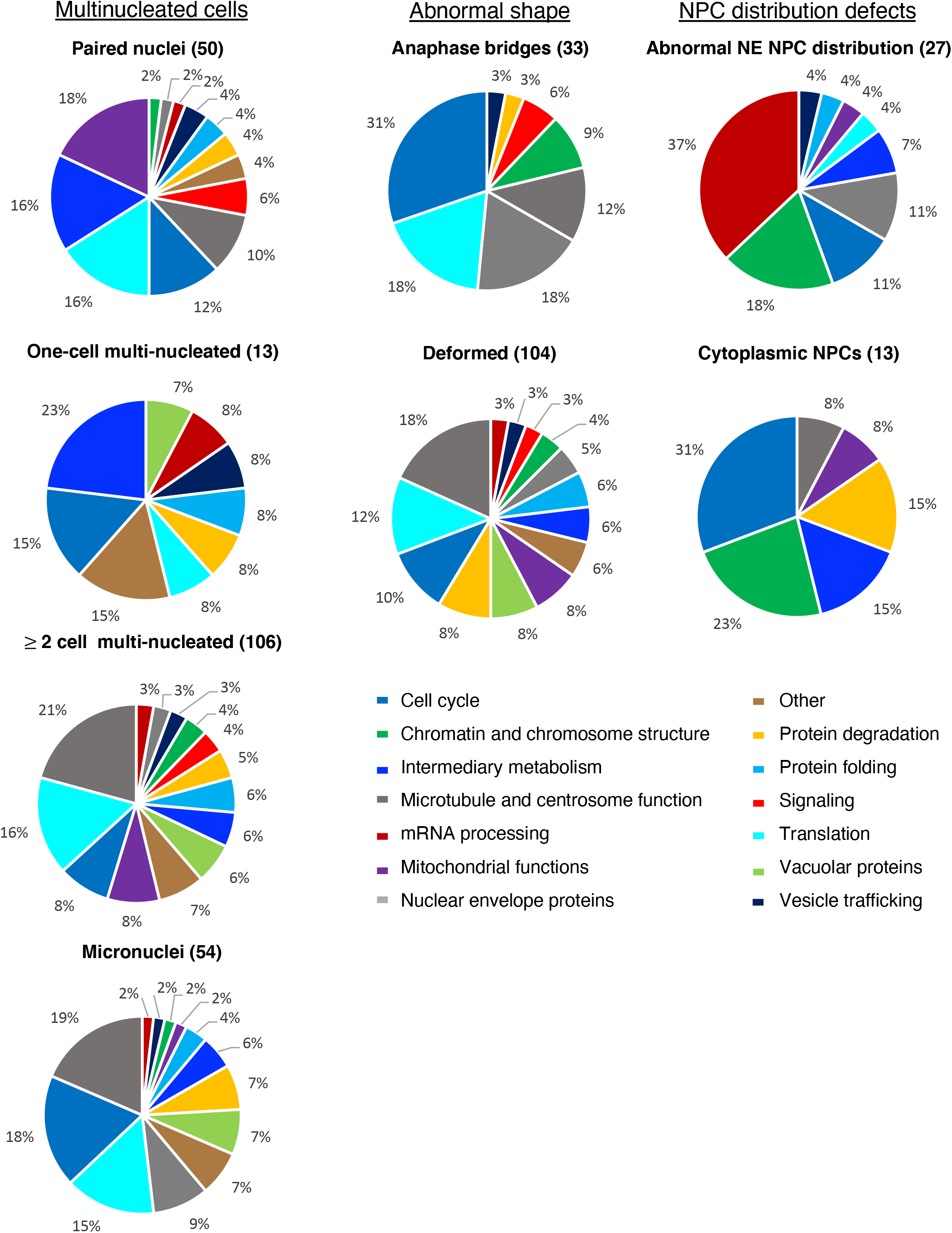
Different phenotypes are a result of the down-regulation of different sets of genes. The contribution of each functional group to a given phenotype, as determined by the number of genes from a particular functional category that, when down-regulated, resulted in the indicated phenotypes. For example, 37% of genes that lead to abnormal NPC NE distribution (top right) are involved in mRNA processing (red sector). The breakdown for the “Nuclear NPC” category is not shown as it encompasses only 2 genes.

### Down-regulation of some, but not all, genes that affect nuclear morphology also affects ER morphology

To gain further insight into how some of the unexpected functional groups may affect nuclear morphology, we examined the effect of their down-regulation on ER organization, as the NE is continuous with the ER (Watson 1955). The underlying hypothesis was that if down-regulating genes of a particular functional category also affected ER morphology, then the effect on nuclear morphology could by affecting ER/nuclear membrane properties. An example of such a case is the down-regulation of lipin (*lpin-1* in *C. elegans*), a phosphatidic acid phosphatase that converts phosphatidic acid to diacylglycerol (Siniossoglou *et al*. 1998; Tange *et al*. 2002; Santos-Rosa *et al*. 2005; Campbell *et al*. 2006; Han *et al*. 2006). In the absence of lipin activity the total amount of cellular phospholipids increases, resulting the ER expansion (Santos-Rosa *et al*. 2005). Down-regulation of *C. elegans lpin-1* results in abnormal ER morphology and deformation of the nucleus (Golden *et al*. 2009; Gorjánácz and Mattaj 2009).

To this end, we repeated select RNAi treatments on worms expressing the ER signal peptide SP12 tagged with GFP (SP12::GFP) (Poteryaev *et al*. 2005). The ER in the early embryo forms a thin reticulate pattern with only rare patches of denser/sheet ER [Figure 6, control RNAi and (Poteryaev *et al*. 2005)]. Under our RNAi conditions, down-regulation of dynein and dynactin, a microtubule-dependent motor, resulted in largely normal ER with a mostly reticular structure, despite the abnormal number of nuclei (Figure 6, *dnc-1* and *dyci-1* RNAi). This is perhaps not surprising given that the underlying defect is likely a failure in spindle positioning and cytokinesis. In contrast, down-regulation of two different ribosome subunits, *rpl-21* and *rps-22*, which also resulted in multi-nucleated cells, did affect ER structure, as seen by the increased abundance of ER patches and whorls (Figure 7A). This suggest that the nuclear defect seen when translation is down-regulated could be a consequence of an ER defect. We speculate a reduced abundance of ribosomes may affect ER morphology indirectly, due to a decrease in translation of one or more ER shaping factors or the induction of stress than may affect ER morphology (Mateus *et al*. 2018). Alternatively, a reduction in the number of ribosomes may affect ER morphology directly, as ribosomes associate with the ER and may contribute to its organization (Shibata *et al*. 2010).

**Figure 6.**
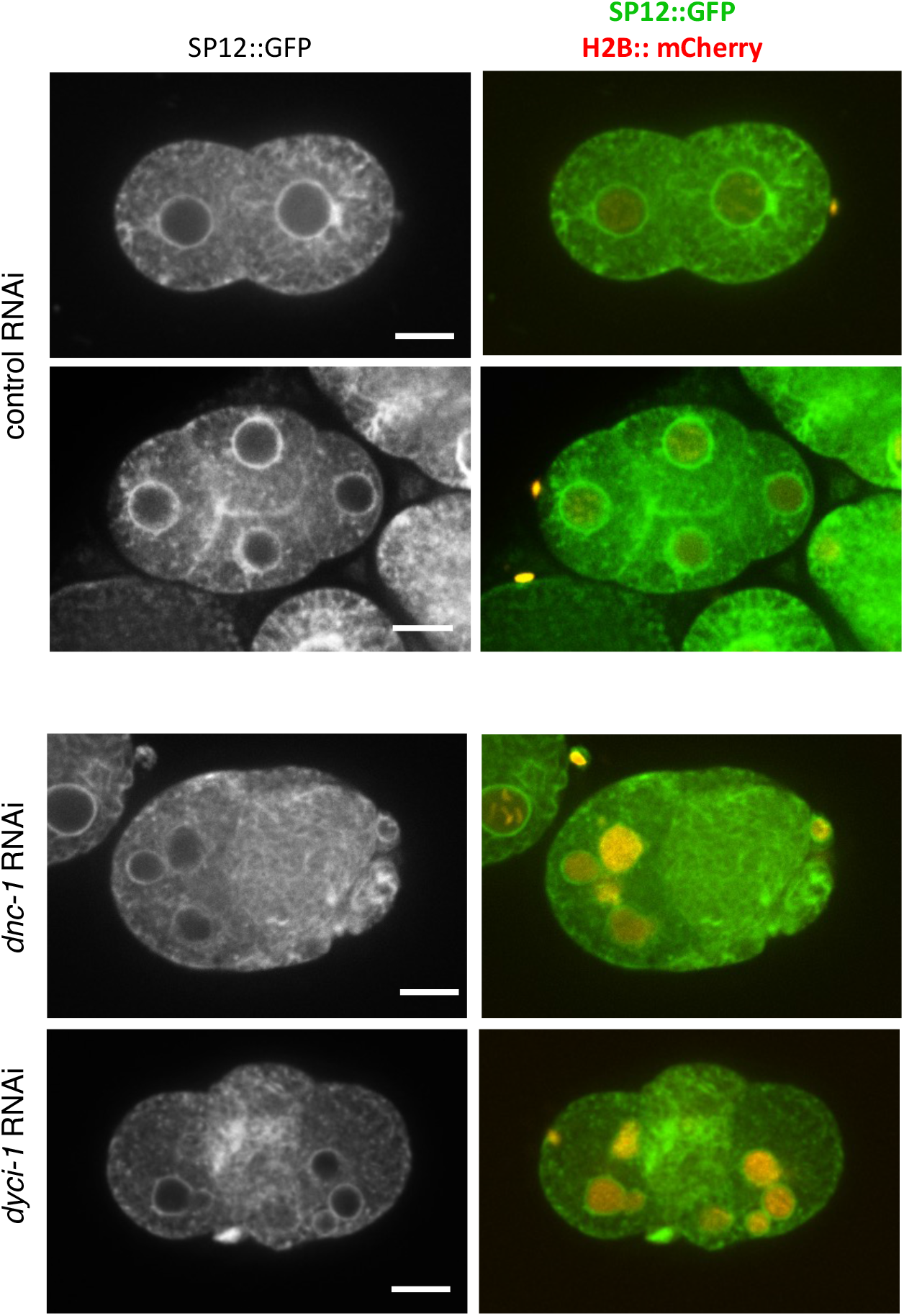
Inactivation of dynein/dynacin proteins has minimal effect on ER structure. Examples of *C. elegans* embryos expressing SP12::GFP (an ER marker) and histone H2B:: mCherry (chromatin), treated with control RNAi against *smd-1* (top two rows) or RNAi against dynacin (*dnc-1*) or dynein (*dyci-1*) subunits (bottom two rows). Scale bar= 10 *µ*m.

**Figure 7.**
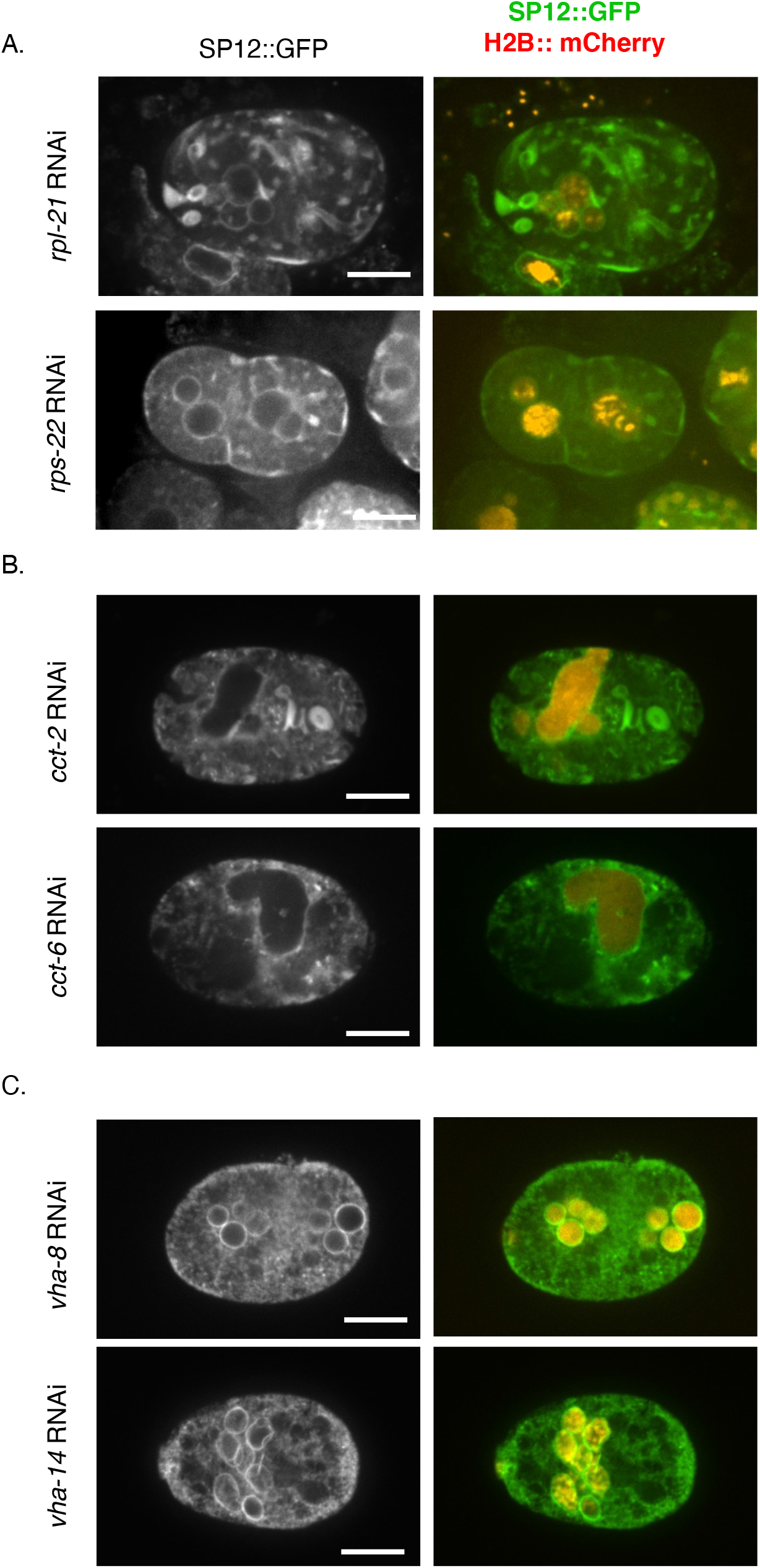
The effects of down-regulating translation machinery, chaperonin and the vacuolar ATPase on ER morphology. Representative images of *C. elegans* embryos as in Fig. 6 following RNAi treatment against genes involved in translation **(A)**, chaperonin **(B)** or vacuolar ATPase **(C)**. Scale bar= 10 *µ*m.

Chaperonin is a chaperon required for actin and tubulin folding (Horwich *et al*. 2007). Down-regulation of chaperonin subunits results in deformed nuclei that are often enlarged (Figure 7B), a phenotype that is often seen when tubulin is down-regulated (Supplemental Table S1) and may be due to failure in chromosome segregation coupled with additional rounds of DNA replication. Chaperonin down-regulation also results in abnormal ER morphology (Figure 7B). Since actin is involved in maintaining ER morphology (Poteryaev *et al*. 2005) down-regulation of chaperonin may affect nuclear and ER morphologies independently.

Finally, we examined the ER following down-regulation of the vacuolar ATPase. Unlike down-regulation of ribosome subunits of chaperonin, the ER in embryos down-regulated for VHA subunits largely retained its reticular structure and did not exhibit extensive areas of ER patches or whorls (Figure 7C). The void regions within the ER may reflect an abnormal ER organization, or they could be regions occupied by large vesicles, such as lysosomes, not visible with the markers that were used. The overall reticular structure of the ER combined with the preponderance of embryos with multi-nucleated cells was reminiscent of the phenotype seen when spindle function and/or cytokinesis was defective (Figure 6 and Supplemental Table S1). This suggests that defects in the worm vacuolar ATPase may affect spindle function, positioning and/or the machinery responsible for cytokinesis. Further studies are required to elucidate the connection between translation, chaperonin and vacuolar ATPase and nuclear morphology.

Taken together, we have shown that RNAi against 182 genes, known for their requirement during embryogenesis, results in 3 major phenotypic categories related to nuclear morphology or NPC distribution. Most of these genes can be subdivided into 13 functional categories, and in several cases our screen uncovered genes for components of the same complex or pathway, all of which affect nuclear morphology or NPC distribution in a similar way. Some of these functional categories were not previously known to affect nuclear morphology. This is significant because one of the hallmarks of cancer is abnormal nuclear size and/or shape, and our data suggest that the underlying defects may originate in a wide range of biological processes. The screen has clearly missed some genes, but the list provided in this study can serve as a solid foundation for the discovery of additional proteins and processes that affect nuclear morphology, especially when partially or fully depleted.

## Acknowledgements

The authors thank the members of the Cohen-Fix lab for advice and support. The authors were supported by an intramural grant from NIDDK #DK069012. The authors declare no competing financial interests.

## References

Audhya, A., A. Desai and K. Oegema, 2007 A role for Rab5 in structuring the endoplasmic reticulum. J Cell Biol 178: 43–56.

Bembenek, J. N., K. J. Verbrugghe, J. Khanikar, G. Csankovszki and R. C. Chan, 2013 Condensin and the spindle midzone prevent cytokinesis failure induced by chromatin bridges in C. elegans embryos. Curr Biol 23: 937–946.

Campbell, J. L., A. Lorenz, K. L. Witkin, T. Hays, J. Loidl et al., 2006 Yeast nuclear envelope subdomains with distinct abilities to resist membrane expansion. Molecular biology of the cell 17: 1768–1778.

Carminati, J. L., and T. Stearns, 1997 Microtubules orient the mitotic spindle in yeast through dynein-dependent interactions with the cell cortex. J Cell Biol 138: 629–641.

Choi, K. Y., Y. J. Ji, B. K. Dhakal, J. R. Yu, C. Cho et al., 2003 Vacuolar-type H+-ATPase E subunit is required for embryogenesis and yolk transfer in Caenorhabditis elegans. Gene 311: 13–23.

Cohen-Fix, O., and P. Askjaer, 2017 Cell biology of the Caenorhabditis elegans Nucleus. Genetics 205: 25–59.

Colaiacovo, M. P., G. M. Stanfield, K. C. Reddy, V. Reinke, S. K. Kim et al., 2002 A targeted RNAi screen for genes involved in chromosome morphogenesis and nuclear organization in the Caenorhabditis elegans germline. Genetics 162: 113–128.

Collins, E. S., S. K. Balchand, J. L. Faraci, P. Wadsworth and W.-L. Lee, 2012 Cell cycle–regulated cortical dynein/dynactin promotes symmetric cell division by differential pole motion in anaphase. Mol Biol Cell 23: 3380–3390.

de Leeuw, R., Y. Gruenbaum and O. Medalia, 2018 Nuclear Lamins: thin filaments with major functions. Trends Cell Biol 28: 34–45.

De Magistris, P., and W. Antonin, 2018 The dynamic nature of the nuclear envelope. Curr Biol 28: R487–R497.

Dobrzynska, A., S. Gonzalo, C. Shanahan and P. Askjaer, 2016 The nuclear lamina in health and disease. Nucleus 7: 233–248.

Donnaloja, F., F. Carnevali, E. Jacchetti and M. T. Raimondi, 2020 Lamin A/C mechanotransduction in laminopathies. Cells 9.

dos Santos, Á., and C. P. Toseland, 2021 Regulation of nuclear mechanics and the Impact on DNA damage. Int J of Mol Sci 22: 3178.

Dujardin, D. L., and R. B. Vallee, 2002 Dynein at the cortex. Curr Opin Cell Biol 14: 44–49.

Echard, A., G. R. Hickson, E. Foley and P.H. O’Farrell, 2004 Terminal cytokinesis events uncovered after an RNAi screen. Curr Biol 14: 1685–1693.

Fraser, A. G., R. S. Kamath, P. Zipperlen, M. Martinez-Campos, M. Sohrmann et al., 2000 Functional genomic analysis of C. elegans chromosome I by systematic RNA interference. Nature 408: 325–330.

Galy, V., W. Antonin, A. Jaedicke, M. Sachse, R. Santarella et al., 2008 A role for gp210 in mitotic nuclear-envelope breakdown. J Cell Sci 121: 317–328.

Golden, A., J. Liu and O. Cohen-Fix, 2009 Inactivation of the C. elegans lipin homolog leads to ER disorganization and to defects in the breakdown and reassembly of the nuclear envelope. J Cell Sci 122: 1970–1978.

Goldman, R. D., D. K. Shumaker, M. R. Erdos, M. Eriksson, A. E. Goldman et al., 2004 Accumulation of mutant lamin A causes progressive changes in nuclear architecture in Hutchinson-Gilford progeria syndrome. Proc Natl Acad Sci U S A 101: 8963–8968.

Gonczy, P., C. Echeverri, K. Oegema, A. Coulson, S. J. Jones et al., 2000 Functional genomic analysis of cell division in C. elegans using RNAi of genes on chromosome III. Nature 408: 331–336.

Gonczy, P., H. Schnabel, T. Kaletta, A. D. Amores, T. Hyman et al., 1999 Dissection of cell division processes in the one cell stage Caenorhabditis elegans embryo by mutational analysis. J Cell Biol 144: 927–946.

Gorjánácz, M., and I. W. Mattaj, 2009 Lipin is required for efficient breakdown of the nuclear envelope in Caenorhabditis elegans. J Cell Sci 122: 1963–1969.

Green, R. A., H. L. Kao, A. Audhya, S. Arur, J. R. Mayers et al., 2011 A high-resolution C. elegans essential gene network based on phenotypic profiling of a complex tissue. Cell 145: 470–482.

Gruenbaum, Y., and R. Foisner, 2015 Lamins: nuclear intermediate filament proteins with fundamental functions in nuclear mechanics and genome regulation. Annu Rev Biochem 84: 131–164.

Hagstrom, K. A., V. F. Holmes, N. R. Cozzarelli and B. J. Meyer, 2002 C. elegans condensin promotes mitotic chromosome architecture, centromere organization, and sister chromatid segregation during mitosis and meiosis. Genes Dev 16: 729–742.

Haithcock, E., Y. Dayani, E. Neufeld, A. J. Zahand, N. Feinstein et al., 2005 Age-related changes of nuclear architecture in Caenorhabditis elegans. Proc Natl Acad Sci U S A 102: 16690–16695.

Han, G. S., W. I. Wu and G. M. Carman, 2006 The Saccharomyces cerevisiae Lipin homolog is a Mg2+-dependent phosphatidate phosphatase enzyme. J Biol Chem 281: 9210–9218.

Heald, R., and F. McKeon, 1990 Mutations of phosphorylation sites in lamin A that prevent nuclear lamina disassembly in mitosis. Cell 61: 579–589.

Horwich, A. L., W. A. Fenton, E. Chapman and G. W. Farr, 2007 Two families of chaperonin: physiology and mechanism. Annu Rev Cell Dev Biol 23: 115–145.

Joseph-Strauss, D., M. Gorjanacz, R. Santarella-Mellwig, E. Voronina, A. Audhya et al., 2012 Sm protein down-regulation leads to defects in nuclear pore complex disassembly and distribution in C. elegans embryos. Dev Biol 365: 445–457.

Karki, S., and E. L. Holzbaur, 1999 Cytoplasmic dynein and dynactin in cell division and intracellular transport. Curr Opin Cell Biol 11: 45–53.

Kotak, S., and P. Gonczy, 2013 Mechanisms of spindle positioning: cortical force generators in the limelight. Curr Opin Cell Biol 25: 741–748.

Laan, L., N. Pavin, J. Husson, G. Romet-Lemonne, M. Van Duijn et al., 2012 Cortical dynein controls microtubule dynamics to generate pulling forces that position microtubule asters. Cell 148: 502–514.

Laurell, E., K. Beck, K. Krupina, G. Theerthagiri, B. Bodenmiller et al., 2011 Phosphorylation of Nup98 by multiple kinases is crucial for NPC disassembly during mitotic entry. Cell 144: 539–550.

Lee, K. K., Y. Gruenbaum, P. Spann, J. Liu and K. L. Wilson, 2000 C. elegans nuclear envelope proteins emerin, MAN1, lamin, and nucleoporins reveal unique timing of nuclear envelope breakdown during mitosis. Mol Biol Cell 11: 3089–3099.

Lee, S. K., W. Li, S. E. Ryu, T. Rhim and J. Ahnn, 2010 Vacuolar (H+)-ATPases in Caenorhabditis elegans: what can we learn about giant H+ pumps from tiny worms? Biochim Biophys Acta 1797: 1687–1695.

Leroux, M. R., and E. P. M. Candido, 1995 Characterization of four new tcp-1-related cct genes from the nematode Caenorhabditis elegans. DNA Cell Biol 14: 951–960.

Leroux, M. R., and E. P. M. Candido, 1997 Subunit characterization of the Caenorhabditis elegans chaperonin containing TCP-1 and expression pattern of the gene encoding CCT-1. Biochem Biophys Res Comm 241: 687–692.

Linder, M. I., M. Kohler, P. Boersema, M. Weberruss, C. Wandke et al., 2017 Mitotic disassembly of nuclear pore complexes involves CDK1- and PLK1-mediated phosphorylation of key interconnecting nucleoporins. Dev Cell 43: 141–156.

Luckner, M., and G. Wanner, 2018 Precise and economic FIB/SEM for CLEM: with 2 nm voxels through mitosis. Histochem Cell Biol 150: 149–170.

Lundin, V. F., M. Srayko, A. A. Hyman and M. R. Leroux, 2008 Efficient chaperone-mediated tubulin biogenesis is essential for cell division and cell migration in C. elegans. Dev Biol 313: 320–334.

Markus, S. M., and W. L. Lee, 2011 Regulated offloading of cytoplasmic dynein from microtubule plus ends to the cortex. Dev Cell 20: 639–651.

Martino, L., S. Morchoisne-Bolhy, D. K. Cheerambathur, L. Van Hove, J. Dumont et al., 2017 Channel nucleoporins recruit PLK-1 to nuclear pore complexes to direct nuclear envelope breakdown in C. elegans. Dev Cell 43: 157–171 e157.

Mateus, D., E. S. Marini, C. Progida and O. Bakke, 2018 Rab7a modulates ER stress and ER morphology. Biochim Biophys Acta Mol Cell Res 1865: 781–793.

Matus, D. Q., X.-Y. Li, S. Durbin, D. Agarwal, Q. Chi et al., 2010 In vivo identification of regulators of cell invasion across basement membranes. Sci sig 3: ra35–ra35.

Mi, H., A. Muruganujan, X. Huang, D. Ebert, C. Mills et al., 2019 Protocol update for large-scale genome and gene function analysis with the PANTHER classification system (v.14.0). Nature protocols 14: 703–721.

Moore, J. K., J. Li and J. A. Cooper, 2008 Dynactin function in mitotic spindle positioning. Traffic 9: 510–527.

Moore, L. L., M. Morrison and M. B. Roth, 1999 HCP-1, a protein involved in chromosome segregation, is localized to the centromere of mitotic chromosomes in Caenorhabditis elegans. J Cell Biol 147: 471–480.

Mukherjee, R. N., P. Chen and D. L. Levy, 2016 Recent advances in understanding nuclear size and shape. Nucleus 7: 167–186.

Nelson, N., 1992 Structural conservation and functional diversity of V-ATPases. J Bioenerg Biomembr 24: 407–414.

Nguyen-Ngoc, T., K. Afshar and P. Gonczy, 2007 Coupling of cortical dynein and G alpha proteins mediates spindle positioning in Caenorhabditis elegans. Nat Cell Biol 9: 1294–1302.

Nishi, Y., E. Rogers, S. M. Robertson and R. Lin, 2008 Polo kinases regulate C. elegans embryonic polarity via binding to DYRK2-primed MEX-5 and MEX-6. Development 135: 687–697.

Noatynska, A., C. Panbianco and M. Gotta, 2010 SPAT-1/Bora acts with Polo-like kinase 1 to regulate PAR polarity and cell cycle progression. Development 137: 3315–3325.

Pathak, R. U., M. Soujanya and R. K. Mishra, 2021 Deterioration of nuclear morphology and architecture: A hallmark of senescence and aging. Ageing Res Rev 67: 101264.

Piano, F., A. J. Schetter, M. Mangone, L. Stein and K. J. Kemphues, 2000 RNAi analysis of genes expressed in the ovary of Caenorhabditis elegans. Curr Biol 10: 1619–1622.

Pintard, L., and B. Bowerman, 2019 Mitotic Cell Division in Caenorhabditis elegans. Genetics 211: 35–73.

Polinko, E. S., and S. Strome, 2000 Depletion of a Cks homolog in C. elegans embryos uncovers a post-metaphase role in both meiosis and mitosis. Curr Biol 10: 1471–1474.

Poteryaev, D., J. M. Squirrell, J. M. Campbell, J. G. White and A. Spang, 2005 Involvement of the actin cytoskeleton and homotypic membrane fusion in ER dynamics in Caenorhabditis elegans. Mol Biol Cell 16: 2139–2153.

Rahman, M., I. Y. Chang, A. Harned, R. Maheshwari, K. Amoateng et al., 2020 C. elegans pronuclei fuse after fertilization through a novel membrane structure. J Cell Biol 219.

Rahman, M. M., M. Munzig, K. Kaneshiro, B. Lee, S. Strome et al., 2015 Caenorhabditis elegans polo-like kinase PLK-1 is required for merging parental genomes into a single nucleus. Mol Biol Cell 26: 4718–4735.

Rahman, M. M., S. Rosu, D. Joseph-Strauss and O. Cohen-Fix, 2014 Down-regulation of tricarboxylic acid (TCA) cycle genes blocks progression through the first mitotic division in Caenorhabditis elegans embryos. Proc Natl Acad Sci U S A 111: 2602–2607.

Rivers, D. M., S. Moreno, M. Abraham and J. Ahringer, 2008 PAR proteins direct asymmetry of the cell cycle regulators Polo-like kinase and Cdc25. J Cell Biol 180: 877–885.

Romero-Bueno, R., P. de la Cruz Ruiz, M. Artal-Sanz, P. Askjaer and A. Dobrzynska, 2019 Nuclear Organization in Stress and Aging. Cells 8 664.

Santos-Rosa, H., J. Leung, N. Grimsey, S. Peak-Chew and S. Siniossoglou, 2005 The yeast lipin Smp2 couples phospholipid biosynthesis to nuclear membrane growth. EMBO J 24: 1931–1941.

Schindelin, J., I. Arganda-Carreras, E. Frise, V. Kaynig, M. Longair et al., 2012 Fiji: an open-source platform for biological-image analysis. Nat Methods 9: 676–682.

Shaw, S. L., E. Yeh, P. Maddox, E. D. Salmon and K. Bloom, 1997 Astral microtubule dynamics in yeast: a microtubule-based searching mechanism for spindle orientation and nuclear migration into the bud. J Cell Biol 139: 985–994.

Shibata, Y., T. Shemesh, W. A. Prinz, A. F. Palazzo, M. M. Kozlov et al., 2010 Mechanisms determining the morphology of the peripheral ER. Cell 143: 774–788.

Siniossoglou, S., H. Santos-Rosa, J. Rappsilber, M. Mann and E. Hurt, 1998 A novel complex of membrane proteins required for formation of a spherical nucleus. EMBO J 17: 6449–6464.

Skop, A. R., and J. G. White, 1998 The dynactin complex is required for cleavage plane specification in early Caenorhabditis elegans embryos. Curr Biol 8: 1110–1116.

Sonnichsen, B., L. B. Koski, A. Walsh, P. Marschall, B. Neumann et al., 2005 Full-genome RNAi profiling of early embryogenesis in Caenorhabditis elegans. Nature 434: 462–469.

Srayko, M., A. Kaya, J. Stamford and A. A. Hyman, 2005 Identification and characterization of factors required for microtubule growth and nucleation in the early C. elegans embryo. Dev Cell 9: 223–236.

Strome, S., J. Powers, M. Dunn, K. Reese, C. J. Malone et al., 2001 Spindle dynamics and the role of gamma-tubulin in early Caenorhabditis elegans embryos. Mol Biol Cell 12: 1751–1764.

Swan, K. A., A. F. Severson, J. C. Carter, P. R. Martin, H. Schnabel et al., 1998 cyk-1: a C. elegans FH gene required for a late step in embryonic cytokinesis. J Cell Sci 111: 2017–2027.

Tange, Y., A. Hirata and O. Niwa, 2002 An evolutionarily conserved fission yeast protein, Ned1, implicated in normal nuclear morphology and chromosome stability, interacts with Dis3, Pim1/RCC1 and an essential nucleoporin. J Cell Sci 115: 4375–4385.

Ungricht, R., and U. Kutay, 2017 Mechanisms and functions of nuclear envelope remodelling. Nat Rev Mol Cell Biol 18: 229–245.

Velez-Aguilera, G., S. Nkombo Nkoula, B. Ossareh-Nazari, J. Link, D. Paouneskou et al., 2020 PLK-1 promotes the merger of the parental genome into a single nucleus by triggering lamina disassembly. Elife 9 e59510.

Watson, M. L., 1955 The nuclear envelope; its structure and relation to cytoplasmic membranes. J Biophys Biochem Cytol 1: 257–270.

Worman, H. J., C. Ostlund and Y. Wang, 2010 Diseases of the nuclear envelope. Cold Spring Harb Perspect Biol 2: a000760.

Zink, D., A. H. Fischer and J. A. Nickerson, 2004 Nuclear structure in cancer cells. Nat Rev Cancer 4: 677–687.

Zipperlen, P., A. G. Fraser, R. S. Kamath, M. Martinez-Campos and J. Ahringer, 2001 Roles for 147 embryonic lethal genes on C.elegans chromosome I identified by RNA interference and video microscopy. EMBO J 20: 3984–3992.

