## Supplemental Tables S1 and S2 for "An RNAi screen for genes that affect nuclear morphology in *Caenorhabditis elegans* reveals the involvement of unexpected processes"

### Supplemental Table S1

| | Functional Category | ORF | Gene | Protein function[1, 2] | Phenotypes | | | | | | | | | Representative image(s) | Previously described nuclear phenotypes[3] |
|---|---|---|---|---|---|---|---|---|---|---|---|---|---|---|---|
| | | | | | Multi-nucleated | | | | Abnormal shape | | NPC distribution defect | | | | |
| | | | | | Paired Nuclei | One-cell stage | 2 or more cell stages | Micronuclei | Anaphase bridges | Deformed | Abnormal NE NPC distribution | Cytoplasmic NPCs | Intra-nuclear NPCs | | |
| | **mRNA processing** | | | | | | | | | | | | | | |
| | Splicing factors | Y116A8C.42 | *snr-1* | Orthologous to human small nuclear riboprotein Sm D3, involved in pre-mRNA splicing (Barbee *et al.* 2002). | | | | | | | √ | | | 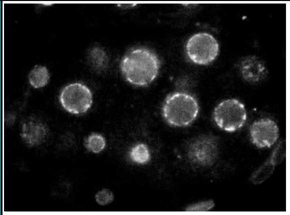 | NPC distribution defect *(Joseph-Strauss et al. 2012)*; nuclear appearance variant; multi-nucleated oocytes, enlarged nuclei (Green *et al.* 2011). |
| | | W08E3.1 | *snr-2* | Orthologous to human small nuclear riboprotein Sm B/B' involved in pre-mRNA splicing (Barbee *et al.* 2002). | | | | | | | √ | | | 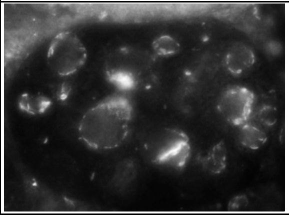 | NPC distribution defect *(Joseph-Strauss et al. 2012)*; nuclear appearance variant, small nuclei (Green *et al.* 2011). |
| | | C52E4.3 | *snr-4* | Orthologous to human small nuclear riboprotein Sm D2 involved in pre-mRNA splicing (Barbee *et al.* 2002). | | | | | | | √ | | | 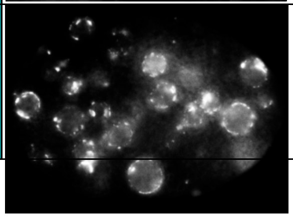 | NPC distribution defect *(Joseph-Strauss et al. 2012)*; nuclear appearance variant, small nuclei (Green *et al.* 2011); severe pleiotropic defects including multiple |

| | | | | | | | | | | | | | | | |
|---|---|---|---|---|---|---|---|---|---|---|---|---|---|---|---|
| | | | | | | | | | | | | | | | pronuclei in early embryos (Sonnichsen *et al.* 2005). |
| | | ZK652.1 | *snr-5* | Orthologous to human small nuclear riboprotein Sm F involved in pre-mRNA splicing (Barbee *et al.* 2002). | | | | | | | √ | | | 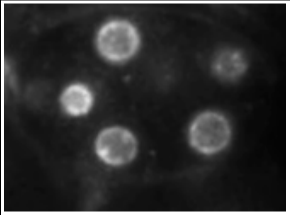 | NPC distribution defect *(Joseph-Strauss et al. 2012)*; nuclear appearance variant, small nuclei (Green *et al.* 2011). |
| | | Y49E10.15 | *snr-6* | Orthologous to human small nuclear riboprotein Sm E involved in pre-mRNA splicing (Barbee *et al.* 2002). | | | | | | | √ | | | 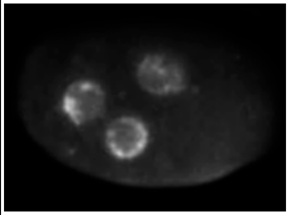 | NPC distribution defect *(Joseph-Strauss et al. 2012)*; nuclear appearance variant; small nuclei, nuclear morphology variation in early embryos (Green *et al.* 2011); severe pleiotropic defects including multiple pronuclei (Sonnichsen *et al.* 2005). |
| | | Y71F9B.4 | *snr-7* | Orthologous to human small nuclear riboprotein Sm G involved in pre-mRNA splicing (Barbee *et al.* 2002). | | | | | | | √ | | | 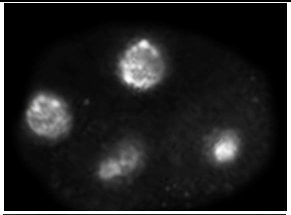 | NPC distribution defect *(Joseph-Strauss et al. 2012);* small nuclei (Green *et al.* 2011). |
| | | B0464.5 | *spk-1* | A serine/threonine kinase orthologous to mammalian serine/arginine-rich protein kinases (SRPKs) that phosphorylate SR proteins, required for splicing (Kuroyanagi *et al.* 2000). | √ | | √ | | | | | | | 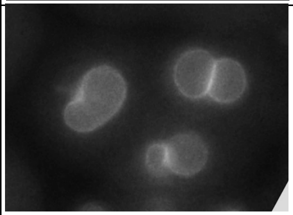 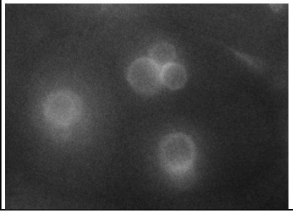 | |

| | | | | | | | | | | | | | | | |
|---|---|---|---|---|---|---|---|---|---|---|---|---|---|---|---|
| | | C41G7.1 | *smn-1* | Required for efficient splicing. Orthologous to the human SMN (Gao *et al.* 2014). | | | | | | | √ | | | 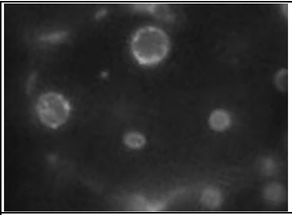 | |
| | | F58D5.1 | *hrp-2* | Orthologous to the mammalian heterogeneous nuclear ribonucleoproteins (hnRNP) Q and R, involved in alternative splicing (Kinnaird *et al.* 2004; Kabat *et al.* 2009). | | √ | | | | √ | | | | 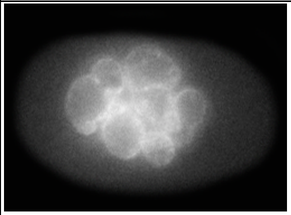 | |
| | | C04H5.6 | *mog-4* | A DEAH box protein, orthologous to the budding yeast *PRP16* gene that is required for splicing (Puoti and Kimble 2000). | | | | | | | √ | | | 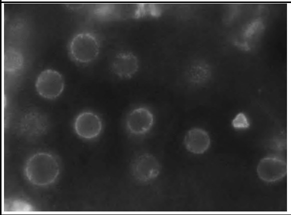 | |
| | | Y47G6A.20 | *rnp-6* | An RNP (RRM RNA binding domain) containing protein, orthologous to the Drosophila HALF-PINT splicing factor. RNP-6 has a role in alternative splicing (Barberan-Soler and Zahler 2008). | | | | | | √ | √ | | | 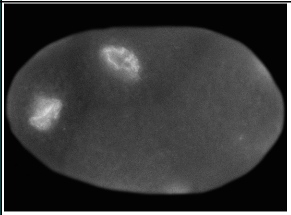 | |
| | mRNA deadenylation | Y56A3A.20 | *ccf-1* | A deadenylase. Orthologous to human CNOT-7 (Nousch *et al.* 2013). | | | √ | √ | | √ | | | | 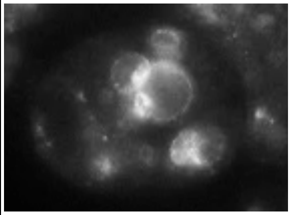 | Multi-nucleated cells (Lall *et al.* 2005); nuclear appearance variant, enlarged nuclei (Green *et al.* 2011). |

| | | | | | | | | | | | | | | | |
|---|---|---|---|---|---|---|---|---|---|---|---|---|---|---|---|
| | mRNA binding factors | F53B7.3 | *isy-1* | An ortholog of an mRNA binding protein involved in splicing (yeast) or micro-RNA processing (human) (Jiang *et al.* 2018). | | | | | | | √ | | | 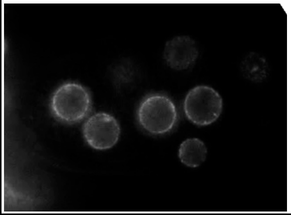 | |
| | | Y18D10A.17 | *car-1* | An ortholog of human LSM14A and LSM14B. CAR-1 functions with an RNA helicase CGH-1 and localizes to P bodies, where it likely represses translation (Audhya *et al.* 2005). | | | √ | | | | | | | 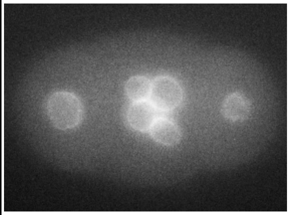 | Anaphase bridges (Audhya *et al.* 2005). |
| | **Translation** | | | | | | | | | | | | | | |
| | Ribosome subunits | T22F3.4 | *rpl-11.1* | A predicted 60S ribosomal protein L11 (Maciejowski *et al.* 2005). | | √ | | | | | | | | 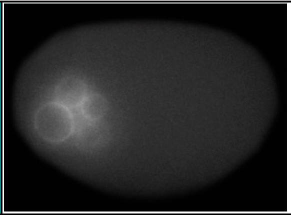 | Nuclear appearance variant (Waters *et al.* 2010); enlarged nuclei (Green *et al.* 2011); severe pleiotropic defects including multiple pronuclei in early embryos (Sonnichsen *et al.* 2005). |
| | | C14B9.7 | *rpl-21* | A predicted 60S ribosomal protein L21. | | | √ | √ | √ | | | | | 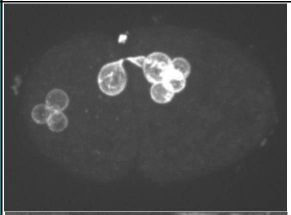 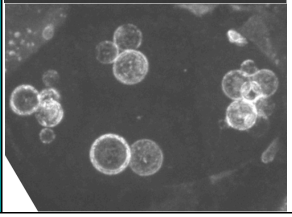 | Severe pleiotropic defects including multiple pronuclei in early embryos (Sonnichsen *et al.* 2005). |

| | | | | | | | | | | | | | | | |
|---|---|---|---|---|---|---|---|---|---|---|---|---|---|---|---|
| | | C27A2.2 | *rpl-22* | A predicted 60S ribosomal protein L22. | | | √ | | | | | | | 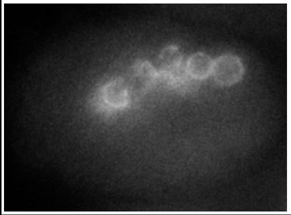 | Chromosome segregation defects (Sonnichsen *et al.* 2005). |
| | | B0336.10 | *rpl-23* | A predicted 60S ribosomal protein L23. | | | √ | √ | √ | | | | | 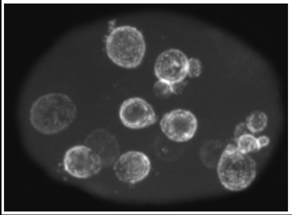 | Severe pleiotropic defects including multiple pronuclei in early embryos (Sonnichsen *et al.* 2005); enlarged nuclei (Green *et al.* 2011). |
| | | F52B5.6 | *rpl-25.2* | A predicted 60S ribosomal protein L25a. | √ | | | | | √ | | | | 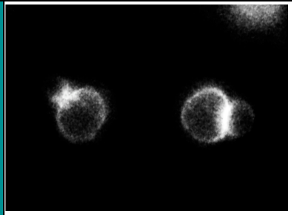 | Severe pleiotropic defects including multiple pronuclei in early embryos (Sonnichsen *et al.* 2005). |
| | | F10E7.7 | *rpl-33* | A predicted 60S ribosomal protein L35a. | √ | | √ | √ | √ | √ | | | | 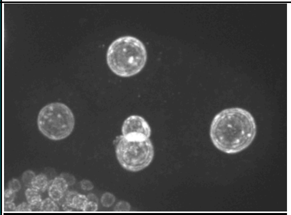 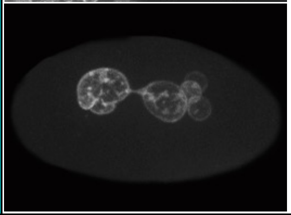 | Severe pleiotropic defects including multiple pronuclei in early embryos (Sonnichsen *et al.* 2005). |

| | | | | | | | | | | | | | | | |
|---|---|---|---|---|---|---|---|---|---|---|---|---|---|---|---|
| | | F40F8.10 | *rps-9* | A predicted 40S ribosomal protein S9. | | | | | √ | | | | | 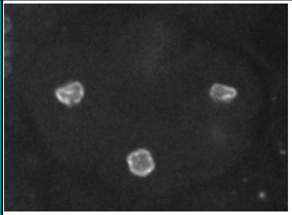 | Severe pleiotropic defects including multiple pronuclei in early embryos (Sonnichsen *et al.* 2005); small nuclei (Green *et al.* 2011). |
| | | F54E7.2 | *rps-12* | A predicted 40S ribosomal protein S12. | | | √ | √ | | √ | | | | 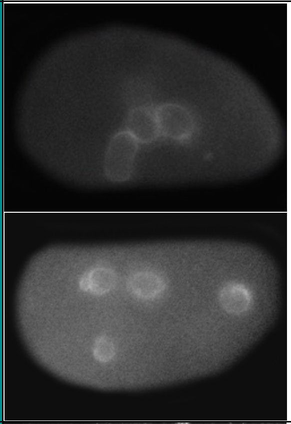 | Severe pleiotropic defects including multiple pronuclei in early embryos (Sonnichsen *et al.* 2005); small nuclei (Green *et al.* 2011). |
| | | F37C12.11 | *rps-21* | A predicted 40S ribosomal protein S21. | | | √ | √ | √ | | | | | 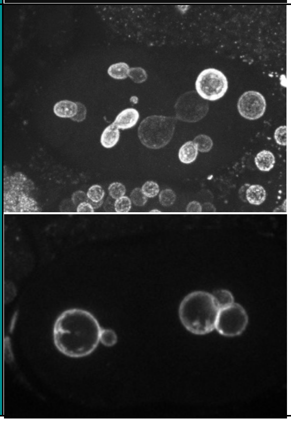 | Severe pleiotropic defects including multiple pronuclei in early embryos (Sonnichsen *et al.* 2005); small nuclei (Green *et al.* 2011). |

| | | | | | | | | | | | | | | | |
|---|---|---|---|---|---|---|---|---|---|---|---|---|---|---|---|
| | | F53A3.3 | *rps-22* | A predicted 40S ribosomal protein S22. | √ | | √ | √ | | √ | | | | 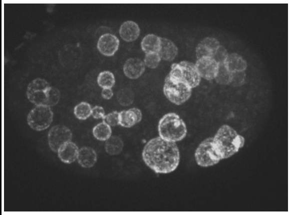 | Severe pleiotropic defects including multiple pronuclei in early embryos (Sonnichsen *et al.* 2005). |
| | tRNA aminoacylation | B0464.1 | *dars-1* | An ortholog of human DARS, an aspartyl(D) tRNA synthetase. | | | √ | | | √ | | | | 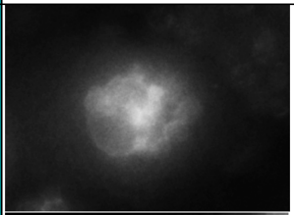 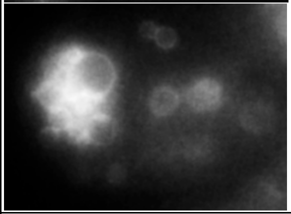 | Severe pleiotropic defects including multiple pronuclei in early embryos (Sonnichsen *et al.* 2005); nuclear appearance variant, small nuclei (Green *et al.* 2011). |
| | | C47E12.1 | *sars-2* | An ortholog of human SARS2, a seryl amino-acyl tRNA synthetase. | | | √ | | | | | | | 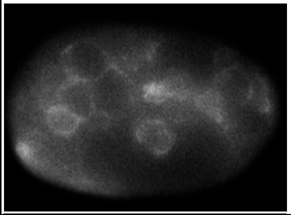 | |
| | | F26F4.10 | *rars-1* | An ortholog of human RARS, an—arginyl-tRNA synthetase. | | | | √ | | | | | | 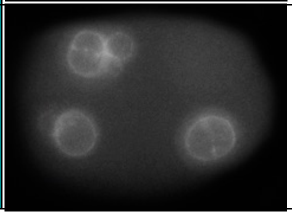 | Severe pleiotropic defects including multiple pronuclei in early embryos (Sonnichsen *et al.* 2005); enlarged nuclei (Green *et al.* 2011). |

| | | | | | | | | | | | | | | | |
|---|---|---|---|---|---|---|---|---|---|---|---|---|---|---|---|
| | | Y87G2A.5 | *glp-4* | An ortholog of human VARS, a valyl-tRNA synthetase (Rastogi *et al.* 2015). | √ | | √ | √ | | √ | | | | 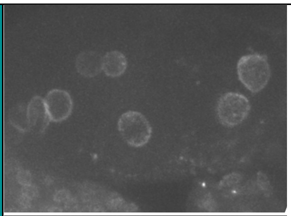 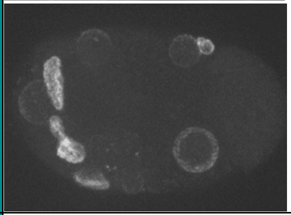 | Severe pleiotropic defects including multiple pronuclei in early embryos (Sonnichsen *et al.* 2005). |
| | | T20H4.3 | *pars-1* | An ortholog of human EPRS, a glutamyl-prolyl-tRNA synthetase. | √ | | √ | | | √ | | | | 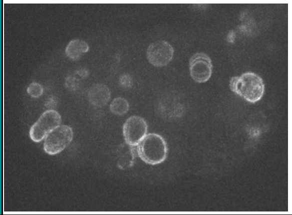 | Severe pleiotropic defects including multiple pronuclei in early embryos (Sonnichsen *et al.* 2005). |
| | | T11G6.1 | *hars-1* | An ortholog of human HARS, a histidyl-tRNA synthetase. | √ | | √ | | | | | | | 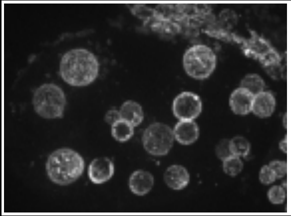 | Sister chromatid separation defective causing deformed nuclei (Sonnichsen *et al.* 2005). |
| | | F28H1.3 | *aars-2* | An ortholog of human AARS, an alanyl-tRNA synthetase. | | | | | | √ | √ | | | 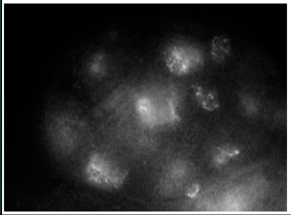 | Severe pleiotropic defects including multiple pronuclei in early embryos (Sonnichsen *et al.* 2005). |

| | | | | | | | | | | | | | | | |
|---|---|---|---|---|---|---|---|---|---|---|---|---|---|---|---|
| | Translation factors | R08D7.3 | *eif-3.D* | An ortholog of human EIF3D, a translation initiation factor (Rhoads *et al.* 2006). | | | √ | | | | | | | 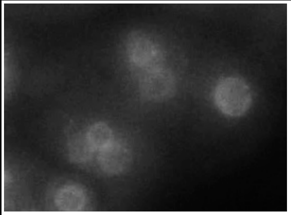 | Severe pleiotropic defects including multiple pronuclei in early embryos (Sonnichsen *et al.* 2005); nuclear appearance variant (Green *et al.* 2011). |
| | | C37C3.2 | *phi-18* | An ortholog of human EIF5, a eukaryotic translation initiation factor 5 (Rhoads *et al.* 2006). | | | √ | | | √ | | | | 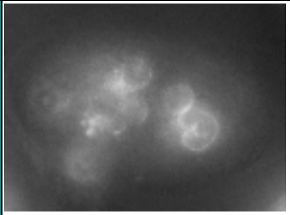 | Severe pleiotropic defects including multiple pronuclei in early embryos (Sonnichsen *et al.* 2005). |
| | | Y47H9C.7 | *eif-2 β* | An ortholog of human EIF2B2, a translation initiation factor (Rhoads *et al.* 2006). | √ | | | | | | | | | 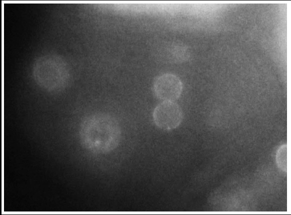 | Severe pleiotropic defects including multiple pronuclei in early embryos (Sonnichsen *et al.* 2005). |
| | | Y39E4B.1 | *abce-1* | An ortholog of human ABCE1 which is involved in translation initiation and ribosome recycling (Zhao *et al.* 2004). | √ | | √ | | | √ | | | | 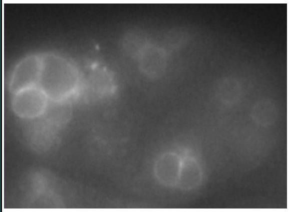 | |

| | | | | | | | | | | | | | | | |
|---|---|---|---|---|---|---|---|---|---|---|---|---|---|---|---|
| | | R03G5.1 | *eef-1A.2* | An ortholog of human EEF1A2, a translation elongation factor (Rhoads *et al.* 2006). | | | | | √ | √ | | | | 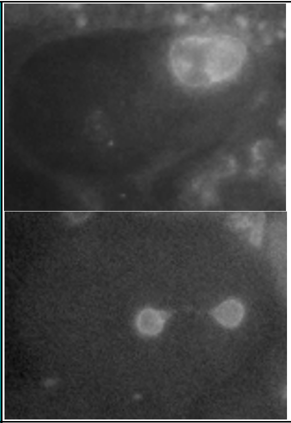 | |
| | | F25H5.4 | *eef-2* | An ortholog of human EEF2, a translation elongation factor (Ofulue and Candido 1991; Rhoads *et al.* 2006). | | | √ | | | √ | | | | 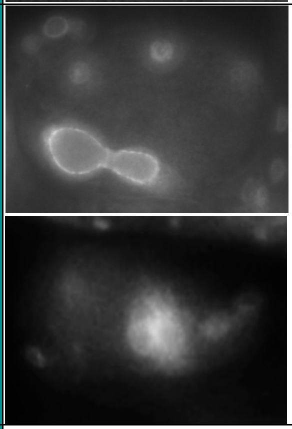 | Nuclear appearance variant (Green *et al.* 2011). |
| | Ribosome biogenesis | T19A6.2 | *ngp-1* | A nucleolar GTP-binding protein orthologous to human GNL2. NGP-1 has a role in non-sense mediated decay. Yeast homolog, Nog2, is involved in ribosome biogenesis (Casadio *et al.* 2015). | | | √ | | | | | | | 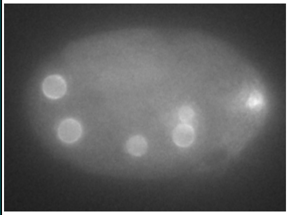 | |

| | | | | | | | | | | | | | | |
|---|---|---|---|---|---|---|---|---|---|---|---|---|---|---|
| | | Y48G1A.4 | *nol-14* | A nucleolar protein, an ortholog of human NOP14, required for ribosome biogenesis. | | | | | | √ | | | | 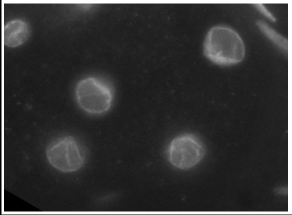 |
| | **Microtubule function and centrosomes** | | | | | | | | | | | | | |
| | Tubulin | C47B2.3 | *tba-2* | α-tubulin, forms microtubules along with β-tubulin (Phillips *et al.* 2004). | | | √ | | | √ | | √ | | 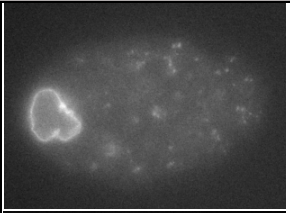 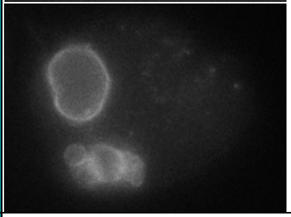 |
| | | F44F4.11 | *tba-4* | α-tubulin, forms microtubules along with β-tubulin. | | | √ | | | √ | | | | 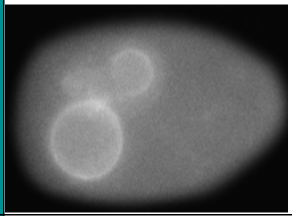 |
| | | C36E8.5 | *tbb-2* | β-tubulin, dimerizes with α-tubulin to form microtubules (Wright and Hunter 2003; Ellis *et al.* 2004). | | | √ | | | √ | | | | 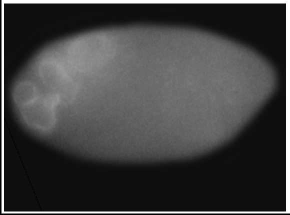 |

| | | | | | | | | | | | | | | | |
|---|---|---|---|---|---|---|---|---|---|---|---|---|---|---|---|
| | Motor proteins | M03D4.1 | *zen-4* | A kinesin motor protein belongs to the MKLP/CHO1 subfamily of kinesin. ZEN-4 bundles antiparallel microtubules at the spindle midzone (Raich *et al.* 1998; Mishima *et al.* 2002; Siddiqui 2002). | | | √ | | | | | | | 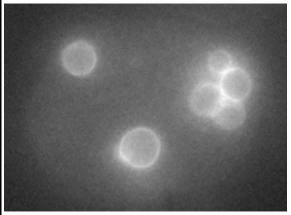 | Multi-nucleated cells (Raich *et al.* 1998; Echard *et al.* 2004). |
| | | C06G3.2 | *klp-18* | A kinesin motor protein belongs to the MCAK/KIF2 subfamily of kinesins. Contributes to acentrosomal spindle bipolarity (Siddiqui 2002; Segbert *et al.* 2003; Wolff *et al.* 2016). | √ | | | √ | | √ | | | | 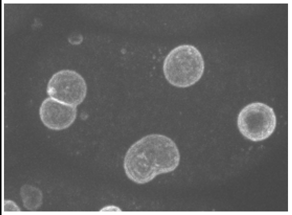 | Multi-nucleated cells (Piano *et al.* 2002). |
| | | Y43F4B.6 | *klp-19* | A kinesin motor protein belongs to the chromokinesin subfamily of kinesins that is associated with chromosomes (Siddiqui 2002; Powers *et al.* 2004). | | | | √ | √ | | | | | 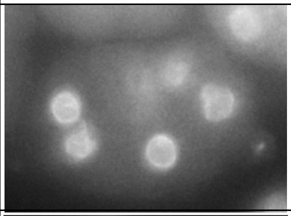 | Anaphase bridges (Powers *et al.* 2004); chromosome segregation defect (Sonnichsen *et al.* 2005). |
| | | T26A5.9 | *dlc-1* | A light chain subunit of dynein, which is a minus-end direct motor. Orthologous to human DYNLL1 and DYNLL2 (O'rourke *et al.* 2007). | | | √ | √ | | √ | | | | 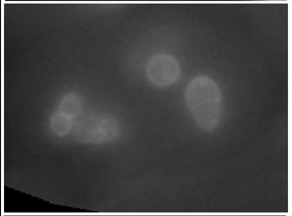 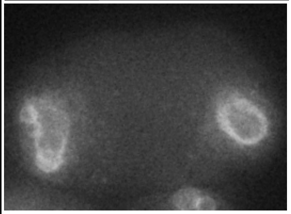 | Multi-nucleated oocytes (Green *et al.* 2011); nuclear appearance variant, enlarged nuclei (Dorsett and Schedl 2009). |

| | | | | | | | | | | | | | | | |
|---|---|---|---|---|---|---|---|---|---|---|---|---|---|---|---|
| | | C17H12.1 | *dyci-1* | A putative intermediate chain subunit of dynein, a minus-end directed motor (O'rourke *et al.* 2007). | √ | | | | | √ | | | | 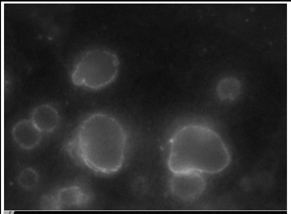 | |
| | | C39E9.14 | *dli-1* | A light intermediate chain subunit of dynein, a microtubule minus-directed motor (Yoder and Han 2001). | | | √ | | | √ | | | | 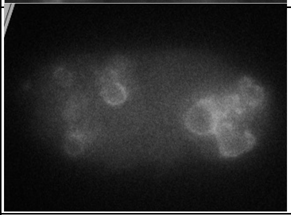 | Multi-nucleated cells (Yoder and Han 2001). |
| | | F53A2.4 | *nud-1* | An ortholog of *Aspergillus* nuclear division protein nudC. NUD-1 associates with dynein (Dawe *et al.* 2001). | | | √ | | | √ | | | | 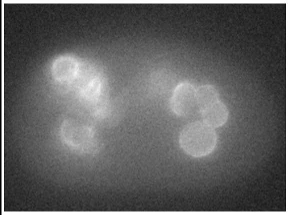 | Multi-nucleated cells (Aumais *et al.* 2003); nuclear appearance variant, small nuclei (Green *et al.* 2011). |
| | | ZK593.5 | *dnc-1* | A subunit of dynactin, a dynein associated protein required for cargo binding. *dnc-1* is an ortholog of p150/GLUED/DCTN1 (Skop and White 1998). | | | √ | √ | | √ | | | | 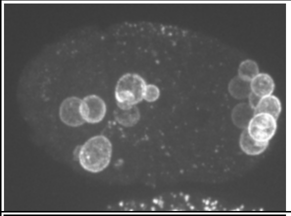 | Chromosome segregation defect (Skop and White 1998). |
| | | Y53F4B.22 | *arp-1* | Actin related protein A, orthologous to human ACTR1. ARP-1 is a component of the dynactin complex (Terasawa *et al.* 2010). | | | √ | √ | | √ | | | | 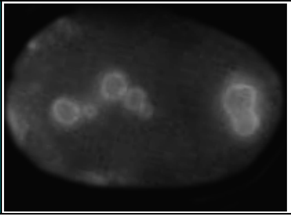 | |

| | | | | | | | | | | | | | | |
|---|---|---|---|---|---|---|---|---|---|---|---|---|---|---|
| | | C49H3.8 | *arp-11* | An actin related protein 10, orthologous to human ACTR10, a component of dynactin complex (Terasawa *et al.* 2010). | | | √ | | | | | | | 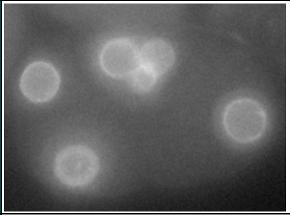 |
| | Tubulin/microtubule-interacting proteins | C05D11.3 | *txdc-9* | A thioredoxin domain-containing protein, orthologous to human TXNDC9. TXDC-9 is required for tubulin acetylation (Ogawa *et al.* 2004; Chen *et al.* 2015). | | | √ | √ | | √ | | | | 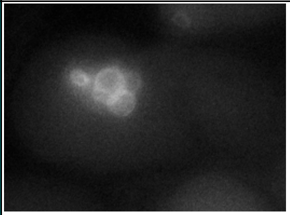 |
| | | T06G6.9 | *pfd-3* | Predicted to be a subunit of prefoldin. PFD-3 is involved in tubulin folding (Lundin *et al.* 2008). | | | √ | | | √ | | | | 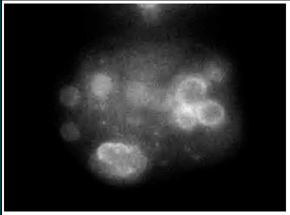 |
| | | F22B5.1 | *evl-20* | A small GTPase, an ortholog of human ARL2. May regulate microtubule structure (Antoshechkin and Han 2002). | √ | | √ | | √ | √ | | | | 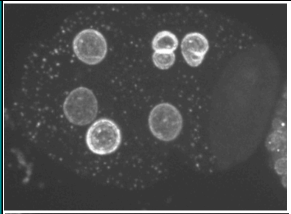 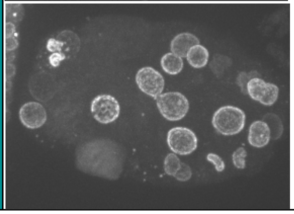 |

| | | | | | | | | | | | | | | |
|---|---|---|---|---|---|---|---|---|---|---|---|---|---|---|
| | positioning | C38C10.4 | *gpr-2* | A G-protein regulator-regulates the GTPase activity of Gα protein GOA-1. Involved directly in regulating spindle positioning (Gonczy *et al.* 2000; Colombo *et al.* 2003; Srinivasan *et al.* 2003). | √ | | √ | | | √ | | | | 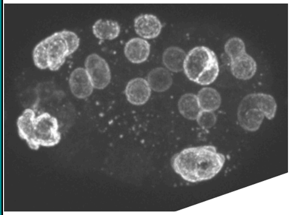 | Multi-nucleated cells (Piano *et al.* 2002). |
| | Meiotic spindle | C28C12.2 | *mesp-1* | MESP-1 interacts with KLP-18 for maintaining acentrosomal spindle bipolarity (Wignall and Villeneuve 2009; Wolff *et al.* 2016). | | | √ | | | | | | | 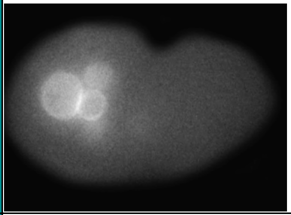 | Multi-nucleated cells (Piano *et al.* 2002). |
| | | F57B10.12 | *mei-2* | A subunit of the katanin complex (along with MEI-1) which has microtubule severing activity. MEI-2 is required for the establishment of the oocyte meiotic spindle (Srayko *et al.* 2000). | √ | | √ | √ | | | | | | 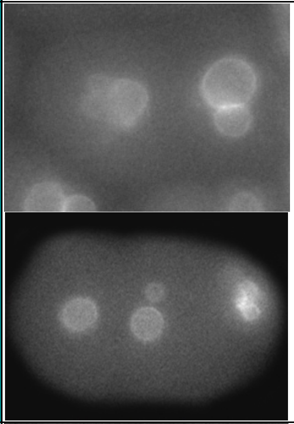 | |
| | Centrosome-associated proteins | C45G3.5 | *gip-2* | A γ-tubulin interacting protein, a component of the γ-tubulin complex (Hannak *et al.* 2002). | | | √ | | √ | | | | | 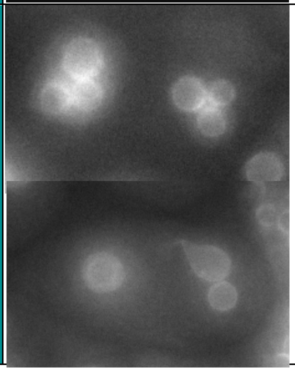 | |

| | | | | | | | | | | | | | | | |
|---|---|---|---|---|---|---|---|---|---|---|---|---|---|---|---|
| | | F58A4.8 | *tbg-1* | γ-tubulin, a centrosome component (Bobinnec *et al.* 2000). | | | √ | √ | | √ | | | | 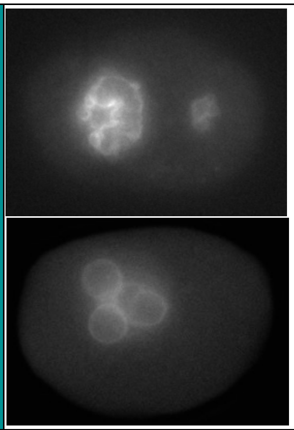 | Multi-nucleated oocyte, nuclear appearance variant; small nuclei (Green *et al.* 2011); multi-nucleated cells (Piano *et al.* 2002). |
| | | F56A3.4 | *spd-5* | A coiled coil, pericentriolar material (PCM) scaffold protein (Hamil *et al.* 2002). | | | √ | √ | √ | √ | | | | 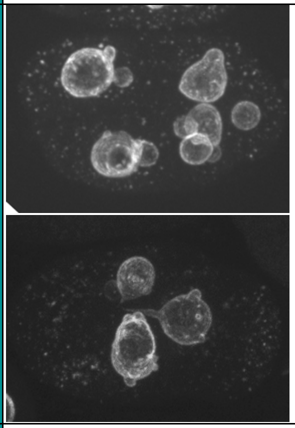 | Multi-nucleated cells (Piano *et al.* 2002). |
| | | F35B12.5 | *sas-5* | A coiled-coil protein that localizes to centrioles and is required for centriole duplication (Delattre *et al.* 2004; Rogala *et al.* 2015). | | | √ | | | √ | | | | 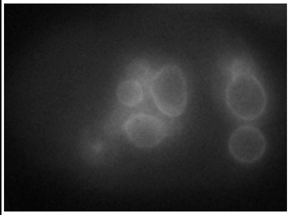 | Multi-nucleated cells; nuclear morphology variation in early embryos (Schmutz and Spang 2005). |

| | | | | | | | | | | | | | | | |
|---|---|---|---|---|---|---|---|---|---|---|---|---|---|---|---|
| | | Y45F10D.9 | *sas-6* | A coiled-coil protein that localizes to centrioles and is required for centriole duplication (Dammermann *et al.* 2004; Leidel *et al.* 2005; Nakazawa *et al.* 2007). | | | √ | | | √ | | | | 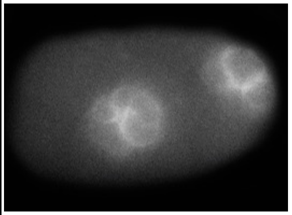 | |
| | | C30B5.1 | *szy-4* | Protein of unknown function. *szy-4 (loss-of-function)* mutation rescues the centrosome duplication failure caused by *zyg-1(it25)* loss-of-function mutants (Kemp *et al.* 2007). | | | √ | √ | | √ | | | | 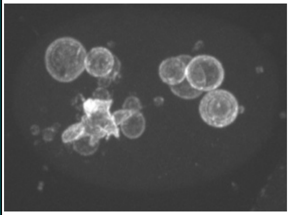 | Multi-nucleated oocytes (Green *et al.* 2011). |
| | **Cell Cycle** | | | | | | | | | | | | | | |
| | Cell cycle regulators | T06E6.2 | *cyb-3* | A B-type cyclin that activates CDK-1 (Sonneville and Gonczy 2004; Van Der Voet *et al.* 2009). | √ | | √ | √ | | √ | √ | √ | | 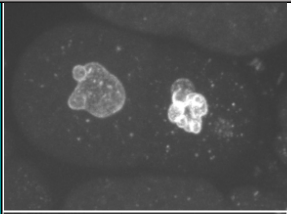 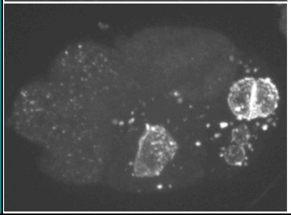 | Multi-nucleated cells, nuclear appearance variant, small nuclei (Green *et al.* 2011). |
| | | Y53C12A.1 | *wee-1.3* | Orthologous to the vertebrate Myt1, belongs to the Wee1 family of kinases. WEE-1.3 represses the activity of Cdk-1 (Lamitina and L'hernault 2002; Burrows *et al.* 2006). | | | | | | | | √ | | 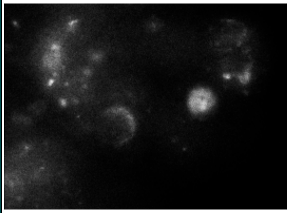 | Multi-nucleated oocytes, nuclear appearance variant, enlarged nuclei (Green *et al.* 2011). |

| | | | | | | | | | | | | | | |
|---|---|---|---|---|---|---|---|---|---|---|---|---|---|---|
| | | C09G4.3 | *cks-1* | A cyclin dependent kinase regulatory subunit, an ortholog of human Cks/Suc (Polinko and Strome 2000). | √ | | √ | √ | √ | √ | √ | √ | | 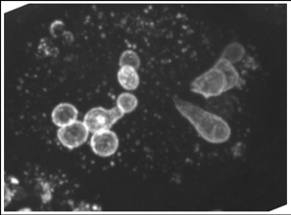 | Pronuclear morphology defect (Polinko and Strome 2000). |
| | | F26E4.1 | *sur-6* | A regulatory subunit of serine/threonine protein phosphatase 2A (PP2A-B). Among the processes regulated by SUR-6 are Ras-mediated signaling, membrane trafficking and centriole duplication (Sieburth *et al.* 1999; Jiu *et al.* 2014). | | | | | √ | | | | | 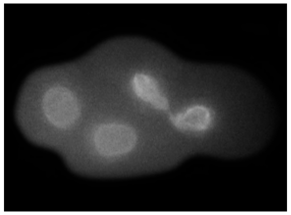 | Anaphase bridges (Song *et al.* 2011); multi-nucleated cells (Fraser *et al.* 2000; Zipperlen *et al.* 2001). |
| | | T09A5.9 | *sds-22* | Ortholog of human protein phosphatase 1 regulatory subunit. Yeast and human orthologs are involved in processes related to mitosis. SDS-22 is required for centriole duplication (Peel et al. 2017). | | | √ | | | | √ | | | 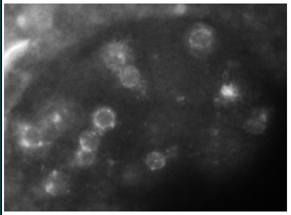 | |
| | | K07C11.2 | *air-1* | An Aurora A serine/threonine kinase. Regulates chromosome segregation and cell division related processes (Schumacher et al. 1998; Hannak et al. 2001). | | | | √ | | | | | | 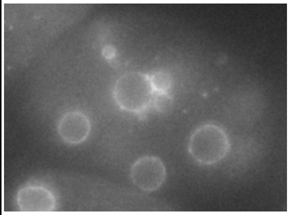 | Nuclear appearance variant, enlarged nuclei (Green *et al.* 2011); multi-nucleated cells (Piano *et al.* 2002; Echard *et al.* 2004); anaphase bridges (Schumacher *et al.* 1998). |
| | | C27A2.3 | *ify-1* | A securin like protein, that is degraded at the onset of anaphase following ubiquitination by the anaphase promoting complex-dependent. Inhibits the activity of separase, | | | √ | √ | | √ | | | | 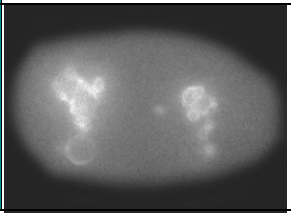 | Pronuclear size defect (Kitagawa *et al.* 2002). |

| | | | | | | | | | | | | | | |
|---|---|---|---|---|---|---|---|---|---|---|---|---|---|---|
| | | | | SEP-1, that is needed for sister chromatid separation (Kitagawa et al. 2002). | | | | | | | | | | | |
| | Anaphase promoting complex | ZK177.6 | *fzy-1* | A Cdc20/Fizzy homolog, a substrate recognition subunit of the anaphase promoting complex (Kitagawa et al. 2002). | | | √ | | | √ | | | | 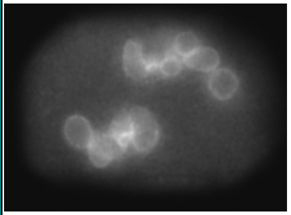 | |
| | | F35G12.9 | *apc-11* | A subunit of anaphase promoting complex, an E3 ubiquitin ligase. Required for anaphase initiation by promoting the ubiquitination of Cyclin B and other proteins (Davis et al. 2002). | | √ | | √ | | √ | | | | 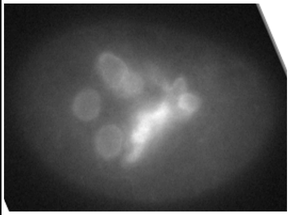 | |
| | DNA replication | F58A4.4 | *pri-1* | An ortholog of DNA polymerase $\alpha$-primase subunit D (Encalada et al. 2000). | | | | | √ | | | | | 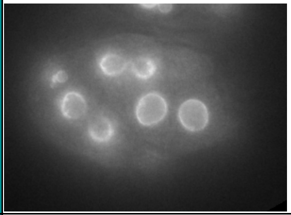 | Multi-nucleated cells (Piano *et al.* 2002). |
| | | W02D9.1 | *pri-2* | An ortholog of DNA polymerase $\alpha$-primase subunit C (Encalada et al. 2000). | | | | √ | √ | | | | | 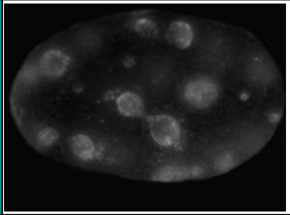 | Multi-nucleated cells (Piano *et al.* 2002); nuclear morphology variation in early embryos (Fraser *et al.* 2000; Zipperlen *et al.* 2001). |
| | | ZC168.3 | *orc-5* | An ortholog of human Orc5, required for the initiation of DNA replication. | | | √ | √ | √ | √ | | | | 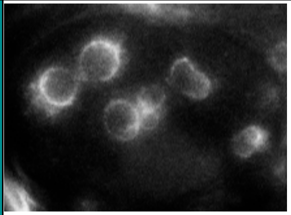 | |

| | | | | | | | | | | | | | | | |
|---|---|---|---|---|---|---|---|---|---|---|---|---|---|---|---|
| | | F32D1.10 | *mcm-7* | An ortholog of mammalian MCM7, part of a helicase complex involved in initiation of DNA replication (Woodward et al. 2006; Sonneville et al. 2012). | √ | | √ | √ | √ | √ | | √ | | 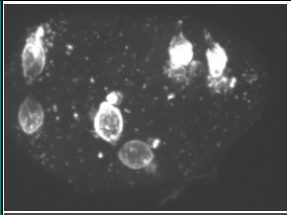 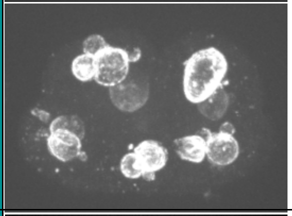 | Multi-nucleated cells (Piano *et al.* 2002). |
| | Ribonucleotide reductase | T23G5.1 | *rnr-1* | A putative large subunit of ribonucleotide reductase involved in deoxyribonucleotide biosynthesis (Hong et al. 1998; Mori et al. 2008). | | | | √ | | √ | | | | 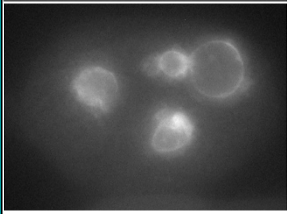 | |
| | | C03C10.3 | *rnr-2* | A putative small subunit of ribonucleotide reductase, involved in deoxyribonucleotide biosynthesis (Mori et al. 2008). | √ | | √ | | | | | | | 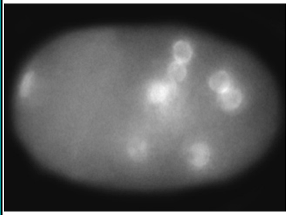 | Enlarged nuclei (Green *et al.* 2011). |
| | Kinetochore proteins | W01B6.9 | *ndc-80* | An ortholog of the yeast kinetochore protein Ndc80. NDC-80 is required for kinetochore-microtubule interactions (Desai et al. 2003). | | | √ | | √ | √ | | | | 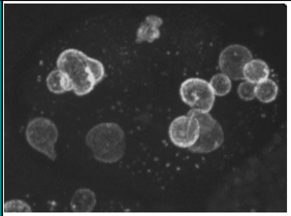 | Pronuclear size defect; multi-nucleated cells (Piano *et al.* 2002). |

| | | | | | | | | | | | | | | | |
|---|---|---|---|---|---|---|---|---|---|---|---|---|---|---|---|
| | | C06A8.5 | *spdl-1* | Homologous to the *Drosophila* coiled coil protein Spindly, which is responsible for recruiting Dynein to kinetochores. SPDL-1 is also required for spindle assembly checkpoint function (Gassmann *et al.* 2008; Yamamoto *et al.* 2008). | | | | | √ | | | | | 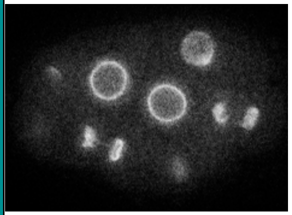 | |
| | | F58A4.3 | *hcp-3* | A Histone H3-like centromeric protein, orthologous to human CENP-A (Buchwitz *et al.* 1999; Oegema *et al.* 2001). | √ | | | | √ | √ | | | | 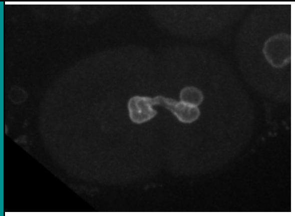 | Nuclear appearance variant (Green *et al.* 2011). |
| | Spindle assembly checkpoint | R06C7.8 | *bub-1* | A serine/threonine kinase orthologous to the yeast and human BUB1. BUB-1 is required for spindle assembly checkpoint signaling (Oegema *et al.* 2001). | √ | | | √ | √ | | | | | 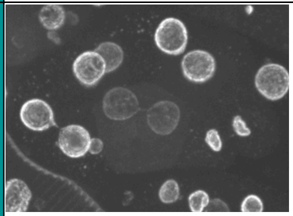 | Multi-nucleated cells (Fraser *et al.* 2000; Dawe *et al.* 2001; Zipperlen *et al.* 2001). |
| | Cytokinesis | K08E3.6 | *cyk-4* | A Rho GTPase-activating protein (GAP) protein. CYK-4 is part of the central spindle complex that is required for central spindle formation and cytokinesis (Jantsch-Plunger *et al.* 2000). | | √ | | | | √ | | | | 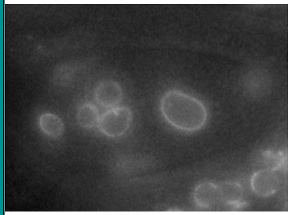 | Nuclear appearance variant, small nuclei (Green *et al.* 2011); multi-nucleated cells (Encalada *et al.* 2000; Piano *et al.* 2002; Green *et al.* 2011). |
| | **Signaling** | | | | | | | | | | | | | | |

| | | | | | | | | | | | | | | | |
|---|---|---|---|---|---|---|---|---|---|---|---|---|---|---|---|
| | MAP kinase signaling | F49E11.1 | *mbk-2* | A Serine/threonine kinase, it is a member of the DYRK (dual-specificity Yak1-related kinase) family of proteins (Pellettieri *et al.* 2003; Raich *et al.* 2003). | √ | | √ | | √ | √ | | | | 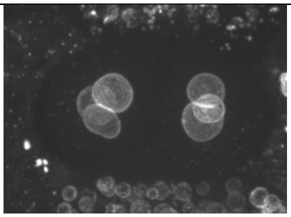 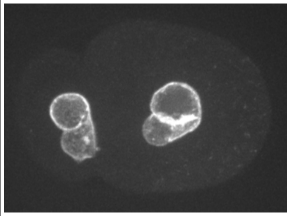 | Anaphase bridges (Pellettieri *et al.* 2003). |
| | | ZK792.6 | *let-60* | A GTP-binding RAS protein. LET-60 activity regulates RAS-mediated signaling as part of the part of the MAP kinase cascade (Han and Sternberg 1990). | √ | | √ | | | | | | | 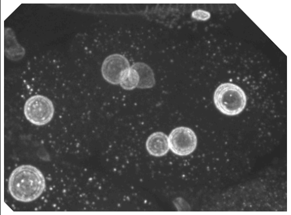 | Enlarged nuclei (Green *et al.* 2011). |
| | Rho GTPase | T19E10.1 | *ect-2* | A Rho guanine exchange factor (GEF) required for cytokinesis and cell polarity in early embryos (Morita *et al.* 2005; Motegi and Sugimoto 2006). | | | √ | | | √ | | | | 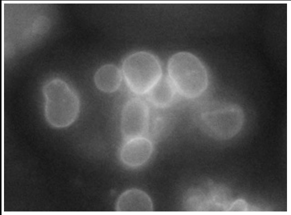 | Multi-nucleated cells (Echard *et al.* 2004). |

| | | | | | | | | | | | | | | | |
|---|---|---|---|---|---|---|---|---|---|---|---|---|---|---|---|
| | Wnt pathway related | C03C10.1 | *kin-19* | A Casein kinase I homolog. KIN-19 is a serine/threonine kinase In the Wnt signaling pathway (Walston *et al.* 2004; Banerjee *et al.* 2010). | √ | | √ | | √ | | | | | 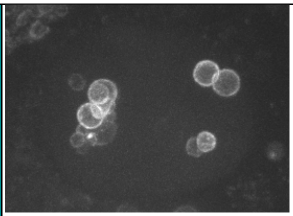 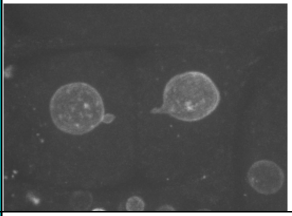 | Small nuclei (Green *et al.* 2011). |
| | | Y18D10A.5 | *gsk-3* | An ortholog of human GSK3B, a kinase involved in Wnt signaling (Schlesinger *et al.* 1999). | | | | | | √ | | | | 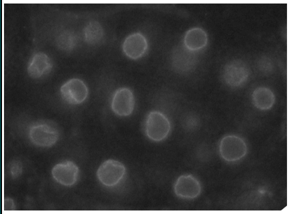 | |
| | **Protein degradation** | | | | | | | | | | | | | | |
| | Proteasome | C30C11.2 | *rpn-3* | A putative 19S non-ATPase subunit of the proteasome complex. Orthologous to human PSMD3 (Davy *et al.* 2001). | | | | | | √ | | | | 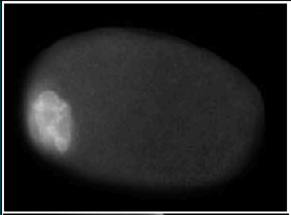 | Nuclear appearance variant; enlarged nuclei (Green *et al.* 2011). |
| | | C52E4.4 | *rpt-1* | A putative 19S ATPase subunit of proteasome complex. Orthologous to human PSMC2 (Davy *et al.* 2001). | | √ | | √ | | √ | | | | 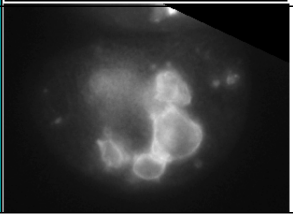 | Nuclear appearance variant, enlarged nuclei (Green *et al.* 2011). |

| | | | | | | | | | | | | | | | |
|---|---|---|---|---|---|---|---|---|---|---|---|---|---|---|---|
| | | Y110A7A.14 | *pas-3* | A putative 20S α-type proteasome subunit. Orthologous to human PSMA4 (Davy *et al.* 2001). | | | | | | √ | | √ | | 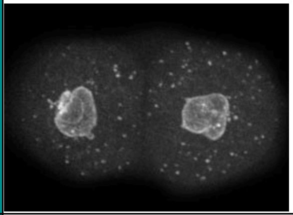 | Nuclear appearance variant (Green *et al.* 2011). |
| | Ubiquitylation | M7.1 | *let-70* | A ubiquitin conjugating enzyme, also known as UBC-2. Works with the anaphase promoting complex (Frazier *et al.* 2004). | | | | √ | | √ | | | | 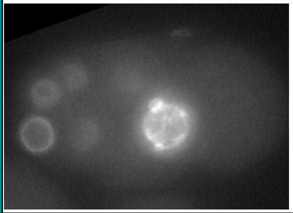 | Nuclear appearance variant (Green *et al.* 2011). |
| | | ZK858.4 | *mel-26* | An ortholog of human SPOP. MEL-26 is the substrate recognition subunit of the CUL-3 ubiquitin ligase. Promotes MEI-1 degradation, thereby facilitating the transition from the meiotic spindle the mitotic one (Dow and Mains 1998; Pintard *et al.* 2003b). | √ | | √ | | | √ | | | | 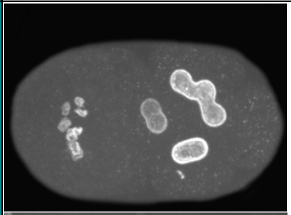 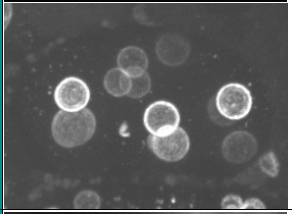 | |
| | | ZK520.4 | *cul-2* | An E3 ubiquitin ligase, orthologous to human CUL2 (Kipreos *et al.* 1996; Kipreos 2005). | | | √ | | | √ | | √ | | 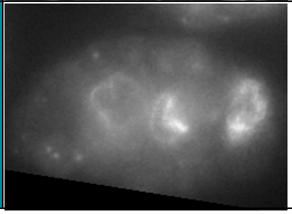 | |

| | | | | | | | | | | | | | | |
|---|---|---|---|---|---|---|---|---|---|---|---|---|---|---|
| | | ZK287.5 | *rbx-1* | Part of the CUL-2 ubiquitin ligase complex. Orthologous to human RBX1 (Sasagawa *et al.* 2003; Kipreos 2005). | | | √ | √ | √ | √ | | | | 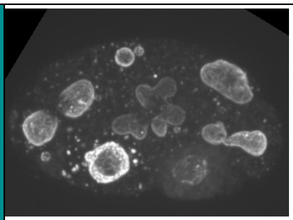 | |
| | | F52C6.12 | | Orthologous to ubiquitin conjugating enzymes (Michelle *et al.* 2009). According to Wormbase it may be a pseudogene. | | | | √ | | √ | | | | 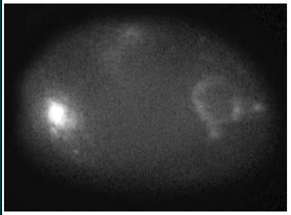 | |
| | COP9 signalosome | Y59A8A.1 | *csn-1* | A COP9 signalosome complex subunit, regulates E3 ubiquitin ligases. Orthologous to human GPS1 (Pintard *et al.* 2003a). | √ | | √ | | | √ | | | | 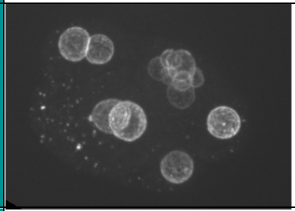 | |
| | | B0547.1 | *csn-5* | A COP9 signalosome complex subunit, which regulates E3 ubiquitin ligases. Orthologous to human COPS5 (Pintard *et al.* 2003a). | | | √ | | | | | | | 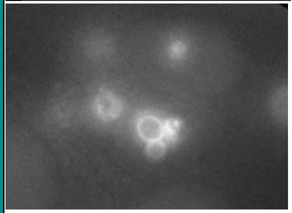 | |
| | **Chromatin and chromosome structure** | | | | | | | | | | | | | | |
| | Condensin complex | M106.1 | *mix-1* | A component of dosage compensation complex, homologous to SMC2. Required for X chromosome dosage compensation and chromosome condensation (Lieb *et al.* 1996; Hagstrom *et al.* 2002). | | | √ | | √ | √ | | | | 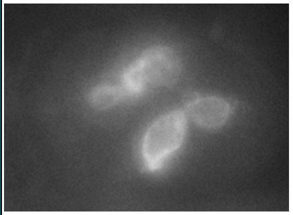 | Anaphase bridges (Csankovszki *et al.* 2009); Pronuclear size defect (Piano *et al.* 2002). |

| | | | | | | | | | | | | | | | |
|---|---|---|---|---|---|---|---|---|---|---|---|---|---|---|---|
| | | Y110A7A.1 | *hcp-6* | A condensin complex subunit, required for chromosome condensation (Stear and Roth 2002). | | | √ | √ | √ | √ | | | | 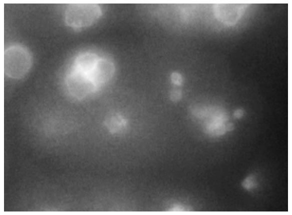 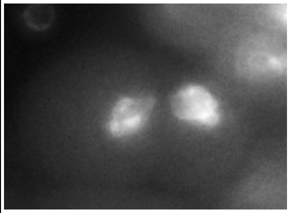 | Anaphase bridges (Csankovszki *et al.* 2009); sister chromatid separation defective causing deformed nuclei (Sonnichsen *et al.* 2005); pronuclear size defect (Piano *et al.* 2002). |
| | | F55C5.4 | *capg-2* | A CAP-G condensin subunit, required for chromosome condensation (Ono *et al.* 2003; Csankovszki *et al.* 2009). | | | √ | | √ | √ | | | | 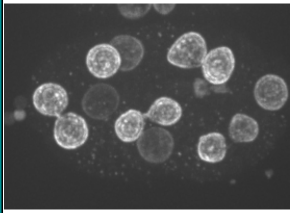 | Anaphase bridges (Csankovszki *et al.* 2009); multi-nucleated cells (Echard 2004). |
| | Cohesion complex | F18E2.3 | *scc-3* | A cohesin complex subunit. Essential for sister chromatid cohesion (Pasierbek *et al.* 2003; Wang *et al.* 2003). | | | | √ | √ | | | | | 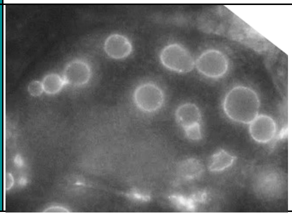 | Sister chromatid separation defective causing deformed nuclei (Sonnichsen *et al.* 2005). |

| | | | | | | | | | | | | | | | |
|---|---|---|---|---|---|---|---|---|---|---|---|---|---|---|---|
| | | Y47D3A.26 | *smc-3* | A cohesin complex subunit. SMC-3 is required for DNA condensation, cohesion and DNA repair (Chan *et al.* 2003; Baudrimont *et al.* 2011). | √ | | | √ | √ | | | | | 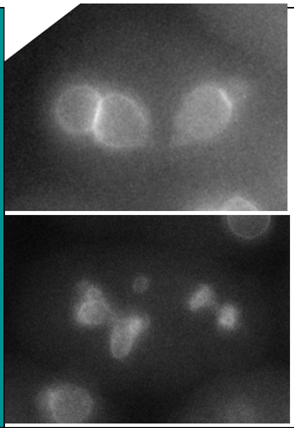 | |
| | Histone genes | ZK131.7 | *his-13* | Histone H3 (Roberts *et al.* 1989). | | | | √ | √ | √ | √ | | | 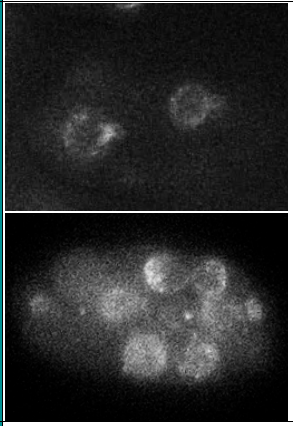 | Anaphase bridges (Kodama *et al.* 2002). |
| | | F45E1.6 | *his-71* | A Histone H3.3 variant (Ooi *et al.* 2006). | | | | | | | √ | | | 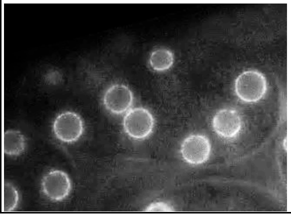 | |

| | | | | | | | | | | | | | | | |
|---|---|---|---|---|---|---|---|---|---|---|---|---|---|---|---|
| | Other | ZK1127.7 | *cin-4* | Orthologous to human topoisomerase II (Stanvitch and Moore 2008). | | | | √ | | √ | | | | 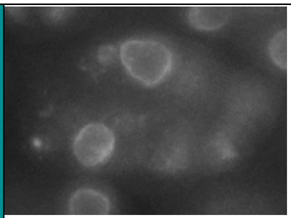 | Multi-nucleated cells, pronuclear size defect (Piano *et al.* 2002); Sister chromatid separation defective causing deformed nuclei (Sonnichsen *et al.* 2005). |
| | | D1081.8 | *phi-7/ cdc-5L* | A putative ortholog of human CDC5L (cell division cycle 5 like). cdc-5L is predicted to have DNA binding activity. | | | | | | | √ | | | 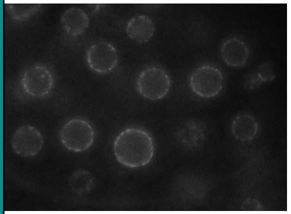 | Multi-nucleated cells in early embryos (Piano *et al.* 2002). |
| | **Nuclear envelope proteins** | | | | | | | | | | | | | | |
| | Nuclear lamina associated proteins | F28B12.3 | *vrk-1* | A serine/threonine vaccinia-related protein kinase. VRK-1 is essential for nuclear envelope reassembly at the end of mitosis (Gorjanacz *et al.* 2007). | | | | | | | | √ | | 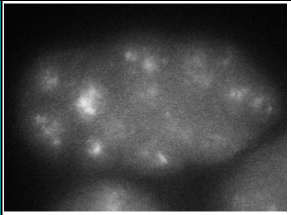 | Nuclear appearance variant (Sonnichsen *et al.* 2005). |

| | | | | | | | | | | | | | | | |
|---|---|---|---|---|---|---|---|---|---|---|---|---|---|---|---|
| | | DY3.2 | *lmn-1* | A nuclear envelope protein required for chromatin organization and structural integrity of the nucleus. It is the only lamin ortholog in *C. elegans* (Riemer *et al.* 1993; Liu *et al.* 2000). | √ | | √ | √ | √ | √ | √ | | | 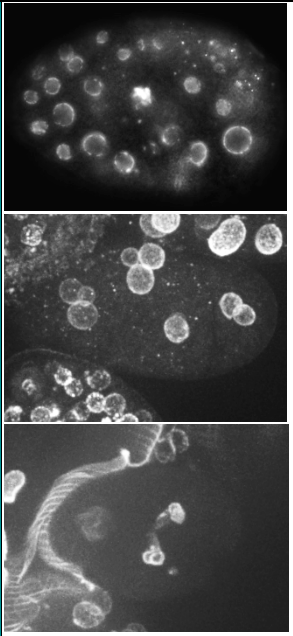 | Multi-nucleated oocytes; small nuclei (Green *et al.* 2011); nuclear appearance variant (Liu *et al.* 2000; Green *et al.* 2011) multi-nucleated cells (Gorjanacz and Mattaj 2009); pronuclear size defect (Fraser *et al.* 2000; Zipperlen *et al.* 2001). |
| | | F57B1.2 | *sun-1* | A SUN-domain containing inner nuclear envelope protein. Associates with ZYG-12 to form the LINC complex. Required for centrosome attachment to the nuclear envelope, nuclear movement and homologous chromosome pairing in meiosis (Malone *et al.* 1999; Malone *et al.* 2003). | | | √ | | | | | | | 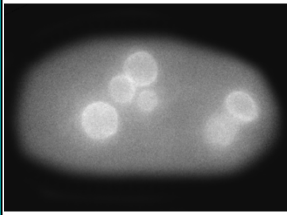 | |

| | | | | | | | | | | | | | | |
|---|---|---|---|---|---|---|---|---|---|---|---|---|---|---|
| | Nuclear import and export | Y48G1A.5 | *xpo-2* | An ortholog of human CSE1L. A putative importin-β, involved in nuclear export. | | | | | √ | √ | | | | 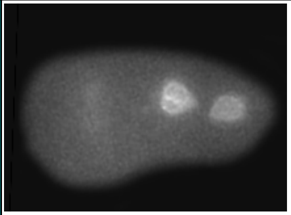 | Pronuclear envelope defect in early embryos (Fraser *et al.* 2000; Zipperlen *et al.* 2001; Sonnichsen *et al.* 2005). |
| | | K01G5.4 | *ran-1* | A Ran GTPase orthologous to human RAN. Mediates nucleocytoplasmic transport (Askjaer *et al.* 2002; Bamba *et al.* 2002). | | | | | | | √ | | | 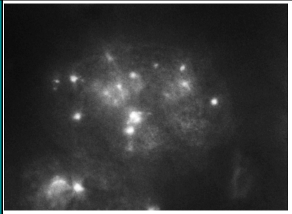 | Multi-nucleated oocytes, enlarged nuclei (Green *et al.* 2011); small nuclei (Askjaer *et al.* 2002; Green *et al.* 2011); pronuclear appearance defect (Sonnichsen *et al.* 2005); pronuclear size defect (Askjaer *et al.* 2002). |
| | | C26D10.1 | *ran-3* | A guanine exchange factor (GEF) of RAN-1, orthologous to human RCC1, involved in nucleocytoplasmic transport (Askjaer *et al.* 2002; Bamba *et al.* 2002). | | | √ | | √ | | √ | √ | | 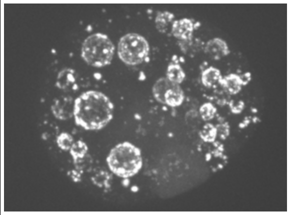 | Nuclear appearance variant, small nuclei (Green *et al.* 2011); multi-nucleated cells (Piano *et al.* 2002); pronuclear appearance defect (Sonnichsen *et al.* 2005); pronuclear size defect (Askjaer *et al.* 2002). |
| | Nuclear pore complex subunits | T01G9.4 | *npp-2* | A subunit for the nuclear pore complex that mediates nucleocytoplasmic transport. *npp-2* is orthologous to human NUP85 (Galy *et al.* 2003). | | | | | | | √ | | | 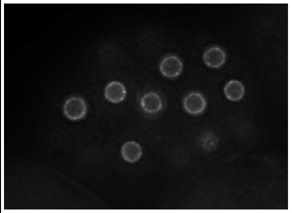 | Pronuclear appearance defect (Sonnichsen *et al.* 2005); nuclear morphology variation in early embryos (Fraser *et al.* 2000; Zipperlen *et al.* 2001; Galy *et al.* 2003). |
| | | F59A2.1 | *npp-9* | A subunit for the nuclear pore complex that mediates nucleocytoplasmic transport. npp-9 is orthologous to Nup358/RANBP2 (RAN binding protein 2) (Galy *et al.* 2003). | | | | | | √ | | √ | | 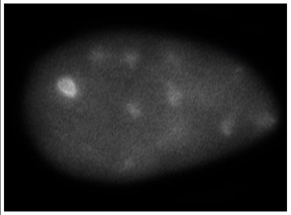 | Pronuclear appearance defect (Sonnichsen *et al.* 2005); nuclear morphology variation in early embryo (Galy *et al.* 2003). |

| | | | | | | | | | | | | | | | |
|---|---|---|---|---|---|---|---|---|---|---|---|---|---|---|---|
| | | Y77E11A.13 | *npp-20* | A subunit for the nuclear pore complex that mediates nucleocytoplasmic transport. NPP-20 is orthologous to SEC13 (Casadio *et al.* 2015). | | | √ | | | √ | √ | | | 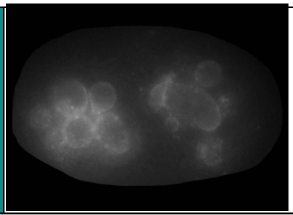 | Nuclear morphology variation in early embryo (Galy *et al.* 2003). |
| | **Vesicle trafficking** | | | | | | | | | | | | | | |
| | | F53G12.1 | *rab-11.1* | A Rab GTPase that functions in endocytosis and regulates endocytic protein sorting (Grant and Hirsh 1999). | | | √ | | | √ | | | | 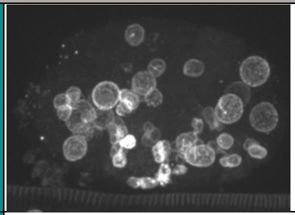 | Multi-nucleated cells (Poteryaev *et al.* 2007); nuclear appearance variant (Green *et al.* 2011). |
| | | K02D10.5 | *snap-29* | A syntaxin-type SNARE, part of t-SNAREs. Involved in vesicle trafficking (Sato *et al.* 2011). | | √ | | | | | | | | 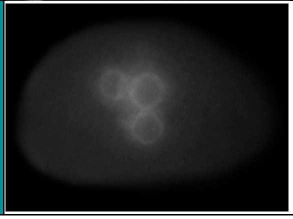 | Multi-nucleated oocytes, small nuclei (Green *et al.* 2011). |

| | | | | | | | | | | | | | | | |
|---|---|---|---|---|---|---|---|---|---|---|---|---|---|---|---|
| | | T01B11.3 | *syx-4* | A syntaxin-type SNARE, part of t-SNAREs. Involved in vesicle trafficking (Jantsch-Plunger and Glotzer 1999). | √ | | √ | √ | √ | √ | | | | 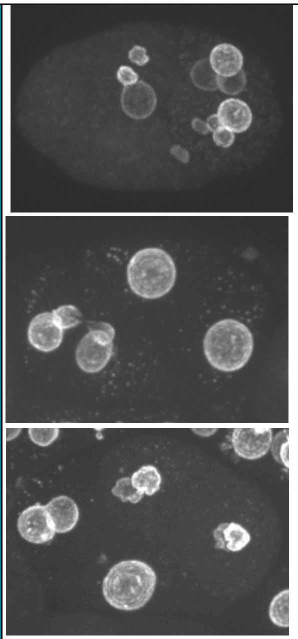 | Multi-nucleated cells (Jantsch-Plunger and Glotzer 1999); enlarged nuclei (Green *et al.* 2011). |
| | | F55A11.2 | *syx-5* | A syntaxin-type SNARE, part of t-SNARE. Involved in vesicle trafficking (Hyenne *et al.* 2015). | √ | | √ | | | √ | | | | 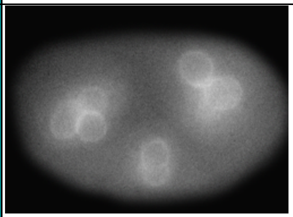 | Multi-nucleated cells (Echard *et al.* 2004). |
| | | M110.5 | *dab-1* | Orthologous to DISABLED, a clathrin binding protein. DAB-1 is required for receptor-mediated endocytosis (Kamikura and Cooper 2003). | | | | | | | √ | | | 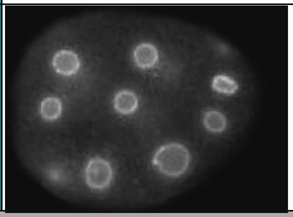 | |
| | **Mitochondrial functions** | | | | | | | | | | | | | | |

| | | | | | | | | | | | | | | |
|---|---|---|---|---|---|---|---|---|---|---|---|---|---|---|
| Mitochondrial fission | C02C6.1 | *dyn-1* | A dynamin ortholog, required for various membrane fusion events, including endocytosis, apoptosis and mitochondrial fission (Clark *et al.* 1997). | √ | | √ | | | | | | | 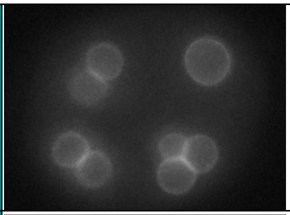 | Multi-nucleated oocytes (Green *et al.* 2011). |
| | T12E12.4 | *drp-1* | A dynamin-related protein required for mitochondrial fission (Labrousse *et al.* 1999). | √ | | √ | | | √ | | | | 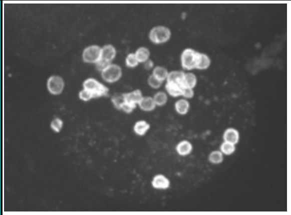 | Multi-nucleated cells (Echard *et al.* 2004). |
| Electron transport chain | H28O16.1 | *atp-1/phi-37* | A putative alpha subunit of mitochondrial ATP synthase, orthologous to human ATP5F1A. | | | | | | √ | √ | | | 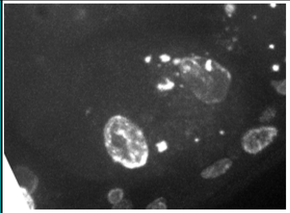 | *(Rahman et al. 2014)* |
| | C34E10.6 | *atp-2* | A putative beta subunit of mitochondrial ATP synthase, orthologous to human ATP5F1B. | | √ | | √ | | √ | | | | 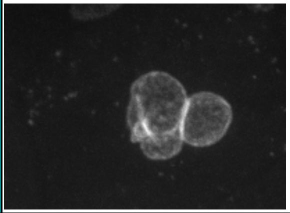 | *(Rahman et al. 2014)* |
| | F27C1.7 | *atp-3* | A putative OSCP subunit of mitochondrial ATP synthase, orthologous to human ATP5O. | | | | | | √ | | √ | | 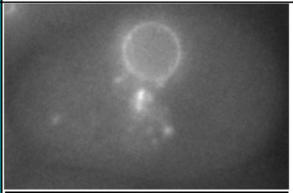 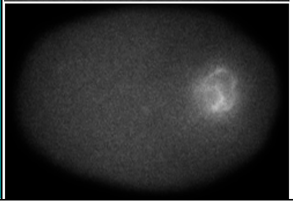 | *(Rahman et al. 2014)* |

| | | | | | | | | | | | | | | | |
|---|---|---|---|---|---|---|---|---|---|---|---|---|---|---|---|
| | | T05H4.12 | *atp-4* | A putative $F_6$ subunit for mitochondrial ATP synthase, orthologous to human ATP5J. | | | | | | √ | | √ | | 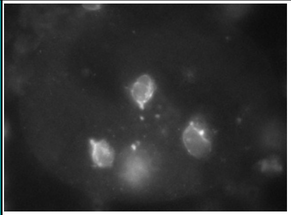 | *(Rahman et al. 2014)* |
| | | C06H2.1 | *atp-5* | A putative D subunit for mitochondrial ATP synthase, orthologous to human ATP5D. | | √ | | √ | | | | | | 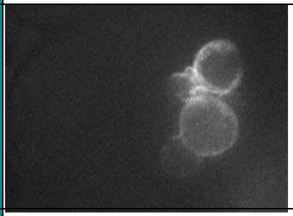 | (Rahman *et al.* 2014) |
| | | C54G4.8 | *cyc-1* | Orthologous to human CYC1 (cytochrome c1). | | | √ | | | | √ | | | 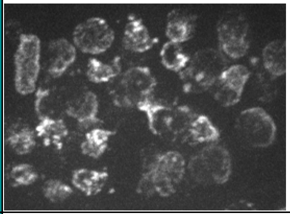 | *(Rahman et al. 2014)* |
| | | F56D2.1 | *ucr-1* | Orthologous to human ubiquinol-cytochrome c reductase core protein. | √ | | | | | | | | | 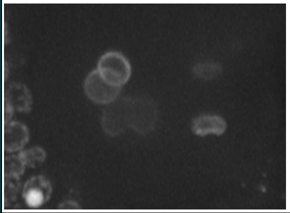 | *(Rahman et al. 2014)* |
| | | F26E4.9 | *cco-1/cox-5b* | Orthologous to human COX5B (cytochrome c oxidase subunit 5B). | √ | | √ | | | | | | | 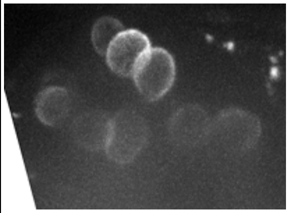 | *(Rahman et al. 2014)* |

| | | | | | | | | | | | | | | | |
|---|---|---|---|---|---|---|---|---|---|---|---|---|---|---|---|
| | | Y37D8A.14 | *cco-2* | Orthologous to human COX5A (cytochrome c oxidase subunit 5A). | | | √ | | | | | | | 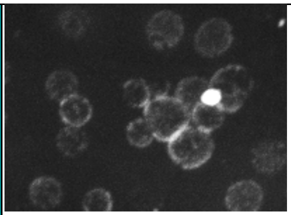 | *(Rahman et al. 2014)* |
| | | F33A8.5 | *sdhd-1* | Orthologous to human succinate dehydrogenase complex subunit D. | √ | | | | | | | | | 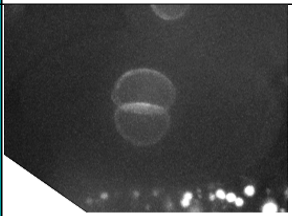 | *(Rahman et al. 2014)* |
| | | T07C4.7 | *mev-1/sdhc-1* | Orthologous to human succinate dehydrogenase complex subunit C. | √ | | | | | | | | | 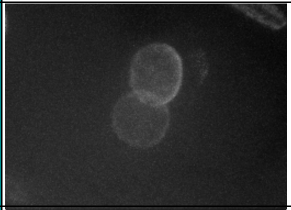 | *(Rahman et al. 2014)* |
| | | K04G7.4 | *nuo-4* | Orthologous to human NADH:ubiquinone oxidoreductase subunit A10. | √ | | | | | | | | | 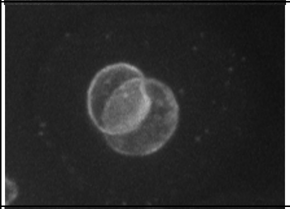 | *(Rahman et al. 2014)* |
| | | *T20H4.5* | | Orthologous to human NADH:ubiquinone oxidoreductase core subunit S8. | √ | √ | | | | | | | | 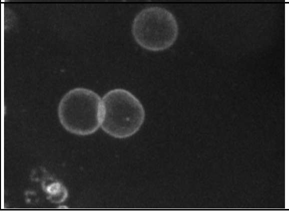 | *(Rahman et al. 2014)* |

| | | | | | | | | | | | | | | | |
|---|---|---|---|---|---|---|---|---|---|---|---|---|---|---|---|
| | Mitochondrial protein Degradation | Y47G6A.10 | *spg-7* | A putative mitochondrial-AAA metalloprotease that regulates mitochondrial protein degradation (Pena *et al.* 2016). | | | √ | √ | | √ | | | | 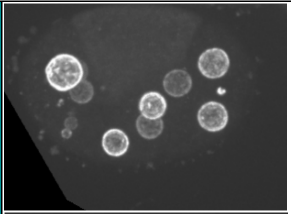 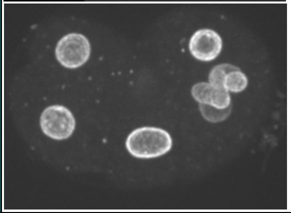 | |
| | **Protein folding** | | | | | | | | | | | | | | |
| | Chaperonin | T05C12.7 | *cct-1* | A subunit of T complex chaperonin. CCT-1 functions as an ATP-dependent molecular chaperone (Leroux and Candido 1995). | | | √ | √ | | √ | | | | 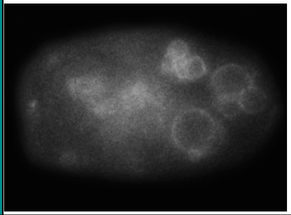 | Nuclear appearance variant, small nuclei (Green *et al.* 2011). |
| | | T21B10.7 | *cct-2* | A subunit of T complex chaperonin. CCT-2 functions as an ATP-dependent molecular chaperone (Leroux and Candido 1995). | | | √ | | | √ | | | | 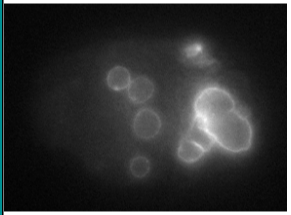 | |
| | | C07G2.3 | *cct-5* | A subunit of T complex chaperonin. CCT-5 functions as an ATP-dependent molecular chaperone (Leroux and Candido 1995). | | | √ | | | √ | | | | 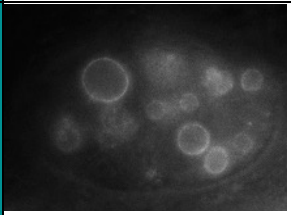 | Multi-nucleated oocytes, nuclear appearance variant (Green *et al.* 2011). |

| | | | | | | | | | | | | | | | |
|---|---|---|---|---|---|---|---|---|---|---|---|---|---|---|---|
| | | F01F1.8 | *cct-6* | A subunit of T complex chaperonin. CCT-6 functions as an ATP-dependent molecular chaperone (Leroux and Candido 1995). | | √ | | | | √ | | | | 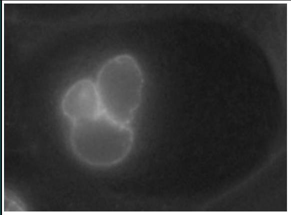 | Multi-nucleated oocytes, nuclear appearance variant (Green *et al.* 2011). |
| | | T10B5.5 | *cct-7* | A putative subunit of T complex chaperonin. CCT-7 functions as an ATP-dependent molecular chaperone (Leroux and Candido 1995). | | | √ | √ | | √ | | | | 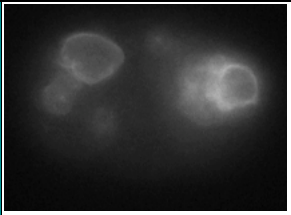 | Severe pleiotropic defects including multiple pronuclei in early embryos (Sonnichsen *et al.* 2005). |
| | | F39B2.10 | *dnj-12* | An Hsp40-like J-domain. DNJ-12 has a role in protein folding along with Hsp70 (Nillegoda *et al.* 2015). | | | √ | √ | | √ | | | | 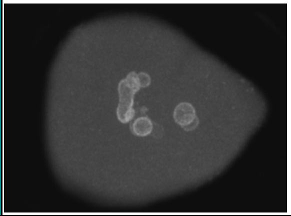 | |
| | | C30C11.4 | *hsp-110* | A member of Hsp70 family of heat shock proteins (Nillegoda *et al.* 2015). | | | √ | √ | | √ | | | | 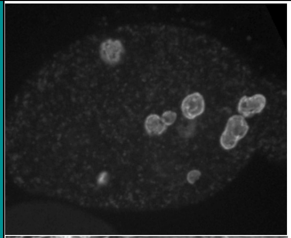 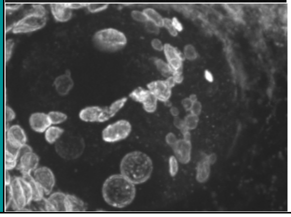 | |

| | | | | | | | | | | | | | | | |
|---|---|---|---|---|---|---|---|---|---|---|---|---|---|---|---|
| | | Y65B4BR.5 | *lcd-2* | An ortholog of human NACA (nascent polypeptide-associated complex subunit alpha) (Arsenovic *et al.* 2012). | | | √ | | | √ | | | √ | 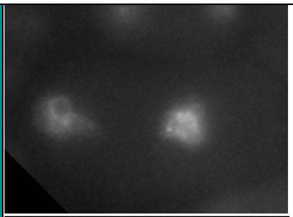 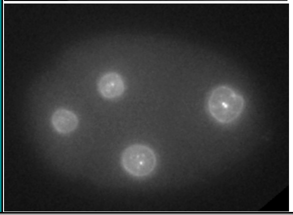 | |
| | **Vacuolar proteins** | | | | | | | | | | | | | | |
| | V-ATPase | R10E11.8 | *vha-1* | A vacuolar V-ATPase c subunit, an ortholog of human ATP6V0C (Oka *et al.* 1997; Lee *et al.* 2010). | | | √ | √ | | √ | | | | 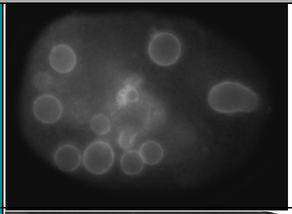 | Multi-nucleated in oocytes, nuclear appearance variant, enlarged nuclei (Green *et al.* 2011). |
| | | R10E11.2 | *vha-2* | A vacuolar V-ATPase c subunit, an ortholog of human ATP6V0C (Oka *et al.* 1997; Lee *et al.* 2010). | | √ | | | | √ | | | | 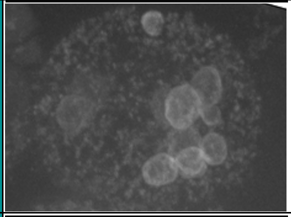 | Nuclear appearance variant, enlarged nuclei (Green *et al.* 2011). |
| | | T01H3.1 | *vha-4* | A vacuolar V-ATPase c'' subunit, an ortholog of human ATP6V0B (Oka *et al.* 1997; Lee *et al.* 2010). | √ | | √ | | | | | | | 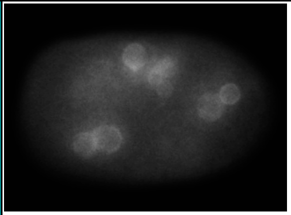 | Multi-nucleated oocytes, nuclear appearance variant (Green *et al.* 2011). |

| | | | | | | | | | | | | | | | |
|---|---|---|---|---|---|---|---|---|---|---|---|---|---|---|---|
| | | C17H12.14 | *vha-8* | A vacuolar V-ATPase E subunit (Oka *et al.* 1997; Choi *et al.* 2003; Ji *et al.* 2006; Lee *et al.* 2010). | √ | | √ | √ | | √ | | | | 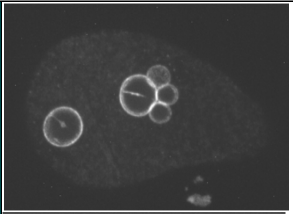 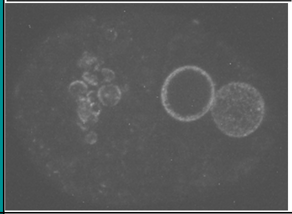 | Multi-nucleated early embryos (Choi *et al.* 2003); nuclear appearance variant (Green *et al.* 2011). |
| | | ZK970.4 | *vha-9* | A vacuolar V-ATPase F subunit, an ortholog of human ATP6V1F (Lee et al. 2010). | | | √ | | | √ | | | | 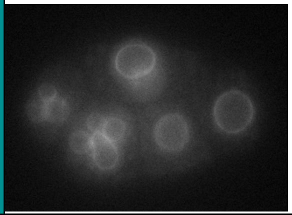 | Nuclear appearance variant, enlarged nuclei (Green *et al.* 2011). |
| | | Y38F2AL.3 | *vha-11* | A vacuolar V-ATPase C subunit, an ortholog of human ATP6V1C1 (Oka and Futai 2000; Lee *et al.* 2010). | | | √ | | | √ | | | | 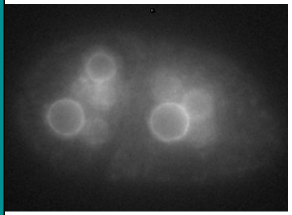 | Small nuclei (Green *et al.* 2011). |
| | | F55H2.2 | *vha-14* | A vacuolar V-ATPase D subunit, an ortholog of human ATP6V1D (Lee *et al.* 2010). | | | √ | | | | | | | 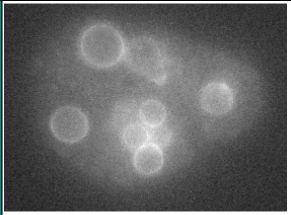 | Nuclear appearance variant (Green *et al.* 2011). |

| | | | | | | | | | | | | | | |
|---|---|---|---|---|---|---|---|---|---|---|---|---|---|---|
| | | ZK637.8 | *unc-32* | A vacuolar V-ATPase a subunit, an ortholog of human ATP6V0A1 (Pujol *et al.* 2001; Lee *et al.* 2010). | | | | | | √ | √ | | | 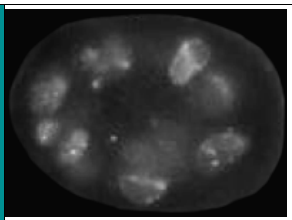 | |
| | **Intermediary metabolism** | | | | | | | | | | | | | | |
| | Glycosylation-related processes | T23G11.2 | *gna-2* | A Glucosamine phosphate N-acetyl transferase. GNA-2 is required for alpha-GalNAc-modifications of mucin-like proteins, and chitin (Johnston *et al.* 2006). | √ | | √ | | | √ | | | | 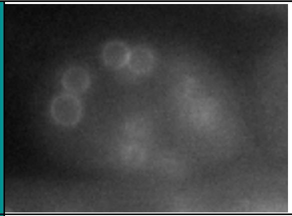 | Multi-nucleated cells (Lee and Schedl 2001). |
| | | F29F11.1 | *sqv-4* | A UDP-glucose 6-dehydrogenase. SQV-4 is essential for the biosynthesis of chondroitin and heparan sulfate proteoglycans (Hwang and Horvitz 2002). | | | √ | | | | | | | 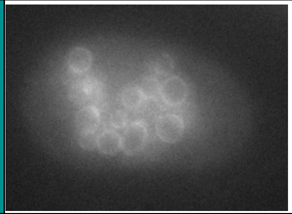 | |
| | | T24D1.1 | *sqv-5* | Chondroitin sulphate synthase, which is similar to human chitin synthase. SQV-5 is required for the biosynthesis of chondroitin (Hwang *et al.* 2003). | √ | | √ | | | | | | | 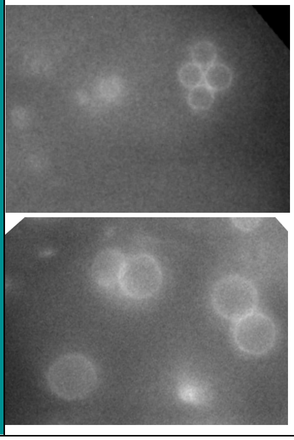 | Multi-nucleated cells (Mizuguchi *et al.* 2003). |

| | | | | | | | | | | | | | | |
|---|---|---|---|---|---|---|---|---|---|---|---|---|---|---|
| | | T01H3.4 | *perm-1* | A sugar modifying enzyme, possibly required for the formation of cytidine diphosphate (CDP)-ascarylose in the eggshell (Olson *et al.* 2012). | | | √ | √ | | √ | | | | 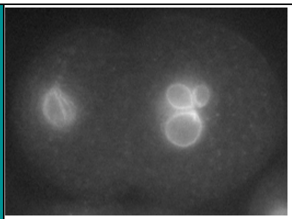 |
| | | ZK686.3 | | An ortholog of human MAGT1 and TUSC3. Predicted to be a protein glycosyltransferase subunit. | √ | | √ | | | √ | | | | 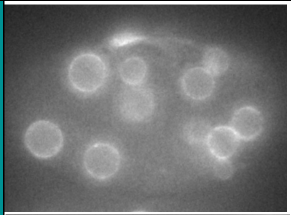 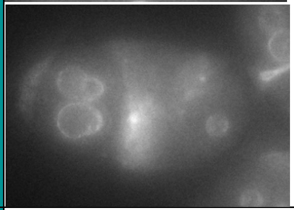 |
| | Cytochrome p450 | Y17G9B.3 | *cyp-31A3* | Homologous to human CYP7B1, a member of Cytochrome P450 family protein (Benenati et al. 2009). | √ | | | | | √ | | | | 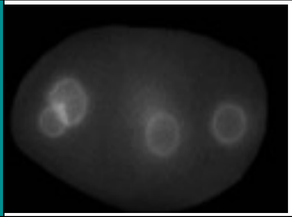 |
| | Phosphate metabolism | C47E12.4 | *pyp-1* | An inorganic pyrophosphatase (Ko et al. 2007). | | | √ | | | √ | | | | 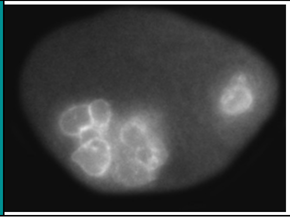 |

| | | | | | | | | | | | | | | | |
|---|---|---|---|---|---|---|---|---|---|---|---|---|---|---|---|
| | Nucleoside and nucleotide biosynthesis | R06C7.5 | *adsl-1* | An ortholog of human ADSL, an adenylosuccinate lyase. ADSL-1 functions de novo purine biosynthesis (Marsac et al. 2019). | | | √ | | | √ | | | | 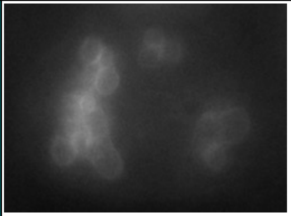 | Nuclear morphology variation (Fraser *et al.* 2000). |
| | | C29E4.8 | *let-754* | An ortholog of human AK2, an adenylate kinase. | | | √ | | | √ | | | | 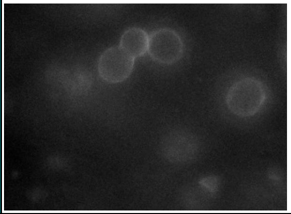 | Nuclear appearance variant (Green *et al.* 2011). |
| | Gluconeogenesis and glycolysis | F01F1.12 | *aldo-2* | An ortholog of human ALDOA and ALDOC, which have Fructose-bisphosphate aldolase activity. ALDO-2 is involved in glucogenesis and glycolysis. | | | √ | | | √ | | √ | | 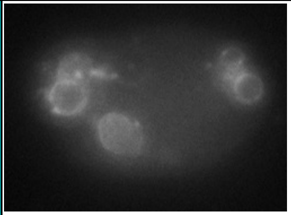 | |
| | | Y46G5A.31 | *gsy-1* | A glycogen synthase, orthologous to human GSY1 and GSY2 (Seo et al. 2018). | | | | | | | √ | | | 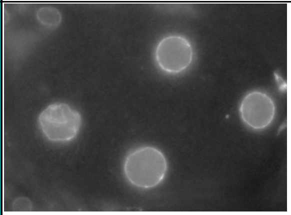 | |
| | | T20G5.2 | *cts-1* | A citrate synthase ortholog (Hada et al. 2019). | √ | | | | | | | | | 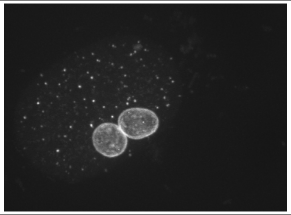 | (*Rahman et al. 2014*) |

| | | | | | | | | | | | | | | | |
|---|---|---|---|---|---|---|---|---|---|---|---|---|---|---|---|
| | | F23B12.5 | *dlat-1* | Orthologous to human dihydrolipoyllysine acetyltransferase, a component of the pyruvate dehydrogenase complex. | √ | | | | | | | | | 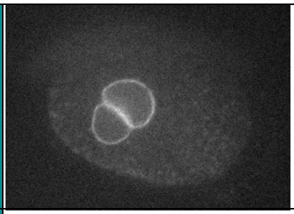 | (*Rahman et al. 2014*) |
| | | F42A8.2 | *sdhb-1* | Orthologous to human succinate dehydrogenase complex iron sulfur subunit B (Huang and Lemire 2009). | √ | | | | | | | | | 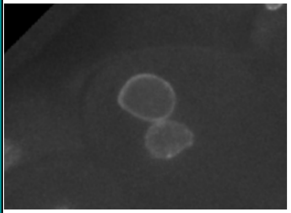 | (*Rahman et al. 2014*) |
| | | F54H12.1 | *aco-2* | Orthologous to human aconitase 2. | √ | | | | | | | | | 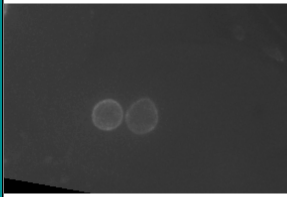 | (*Rahman et al. 2014*) |
| | | F46E10.10 | *mdh-2* | Orthologous to human malate dehydrogenase. | √ | | | | | | | | | 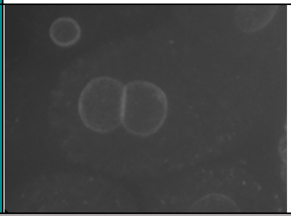 | (*Rahman et al. 2014*) |
| | **Other** | | | | | | | | | | | | | | |

| | | | | | | | | | | | | | | | |
|---|---|---|---|---|---|---|---|---|---|---|---|---|---|---|---|
| | | F22D6.6 | *ekl-1* | A putative small RNA regulator. EKL-1 is a tudor domain containing protein (Rocheleau et al. 2008). | √ | | √ | | | √ | | | √ | 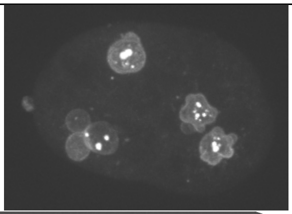 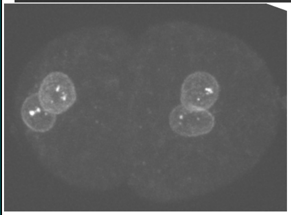 | Anaphase bridges, nuclear appearance variant (Claycomb *et al.* 2009). |
| | | F20D12.1 | *csr-1* | An Argonaute protein. CSR-1 functions by cleaving siRNA-paired mRNA (Yigit et al. 2006; Wedeles et al. 2013). | | | √ | √ | | √ | | | | 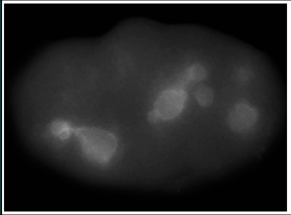 | Anaphase bridges, nuclear appearance variant (Claycomb *et al.* 2009). |
| | | F41E6.4 | *smk-1* | An ortholog of human subunits of the PP4 protein phosphatase (Wolff et al. 2006). | | | | √ | | | | | | 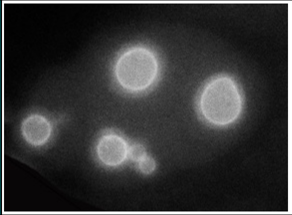 | |
| | | R119.4 | *pqn-59* | Orthologous to human ubiquitin associated proteins UBAP and UBAPL. | | √ | | | | √ | | | | 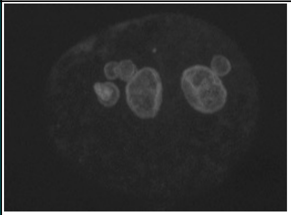 | |

| | | | | | | | | | | | | | | | |
|---|---|---|---|---|---|---|---|---|---|---|---|---|---|---|---|
| | | B0365.3 | *eat-6* | A sodium/potassium ATPase, orthologous to human ATP1A1 and ATP1A3 (Davis et al. 1995). | | | √ | | | | | | | 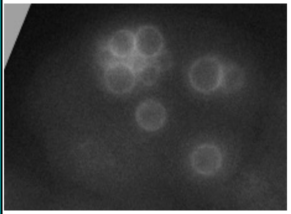 | |
| | | K04F10.4 | *bli-4* | An endoprotease, belongs to the kex2/subtilisin-like proprotein convertase family that processes proproteins for secretion (Thacker et al. 1995). | | | | | | √ | | | | 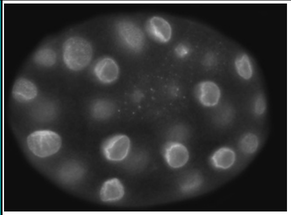 | |
| | | Y37D8A.10 | *spcs-2* | An ortholog of human SPCS2, a signal peptidase complex subunit. | | | √ | √ | | | | | | 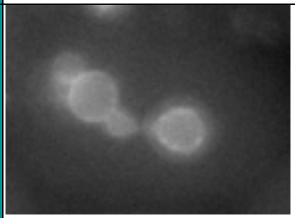 | Nuclear appearance variant (Green *et al.* 2011). |
| | | ZC101.2 | *unc-52* | Orthologous to the human HSPG2 perlecan, a basement membrane heparan sulfate proteoglycan. Part of the worm extracellular matrix (Rogalski et al. 1995). | | √ | | | | √ | | | | 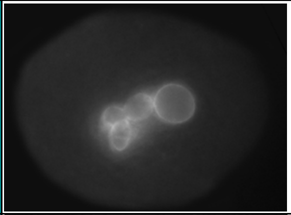 | |
| | | C17G10.2 | *ttc-4* | An ortholog of human TTC4, a tetratricopeptide repeat domain containing protein of unknown function. | | | √ | | | | | | | 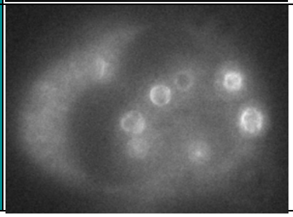 | |

| | | | | | | | | | | | | | | |
|---|---|---|---|---|---|---|---|---|---|---|---|---|---|---|
| | | F37C12.14 | | Unknown function. | √ | | √ | | | | | | | 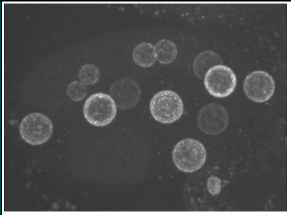 |
| | | T28D6.6 | | Orthologous to human DRG1 (developmentally regulated GTP binding protein 1). | | | √ | √ | | √ | | | | 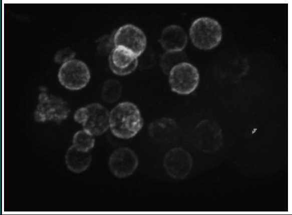 |
| | | Y18D10A.20 | *pfn-1* | A profilin, an actin associated protein (Severson *et al.* 2002; Velarde *et al.* 2007). | | | √ | | | | | | | 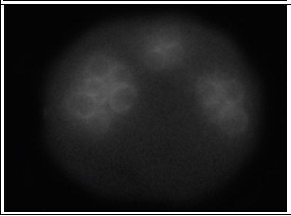 |

[1] Protein function is based on WormBase and the indicated references (attached below), if available.
[2] Ortholog designations are based on (Shaye and Greenwald 2011; Kim *et al.* 2018).
[3] References originating from this screen are italicized.

**Supplemental Table S2**: Connectivity of phenotypic categories.

| Phenotypes | Phenotypes | | | | | | | | |
|---|---|---|---|---|---|---|---|---|---|
| | Paired Nuclei | 1-cell multi-nuc. | ≥ 2 cell Multi nuc. | Micronuclei | Anaphase bridges | Deformed | Abnormal NE NPC distribution | Cytoplasmic NPCs | Intra nuclear NPCs |
| **Paired nuclei** | (50)[a] | | | | | | | | |
| **1-cell Multi-nucleated** | 1[b] | (13) | | | | | | | |
| **≥ 2 cell Multi nucleated** | 32 | 0 | (106) | | | | | | |
| **Micronuclei** | 12 | 4 | 36 | (54) | | | | | |
| **Anaphase bridges** | 11 | 0 | 20 | 18 | (33) | | | | |
| **Deformed** | 26 | 9 | 71 | 40 | 18 | (104) | | | |
| **Abnormal NE NPC distribution** | 3 | 0 | 7 | 4 | 4 | 9 | (27) | | |
| **Cytoplasmic NPCs** | 3 | 0 | 7 | 3 | 3 | 10 | 3 | (13) | |
| **Intra-nuclear NPCs** | 1 | 0 | 2 | 0 | 0 | 2 | 0 | 0 | (2) |

[a] Numbers in parentheses in grey boxes indicate the total number of RNAi that resulted in the indicated phenotype.
[b] Numbers indicate the number of RNAi that resulted in the indicated two phenotypes (note that additional phenotypes are also possible).

Antoshechkin, I., and M. Han, 2002 The C. elegans evl-20 gene is a homolog of the small GTPase ARL2 and regulates cytoskeleton dynamics during cytokinesis and morphogenesis. Dev Cell 2**:** 579-591.

Arsenovic, P. T., A. T. Maldonado, V. D. Colleluori and T. A. Bloss, 2012 Depletion of the C. elegans NAC engages the unfolded protein response, resulting in increased chaperone expression and apoptosis. PLoS One 7**:** e44038.

Askjaer, P., V. Galy, E. Hannak and I. W. Mattaj, 2002 Ran GTPase cycle and importins alpha and beta are essential for spindle formation and nuclear envelope assembly in living Caenorhabditis elegans embryos. Mol Biol Cell 13**:** 4355-4370.

Audhya, A., F. Hyndman, I. X. McLeod, A. S. Maddox, J. R. Yates, 3rd *et al.*, 2005 A complex containing the Sm protein CAR-1 and the RNA helicase CGH-1 is required for embryonic cytokinesis in Caenorhabditis elegans. J Cell Biol 171**:** 267-279.

Aumais, J. P., S. N. Williams, W. Luo, M. Nishino, K. A. Caldwell *et al.*, 2003 Role for NudC, a dynein-associated nuclear movement protein, in mitosis and cytokinesis. J Cell Sci 116**:** 1991-2003.

Bamba, C., Y. Bobinnec, M. Fukuda and E. Nishida, 2002 The GTPase Ran regulates chromosome positioning and nuclear envelope assembly in vivo. Curr Biol 12**:** 503-507.

Banerjee, D., X. Chen, S. Y. Lin and F. J. Slack, 2010 kin-19/casein kinase Ialpha has dual functions in regulating asymmetric division and terminal differentiation in C. elegans epidermal stem cells. Cell Cycle 9**:** 4748-4765.

Barbee, S. A., A. L. Lublin and T. C. Evans, 2002 A novel function for the Sm proteins in germ granule localization during C. elegans embryogenesis. Curr Biol 12**:** 1502-1506.

Barberan-Soler, S., and A. M. Zahler, 2008 Alternative splicing regulation during C. elegans development: splicing factors as regulated targets. PLoS Genet 4**:** e1000001.

Baudrimont, A., A. Penkner, A. Woglar, Y. M. Mamnun, M. Hulek *et al.*, 2011 A new thermosensitive smc-3 allele reveals involvement of cohesin in homologous recombination in C. elegans. PLoS One 6**:** e24799.

Bobinnec, Y., M. Fukuda and E. Nishida, 2000 Identification and characterization of Caenorhabditis elegans gamma-tubulin in dividing cells and differentiated tissues. J Cell Sci 113 Pt 21**:** 3747-3759.

Buchwitz, B. J., K. Ahmad, L. L. Moore, M. B. Roth and S. Henikoff, 1999 A histone-H3-like protein in C. elegans. Nature 401**:** 547-548.

Burrows, A. E., B. K. Sceurman, M. E. Kosinski, C. T. Richie, P. L. Sadler *et al.*, 2006 The C. elegans Myt1 ortholog is required for the proper timing of oocyte maturation. Development 133**:** 697-709.

Casadio, A., D. Longman, N. Hug, L. Delavaine, R. Vallejos Baier *et al.*, 2015 Identification and characterization of novel factors that act in the nonsense-mediated mRNA decay pathway in nematodes, flies and mammals. EMBO Rep 16**:** 71-78.

Chan, R. C., A. Chan, M. Jeon, T. F. Wu, D. Pasqualone *et al.*, 2003 Chromosome cohesion is regulated by a clock gene paralogue TIM-1. Nature 423**:** 1002-1009.

Chen, X., M. D. Cuadros and M. Chalfie, 2015 Identification of nonviable genes affecting touch sensitivity in Caenorhabditis elegans using neuronally enhanced feeding RNA interference. G3 (Bethesda) 5**:** 467-475.

Choi, K. Y., Y. J. Ji, B. K. Dhakal, J. R. Yu, C. Cho *et al.*, 2003 Vacuolar-type H+-ATPase E subunit is required for embryogenesis and yolk transfer in Caenorhabditis elegans. Gene 311**:** 13-23.

Clark, S. G., D. L. Shurland, E. M. Meyerowitz, C. I. Bargmann and A. M. van der Bliek, 1997 A dynamin GTPase mutation causes a rapid and reversible temperature-inducible locomotion defect in C. elegans. Proc Natl Acad Sci U S A 94**:** 10438-10443.

Claycomb, J. M., P. J. Batista, K. M. Pang, W. Gu, J. J. Vasale *et al.*, 2009 The Argonaute CSR-1 and its 22G-RNA cofactors are required for holocentric chromosome segregation. Cell 139**:** 123-134.

Colombo, K., S. W. Grill, R. J. Kimple, F. S. Willard, D. P. Siderovski *et al.*, 2003 Translation of polarity cues into asymmetric spindle positioning in Caenorhabditis elegans embryos. Science 300**:** 1957-1961.

Csankovszki, G., K. Collette, K. Spahl, J. Carey, M. Snyder *et al.*, 2009 Three distinct condensin complexes control C. elegans chromosome dynamics. Curr Biol 19**:** 9-19.

Dammermann, A., T. Muller-Reichert, L. Pelletier, B. Habermann, A. Desai *et al.*, 2004 Centriole assembly requires both centriolar and pericentriolar material proteins. Dev Cell 7**:** 815-829.

Davy, A., P. Bello, N. Thierry-Mieg, P. Vaglio, J. Hitti *et al.*, 2001 A protein-protein interaction map of the Caenorhabditis elegans 26S proteasome. EMBO Rep 2**:** 821-828.

Dawe, A. L., K. A. Caldwell, P. M. Harris, N. R. Morris and G. A. Caldwell, 2001 Evolutionarily conserved nuclear migration genes required for early embryonic development in Caenorhabditis elegans. Dev Genes Evol 211**:** 434-441.

Delattre, M., S. Leidel, K. Wani, K. Baumer, J. Bamat *et al.*, 2004 Centriolar SAS-5 is required for centrosome duplication in C. elegans. Nat Cell Biol 6**:** 656-664.

Dorsett, M., and T. Schedl, 2009 A role for dynein in the inhibition of germ cell proliferative fate. Mol Cell Biol 29**:** 6128-6139.

Dow, M. R., and P. E. Mains, 1998 Genetic and molecular characterization of the caenorhabditis elegans gene, mel-26, a postmeiotic negative regulator of mei-1, a meiotic-specific spindle component. Genetics 150**:** 119-128.

Echard, A., 2004 [Dissecting cytokinesis: proteomic and genomic approaches]. Med Sci (Paris) 20**:** 845-846.

Echard, A., G. R. Hickson, E. Foley and P. H. O'Farrell, 2004 Terminal cytokinesis events uncovered after an RNAi screen. Curr Biol 14**:** 1685-1693.

Ellis, G. C., J. B. Phillips, S. O'Rourke, R. Lyczak and B. Bowerman, 2004 Maternally expressed and partially redundant beta-tubulins in Caenorhabditis elegans are autoregulated. J Cell Sci 117**:** 457-464.

Encalada, S. E., P. R. Martin, J. B. Phillips, R. Lyczak, D. R. Hamill *et al.*, 2000 DNA replication defects delay cell division and disrupt cell polarity in early Caenorhabditis elegans embryos. Dev Biol 228**:** 225-238.

Fraser, A. G., R. S. Kamath, P. Zipperlen, M. Martinez-Campos, M. Sohrmann *et al.*, 2000 Functional genomic analysis of C. elegans chromosome I by systematic RNA interference. Nature 408**:** 325-330.

Frazier, T., D. Shakes, U. Hota and L. Boyd, 2004 Caenorhabditis elegans UBC-2 functions with the anaphase-promoting complex but also has other activities. J Cell Sci 117**:** 5427-5435.

Galy, V., I. W. Mattaj and P. Askjaer, 2003 Caenorhabditis elegans nucleoporins Nup93 and Nup205 determine the limit of nuclear pore complex size exclusion in vivo. Mol Biol Cell 14**:** 5104-5115.

Gao, X., Y. Teng, J. Luo, L. Huang, M. Li *et al.*, 2014 The survival motor neuron gene smn-1 interacts with the U2AF large subunit gene uaf-1 to regulate Caenorhabditis elegans lifespan and motor functions. RNA Biol 11**:** 1148-1160.

Gassmann, R., A. Essex, J. S. Hu, P. S. Maddox, F. Motegi *et al.*, 2008 A new mechanism controlling kinetochore-microtubule interactions revealed by comparison of two dynein-targeting components: SPDL-1 and the Rod/Zwilch/Zw10 complex. Genes Dev 22**:** 2385-2399.

Gonczy, P., C. Echeverri, K. Oegema, A. Coulson, S. J. Jones *et al.*, 2000 Functional genomic analysis of cell division in C. elegans using RNAi of genes on chromosome III. Nature 408**:** 331-336.

Gorjanacz, M., E. P. Klerkx, V. Galy, R. Santarella, C. Lopez-Iglesias *et al.*, 2007 Caenorhabditis elegans BAF-1 and its kinase VRK-1 participate directly in post-mitotic nuclear envelope assembly. EMBO J 26**:** 132-143.

Gorjanacz, M., and I. W. Mattaj, 2009 Lipin is required for efficient breakdown of the nuclear envelope in Caenorhabditis elegans. J Cell Sci 122**:** 1963-1969.

Grant, B., and D. Hirsh, 1999 Receptor-mediated endocytosis in the Caenorhabditis elegans oocyte. Mol Biol Cell 10**:** 4311-4326.

Green, R. A., H. L. Kao, A. Audhya, S. Arur, J. R. Mayers *et al.*, 2011 A high-resolution C. elegans essential gene network based on phenotypic profiling of a complex tissue. Cell 145**:** 470-482.

Hagstrom, K. A., V. F. Holmes, N. R. Cozzarelli and B. J. Meyer, 2002 C. elegans condensin promotes mitotic chromosome architecture, centromere organization, and sister chromatid segregation during mitosis and meiosis. Genes Dev 16**:** 729-742.

Hamil, K. G., Q. Liu, P. Sivashanmugam, S. Yenugu, R. Soundararajan *et al.*, 2002 Cystatin 11: a new member of the cystatin type 2 family. Endocrinology 143**:** 2787-2796.

Han, M., and P. W. Sternberg, 1990 let-60, a gene that specifies cell fates during C. elegans vulval induction, encodes a ras protein. Cell 63**:** 921-931.

Hannak, E., K. Oegema, M. Kirkham, P. Gonczy, B. Habermann *et al.*, 2002 The kinetically dominant assembly pathway for centrosomal asters in Caenorhabditis elegans is gamma-tubulin dependent. J Cell Biol 157**:** 591-602.

Hwang, H. Y., and H. R. Horvitz, 2002 The Caenorhabditis elegans vulval morphogenesis gene sqv-4 encodes a UDP-glucose dehydrogenase that is temporally and spatially regulated. Proc Natl Acad Sci U S A 99**:** 14224-14229.

Hwang, H. Y., S. K. Olson, J. R. Brown, J. D. Esko and H. R. Horvitz, 2003 The Caenorhabditis elegans genes sqv-2 and sqv-6, which are required for vulval morphogenesis, encode glycosaminoglycan galactosyltransferase II and xylosyltransferase. J Biol Chem 278**:** 11735-11738.

Hyenne, V., A. Apaydin, D. Rodriguez, C. Spiegelhalter, S. Hoff-Yoessle *et al.*, 2015 RAL-1 controls multivesicular body biogenesis and exosome secretion. J Cell Biol 211**:** 27-37.

Jantsch-Plunger, V., and M. Glotzer, 1999 Depletion of syntaxins in the early Caenorhabditis elegans embryo reveals a role for membrane fusion events in cytokinesis. Curr Biol 9**:** 738-745.

Jantsch-Plunger, V., P. Gonczy, A. Romano, H. Schnabel, D. Hamill *et al.*, 2000 CYK-4: A Rho family gtpase activating protein (GAP) required for central spindle formation and cytokinesis. J Cell Biol 149**:** 1391-1404.

Ji, Y. J., K. Y. Choi, H. O. Song, B. J. Park, J. R. Yu *et al.*, 2006 VHA-8, the E subunit of V-ATPase, is essential for pH homeostasis and larval development in C. elegans. FEBS Lett 580**:** 3161-3166.

Jiang, W., Y. Wei, Y. Long, A. Owen, B. Wang *et al.*, 2018 A genetic program mediates cold-warming response and promotes stress-induced phenoptosis in C. elegans. Elife 7.

Jiu, Y., K. Hasygar, L. Tang, Y. Liu, C. I. Holmberg *et al.*, 2014 par-1, atypical pkc, and PP2A/B55 sur-6 are implicated in the regulation of exocyst-mediated membrane trafficking in Caenorhabditis elegans. G3 (Bethesda) 4**:** 173-183.

Johnston, W. L., A. Krizus and J. W. Dennis, 2006 The eggshell is required for meiotic fidelity, polar-body extrusion and polarization of the C. elegans embryo. BMC Biol 4**:** 35.

Joseph-Strauss, D., M. Gorjanacz, R. Santarella-Mellwig, E. Voronina, A. Audhya *et al.*, 2012 Sm protein down-regulation leads to defects in nuclear pore complex disassembly and distribution in C. elegans embryos. Dev Biol 365**:** 445-457.

Kabat, J. L., S. Barberan-Soler and A. M. Zahler, 2009 HRP-2, the Caenorhabditis elegans homolog of mammalian heterogeneous nuclear ribonucleoproteins Q and R, is an alternative splicing factor that binds to UCUAUC splicing regulatory elements. J Biol Chem 284**:** 28490-28497.

Kamikura, D. M., and J. A. Cooper, 2003 Lipoprotein receptors and a disabled family cytoplasmic adaptor protein regulate EGL-17/FGF export in C. elegans. Genes Dev 17**:** 2798-2811.

Kemp, C. A., M. H. Song, M. K. Addepalli, G. Hunter and K. O'Connell, 2007 Suppressors of zyg-1 define regulators of centrosome duplication and nuclear association in Caenorhabditis elegans. Genetics 176**:** 95-113.

Kim, W., R. S. Underwood, I. Greenwald and D. D. Shaye, 2018 OrthoList 2: A New Comparative Genomic Analysis of Human and Caenorhabditis elegans Genes. Genetics 210**:** 445-461.

Kinnaird, J. H., K. Maitland, G. A. Walker, I. Wheatley, F. J. Thompson *et al.*, 2004 HRP-2, a heterogeneous nuclear ribonucleoprotein, is essential for embryogenesis and oogenesis in Caenorhabditis elegans. Exp Cell Res 298**:** 418-430.

Kipreos, E. T., 2005 Ubiquitin-mediated pathways in C. elegans. WormBook**:** 1-24.

Kipreos, E. T., L. E. Lander, J. P. Wing, W. W. He and E. M. Hedgecock, 1996 cul-1 is required for cell cycle exit in C. elegans and identifies a novel gene family. Cell 85**:** 829-839.

Kitagawa, R., E. Law, L. Tang and A. M. Rose, 2002 The Cdc20 homolog, FZY-1, and its interacting protein, IFY-1, are required for proper chromosome segregation in Caenorhabditis elegans. Curr Biol 12**:** 2118-2123.

Kodama, Y., J. H. Rothman, A. Sugimoto and M. Yamamoto, 2002 The stem-loop binding protein CDL-1 is required for chromosome condensation, progression of cell death and morphogenesis in Caenorhabditis elegans. Development 129**:** 187-196.

Kuroyanagi, H., T. Kimura, K. Wada, N. Hisamoto, K. Matsumoto *et al.*, 2000 SPK-1, a C. elegans SR protein kinase homologue, is essential for embryogenesis and required for germline development. Mech Dev 99**:** 51-64.

Labrousse, A. M., M. D. Zappaterra, D. A. Rube and A. M. van der Bliek, 1999 C. elegans dynamin-related protein DRP-1 controls severing of the mitochondrial outer membrane. Mol Cell 4**:** 815-826.

Lall, S., F. Piano and R. E. Davis, 2005 Caenorhabditis elegans decapping proteins: localization and functional analysis of Dcp1, Dcp2, and DcpS during embryogenesis. Mol Biol Cell 16**:** 5880-5890.

Lamitina, S. T., and S. W. L'Hernault, 2002 Dominant mutations in the Caenorhabditis elegans Myt1 ortholog wee-1.3 reveal a novel domain that controls M-phase entry during spermatogenesis. Development 129**:** 5009-5018.

Lee, M. H., and T. Schedl, 2001 Identification of in vivo mRNA targets of GLD-1, a maxi-KH motif containing protein required for C. elegans germ cell development. Genes Dev 15**:** 2408-2420.

Lee, S. K., W. Li, S. E. Ryu, T. Rhim and J. Ahnn, 2010 Vacuolar (H+)-ATPases in Caenorhabditis elegans: what can we learn about giant H+ pumps from tiny worms? Biochim Biophys Acta 1797**:** 1687-1695.

Leidel, S., M. Delattre, L. Cerutti, K. Baumer and P. Gonczy, 2005 SAS-6 defines a protein family required for centrosome duplication in C. elegans and in human cells. Nat Cell Biol 7**:** 115-125.

Leroux, M. R., and E. P. Candido, 1995 Characterization of four new tcp-1-related cct genes from the nematode Caenorhabditis elegans. DNA Cell Biol 14**:** 951-960.

Lieb, J. D., E. E. Capowski, P. Meneely and B. J. Meyer, 1996 DPY-26, a link between dosage compensation and meiotic chromosome segregation in the nematode. Science 274**:** 1732-1736.

Liu, J., T. Rolef Ben-Shahar, D. Riemer, M. Treinin, P. Spann *et al.*, 2000 Essential roles for Caenorhabditis elegans lamin gene in nuclear organization, cell cycle progression, and spatial organization of nuclear pore complexes. Mol Biol Cell 11**:** 3937-3947.

Lundin, V. F., M. Srayko, A. A. Hyman and M. R. Leroux, 2008 Efficient chaperone-mediated tubulin biogenesis is essential for cell division and cell migration in C. elegans. Dev Biol 313**:** 320-334.

Maciejowski, J., J. H. Ahn, P. G. Cipriani, D. J. Killian, A. L. Chaudhary *et al.*, 2005 Autosomal genes of autosomal/X-linked duplicated gene pairs and germ-line proliferation in Caenorhabditis elegans. Genetics 169**:** 1997-2011.

Malone, C. J., W. D. Fixsen, H. R. Horvitz and M. Han, 1999 UNC-84 localizes to the nuclear envelope and is required for nuclear migration and anchoring during C. elegans development. Development 126**:** 3171-3181.

Malone, C. J., L. Misner, N. Le Bot, M. C. Tsai, J. M. Campbell *et al.*, 2003 The C. elegans hook protein, ZYG-12, mediates the essential attachment between the centrosome and nucleus. Cell 115**:** 825-836.

Michelle, C., P. Vourc'h, L. Mignon and C. R. Andres, 2009 What was the set of ubiquitin and ubiquitin-like conjugating enzymes in the eukaryote common ancestor? J Mol Evol 68**:** 616-628.

Mishima, M., S. Kaitna and M. Glotzer, 2002 Central spindle assembly and cytokinesis require a kinesin-like protein/RhoGAP complex with microtubule bundling activity. Dev Cell 2**:** 41-54.

Mizuguchi, S., T. Uyama, H. Kitagawa, K. H. Nomura, K. Dejima *et al.*, 2003 Chondroitin proteoglycans are involved in cell division of Caenorhabditis elegans. Nature 423**:** 443-448.

Morita, K., K. Hirono and M. Han, 2005 The Caenorhabditis elegans ect-2 RhoGEF gene regulates cytokinesis and migration of epidermal P cells. EMBO Rep 6**:** 1163-1168.

Motegi, F., and A. Sugimoto, 2006 Sequential functioning of the ECT-2 RhoGEF, RHO-1 and CDC-42 establishes cell polarity in Caenorhabditis elegans embryos. Nat Cell Biol 8**:** 978-985.

Nakazawa, Y., M. Hiraki, R. Kamiya and M. Hirono, 2007 SAS-6 is a cartwheel protein that establishes the 9-fold symmetry of the centriole. Curr Biol 17**:** 2169-2174.

Nillegoda, N. B., J. Kirstein, A. Szlachcic, M. Berynskyy, A. Stank *et al.*, 2015 Crucial HSP70 co-chaperone complex unlocks metazoan protein disaggregation. Nature 524**:** 247-251.

Nousch, M., N. Techritz, D. Hampel, S. Millonigg and C. R. Eckmann, 2013 The Ccr4-Not deadenylase complex constitutes the main poly(A) removal activity in C. elegans. J Cell Sci 126**:** 4274-4285.

O'Rourke, S. M., M. D. Dorfman, J. C. Carter and B. Bowerman, 2007 Dynein modifiers in C. elegans: light chains suppress conditional heavy chain mutants. PLoS Genet 3**:** e128.

Oegema, K., A. Desai, S. Rybina, M. Kirkham and A. A. Hyman, 2001 Functional analysis of kinetochore assembly in Caenorhabditis elegans. J Cell Biol 153**:** 1209-1226.

Ofulue, E. N., and E. P. Candido, 1991 Molecular cloning and characterization of the Caenorhabditis elegans elongation factor 2 gene (eft-2). DNA Cell Biol 10**:** 603-611.

Ogawa, S., Y. Matsubayashi and E. Nishida, 2004 An evolutionarily conserved gene required for proper microtubule architecture in Caenorhabditis elegans. Genes Cells 9**:** 83-93.

Oka, T., and M. Futai, 2000 Requirement of V-ATPase for ovulation and embryogenesis in Caenorhabditis elegans. J Biol Chem 275**:** 29556-29561.

Oka, T., R. Yamamoto and M. Futai, 1997 Three vha genes encode proteolipids of Caenorhabditis elegans vacuolar-type ATPase. Gene structures and preferential expression in an H-shaped excretory cell and rectal cells. J Biol Chem 272**:** 24387-24392.

Olson, S. K., G. Greenan, A. Desai, T. Muller-Reichert and K. Oegema, 2012 Hierarchical assembly of the eggshell and permeability barrier in C. elegans. J Cell Biol 198**:** 731-748.

Ono, T., A. Losada, M. Hirano, M. P. Myers, A. F. Neuwald *et al.*, 2003 Differential contributions of condensin I and condensin II to mitotic chromosome architecture in vertebrate cells. Cell 115**:** 109-121.

Ooi, S. L., J. R. Priess and S. Henikoff, 2006 Histone H3.3 variant dynamics in the germline of Caenorhabditis elegans. PLoS Genet 2**:** e97.

Pasierbek, P., M. Fodermayr, V. Jantsch, M. Jantsch, D. Schweizer *et al.*, 2003 The Caenorhabditis elegans SCC-3 homologue is required for meiotic synapsis and for proper chromosome disjunction in mitosis and meiosis. Exp Cell Res 289**:** 245-255.

Pellettieri, J., V. Reinke, S. K. Kim and G. Seydoux, 2003 Coordinate activation of maternal protein degradation during the egg-to-embryo transition in C. elegans. Dev Cell 5**:** 451-462.

Pena, S., T. Sherman, P. S. Brookes and K. Nehrke, 2016 The Mitochondrial Unfolded Protein Response Protects against Anoxia in Caenorhabditis elegans. PLoS One 11**:** e0159989.

Phillips, J. B., R. Lyczak, G. C. Ellis and B. Bowerman, 2004 Roles for two partially redundant alpha-tubulins during mitosis in early Caenorhabditis elegans embryos. Cell Motil Cytoskeleton 58**:** 112-126.

Piano, F., A. J. Schetter, D. G. Morton, K. C. Gunsalus, V. Reinke *et al.*, 2002 Gene clustering based on RNAi phenotypes of ovary-enriched genes in C. elegans. Curr Biol 12**:** 1959-1964.

Pintard, L., T. Kurz, S. Glaser, J. H. Willis, M. Peter *et al.*, 2003a Neddylation and deneddylation of CUL-3 is required to target MEI-1/Katanin for degradation at the meiosis-to-mitosis transition in C. elegans. Curr Biol 13**:** 911-921.

Pintard, L., J. H. Willis, A. Willems, J. L. Johnson, M. Srayko *et al.*, 2003b The BTB protein MEL-26 is a substrate-specific adaptor of the CUL-3 ubiquitin-ligase. Nature 425**:** 311-316.

Polinko, E. S., and S. Strome, 2000 Depletion of a Cks homolog in C. elegans embryos uncovers a post-metaphase role in both meiosis and mitosis. Curr Biol 10**:** 1471-1474.

Poteryaev, D., H. Fares, B. Bowerman and A. Spang, 2007 Caenorhabditis elegans SAND-1 is essential for RAB-7 function in endosomal traffic. EMBO J 26**:** 301-312.

Powers, J., D. J. Rose, A. Saunders, S. Dunkelbarger, S. Strome *et al.*, 2004 Loss of KLP-19 polar ejection force causes misorientation and missegregation of holocentric chromosomes. J Cell Biol 166**:** 991-1001.

Pujol, N., C. Bonnerot, J. J. Ewbank, Y. Kohara and D. Thierry-Mieg, 2001 The Caenorhabditis elegans unc-32 gene encodes alternative forms of a vacuolar ATPase a subunit. J Biol Chem 276**:** 11913-11921.

Puoti, A., and J. Kimble, 2000 The hermaphrodite sperm/oocyte switch requires the Caenorhabditis elegans homologs of PRP2 and PRP22. Proc Natl Acad Sci U S A 97**:** 3276-3281.

Rahman, M. M., S. Rosu, D. Joseph-Strauss and O. Cohen-Fix, 2014 Down-regulation of tricarboxylic acid (TCA) cycle genes blocks progression through the first mitotic division in Caenorhabditis elegans embryos. Proc Natl Acad Sci U S A 111: 2602-2607.

Raich, W. B., C. Moorman, C. O. Lacefield, J. Lehrer, D. Bartsch *et al.*, 2003 Characterization of Caenorhabditis elegans homologs of the Down syndrome candidate gene DYRK1A. Genetics 163: 571-580.

Raich, W. B., A. N. Moran, J. H. Rothman and J. Hardin, 1998 Cytokinesis and midzone microtubule organization in Caenorhabditis elegans require the kinesin-like protein ZEN-4. Mol Biol Cell 9: 2037-2049.

Rastogi, S., B. Borgo, N. Pazdernik, P. Fox, E. R. Mardis *et al.*, 2015 Caenorhabditis elegans glp-4 Encodes a Valyl Aminoacyl tRNA Synthetase. G3 (Bethesda) 5: 2719-2728.

Rhoads, R. E., T. D. Dinkova and N. L. Korneeva, 2006 Mechanism and regulation of translation in C. elegans. WormBook: 1-18.

Riemer, D., H. Dodemont and K. Weber, 1993 A nuclear lamin of the nematode Caenorhabditis elegans with unusual structural features; cDNA cloning and gene organization. Eur J Cell Biol 62: 214-223.

Roberts, S. B., S. W. Emmons and G. Childs, 1989 Nucleotide sequences of Caenorhabditis elegans core histone genes. Genes for different histone classes share common flanking sequence elements. J Mol Biol 206: 567-577.

Rogala, K. B., N. J. Dynes, G. N. Hatzopoulos, J. Yan, S. K. Pong *et al.*, 2015 The Caenorhabditis elegans protein SAS-5 forms large oligomeric assemblies critical for centriole formation. Elife 4: e07410.

Sasagawa, Y., T. Urano, Y. Kohara, H. Takahashi and A. Higashitani, 2003 Caenorhabditis elegans RBX1 is essential for meiosis, mitotic chromosomal condensation and segregation, and cytokinesis. Genes Cells 8: 857-872.

Sato, M., K. Saegusa, K. Sato, T. Hara, A. Harada *et al.*, 2011 Caenorhabditis elegans SNAP-29 is required for organellar integrity of the endomembrane system and general exocytosis in intestinal epithelial cells. Mol Biol Cell 22: 2579-2587.

Schlesinger, A., C. A. Shelton, J. N. Maloof, M. Meneghini and B. Bowerman, 1999 Wnt pathway components orient a mitotic spindle in the early Caenorhabditis elegans embryo without requiring gene transcription in the responding cell. Genes Dev 13: 2028-2038.

Schmutz, C., and A. Spang, 2005 Knockdown of the centrosomal component SAS-5 results in defects in nuclear morphology in Caenorhabditis elegans. Eur J Cell Biol 84: 75-82.

Schumacher, J. M., N. Ashcroft, P. J. Donovan and A. Golden, 1998 A highly conserved centrosomal kinase, AIR-1, is required for accurate cell cycle progression and segregation of developmental factors in Caenorhabditis elegans embryos. Development 125: 4391-4402.

Segbert, C., R. Barkus, J. Powers, S. Strome, W. M. Saxton *et al.*, 2003 KLP-18, a Klp2 kinesin, is required for assembly of acentrosomal meiotic spindles in Caenorhabditis elegans. Mol Biol Cell 14: 4458-4469.

Severson, A. F., D. L. Baillie and B. Bowerman, 2002 A Formin Homology protein and a profilin are required for cytokinesis and Arp2/3-independent assembly of cortical microfilaments in C. elegans. Curr Biol 12: 2066-2075.

Shaye, D. D., and I. Greenwald, 2011 OrthoList: a compendium of C. elegans genes with human orthologs. PLoS One 6**:** e20085.
Siddiqui, S. S., 2002 Metazoan motor models: kinesin superfamily in C. elegans. Traffic 3**:** 20-28.
Sieburth, D. S., M. Sundaram, R. M. Howard and M. Han, 1999 A PP2A regulatory subunit positively regulates Ras-mediated signaling during Caenorhabditis elegans vulval induction. Genes Dev 13**:** 2562-2569.
Skop, A. R., and J. G. White, 1998 The dynactin complex is required for cleavage plane specification in early Caenorhabditis elegans embryos. Curr Biol 8**:** 1110-1116.
Song, M. H., Y. Liu, D. E. Anderson, W. J. Jahng and K. F. O'Connell, 2011 Protein phosphatase 2A-SUR-6/B55 regulates centriole duplication in C. elegans by controlling the levels of centriole assembly factors. Dev Cell 20**:** 563-571.
Sonneville, R., and P. Gonczy, 2004 Zyg-11 and cul-2 regulate progression through meiosis II and polarity establishment in C. elegans. Development 131**:** 3527-3543.
Sonnichsen, B., L. B. Koski, A. Walsh, P. Marschall, B. Neumann *et al.*, 2005 Full-genome RNAi profiling of early embryogenesis in Caenorhabditis elegans. Nature 434**:** 462-469.
Srayko, M., D. W. Buster, O. A. Bazirgan, F. J. McNally and P. E. Mains, 2000 MEI-1/MEI-2 katanin-like microtubule severing activity is required for Caenorhabditis elegans meiosis. Genes Dev 14**:** 1072-1084.
Srinivasan, D. G., R. M. Fisk, H. Xu and S. van den Heuvel, 2003 A complex of LIN-5 and GPR proteins regulates G protein signaling and spindle function in C elegans. Genes Dev 17**:** 1225-1239.
Stanvitch, G., and L. L. Moore, 2008 cin-4, a gene with homology to topoisomerase II, is required for centromere resolution by cohesin removal from sister kinetochores during mitosis. Genetics 178**:** 83-97.
Stear, J. H., and M. B. Roth, 2002 Characterization of HCP-6, a C. elegans protein required to prevent chromosome twisting and merotelic attachment. Genes Dev 16**:** 1498-1508.
Terasawa, M., M. Toya, F. Motegi, M. Mana, K. Nakamura *et al.*, 2010 Caenorhabditis elegans ortholog of the p24/p22 subunit, DNC-3, is essential for the formation of the dynactin complex by bridging DNC-1/p150(Glued) and DNC-2/dynamitin. Genes Cells 15**:** 1145-1157.
van der Voet, H., G. W. van der Heijden, P. M. Bos, S. Bosgra, P. E. Boon *et al.*, 2009 A model for probabilistic health impact assessment of exposure to food chemicals. Food Chem Toxicol 47**:** 2926-2940.
Velarde, N., K. C. Gunsalus and F. Piano, 2007 Diverse roles of actin in C. elegans early embryogenesis. BMC Dev Biol 7**:** 142.
Walston, T., C. Tuskey, L. Edgar, N. Hawkins, G. Ellis *et al.*, 2004 Multiple Wnt signaling pathways converge to orient the mitotic spindle in early C. elegans embryos. Dev Cell 7**:** 831-841.
Wang, F., J. Yoder, I. Antoshechkin and M. Han, 2003 Caenorhabditis elegans EVL-14/PDS-5 and SCC-3 are essential for sister chromatid cohesion in meiosis and mitosis. Mol Cell Biol 23**:** 7698-7707.

Waters, K., A. Z. Yang and V. Reinke, 2010 Genome-wide analysis of germ cell proliferation in C.elegans identifies VRK-1 as a key regulator of CEP-1/p53. Dev Biol 344**:** 1011-1025.

Wignall, S. M., and A. M. Villeneuve, 2009 Lateral microtubule bundles promote chromosome alignment during acentrosomal oocyte meiosis. Nat Cell Biol 11**:** 839-844.

Wolff, I. D., M. V. Tran, T. J. Mullen, A. M. Villeneuve and S. M. Wignall, 2016 Assembly of Caenorhabditis elegans acentrosomal spindles occurs without evident microtubule-organizing centers and requires microtubule sorting by KLP-18/kinesin-12 and MESP-1. Mol Biol Cell 27**:** 3122-3131.

Wright, A. J., and C. P. Hunter, 2003 Mutations in a beta-tubulin disrupt spindle orientation and microtubule dynamics in the early Caenorhabditis elegans embryo. Mol Biol Cell 14**:** 4512-4525.

Yamamoto, T. G., S. Watanabe, A. Essex and R. Kitagawa, 2008 SPDL-1 functions as a kinetochore receptor for MDF-1 in Caenorhabditis elegans. J Cell Biol 183**:** 187-194.

Yoder, J. H., and M. Han, 2001 Cytoplasmic dynein light intermediate chain is required for discrete aspects of mitosis in Caenorhabditis elegans. Mol Biol Cell 12**:** 2921-2933.

Zhao, Z., L. L. Fang, R. Johnsen and D. L. Baillie, 2004 ATP-binding cassette protein E is involved in gene transcription and translation in Caenorhabditis elegans. Biochem Biophys Res Commun 323**:** 104-111.

Zipperlen, P., A. G. Fraser, R. S. Kamath, M. Martinez-Campos and J. Ahringer, 2001 Roles for 147 embryonic lethal genes on C.elegans chromosome I identified by RNA interference and video microscopy. EMBO J 20**:** 3984-3992.
